# SYMBIOSIS: Synthetic Manipulable Biobricks via Orthogonal Serine Integrase Systems

**DOI:** 10.1101/2021.11.03.467214

**Authors:** Fang Ba, Yushi Liu, Wan-Qiu Liu, Xintong Tian, Jian Li

## Abstract

Serine integrases are emerging as one of the most powerful biological tools for synthetic biology. They have been widely used across genome engineering and genetic circuit design. However, developing serine integrase-based tools for directly/precisely manipulating synthetic biobricks is still missing. Here, we report SYMBIOSIS, a versatile method that can robustly manipulate DNA parts *in vivo* and *in vitro*. First, we proposed a “Keys match Locks” model to demonstrate that three orthogonal serine integrases are able to irreversibly and stably switch on seven synthetic biobricks with high accuracy *in vivo*. Then, we demonstrated that purified integrases can facilitate the assembly of “Donor” and “Acceptor” plasmids *in vitro* to construct composite plasmids. Finally, we used SYMBIOSIS to assemble different chromoprotein genes and create novel colored *Escherichia coli*. We anticipate that our SYMBIOSIS strategy will accelerate synthetic biobricks manipulation, genetic circuit design, and multiple plasmids assembly for synthetic biology with broad potential applications.

## Introduction

DNA is essential for cellular processes and behaviors as the instructions for biological processes are encoded in DNA molecules. Changing DNA sequences to reprogram cells can endow them with new, specialized properties. Over the last two decades, synthetic biology, taking the core mission to (re)design and construct new biological parts, devices, and systems, has been rapidly developed to require even more controllable and efficient tools for DNA manipulation and modification^1–4^. To meet this demand, synthetic biologists have paid much more attention to DNA-modifying enzymes, which can catalyze DNA sequence variations (e.g., deletion, insertion, and mutation) for precise and predictable programming of living organisms^5^.

Recombinase is a class of enzymes that catalyzes DNA-DNA site-specific recombination events within single or among different DNA strands^6^. Serine integrase, a subfamily of recombinase, can recognize and catalyze recombination between two specific DNA sites (approximately 50 bp for each) called *attP* (attachment site by Phage) and *attB* (attachment site by Bacteria)^7–11^. Depending on the *attP*/*attB* position and orientation, the DNA fragments can be deleted, integrated, recombined, or inverted^7^. The features of site-specific, orthogonality, irreversibility, high affinity, and high efficiency make serine integrase a powerful genetic tool in synthetic biology^7^. For example, serine integrase has been widely used for linear DNA fragment assembly^12^, genome engineering^13^, and genetic circuit design^14–20^. In particular, a series of complicated genetic circuits were constructed recently using serine integrase technologies such as Boolean logic gates^14–19^, memory circuits^14–20^, state machine^16^, binary counter^17^, message storage^16, 19^, and gene expression cascades^20^. However, the “input” and “output” signals within one genetic circuit often only have a one-to-one corresponding relationship, which means that a finite number of orthogonal serine integrases can only switch on an equal number of genetic circuits; therefore, the output signals are limited to the number of serine integrases present^16^. Moreover, the common problems of inducible recombinase expression leakage and cell toxicity *in vivo* still remain unsolved^17, 18, 21^.

In synthetic biology, the general strategy for biobrick construction and assembly often uses one of the following methods: Restriction Enzyme Digestion and Ligation^22^, TOPO Cloning^23, 24^, Golden Gate Assembly^25^, Gateway Cloning^26^, and Gibson Assembly^27^. However, these approaches typically require the preparation of multiple linear DNA fragments in advance by Polymerase Chain Reaction (PCR), which is time-consuming and labor-intensive. Therefore, a simple, precise, and efficient method is highly desirable for the one-step assembly of circular brobricks (generally considered as plasmids). It is worth noting that a tyrosine recombinase (another subfamily of recombinase) based *in vitro* DNA assembly method “ACEMBL” has been established by using Cre-LoxP recombination^28^. Cre recombinase, one of the tyrosine recombinases, cleaves single DNA strands forming covalent 3′-phosphotyrosine bonds with the DNA backbone and rejoins the strands via a Holliday junction-like intermediate state^6^. However, if one plasmid contains more than one homologous LoxP site, it can form many unpredictable variants before the next round of assembly^28, 29^. The exact DNA sequences of all possible variants are subsequently subjected to post-verification and selection by restriction enzyme digestions or DNA sequencing, which limits precise, predictable, and efficient manipulation of plasmid assembly. Moreover, unlike serine integrases, the tyrosine recombinase Cre mediated Cre-LoxP reaction is reversible, often leading to unpredictable variation and reversibly degradation^6^. By contrast, serine integrases make double strand breaks in DNA forming covalent 5′-phosphoserine bonds with the DNA backbone before religation^6^. Notably, serine integrase-driven DNA recombination is highly directional, orthogonal, and irreversible (when no presence of an accessory recombination directionality factor, RDF), which makes the system a reliable choice for catalyzing *in vitro*DNA-DNA recombination^7^.

In this study, we introduce SYMBIOSIS (<u>Sy</u>nthetic <u>M</u>anipulable <u>Bi</u>obricks via <u>O</u>rthogonal <u>S</u>erine Integrase <u>S</u>ystems), aiming to provide a strategy that can precisely switch on scalable genetic circuits in *Escherichia coli* and expand the toolkit to establish a method for *in vitro* multiple plasmids assembly. The foundational principle is that we can manipulate DNA parts through serine integrases-driven DNA recombination. To demonstrate the feasibility of SYMBIOSIS *in vivo*, we initially constructed three plasmids with constitutive promoters and each of them harbored an orthogonal serine integrase TP901-1, Bxb1, or Phic31, respectively. We defined each above plasmid as an “input” signal once it was transformed into *E. coli* cells. Meanwhile, seven different chromoproteins were chosen as “output” signals. By observing whether the changes in the color of *E. coli* cells, we then can determine the “on/off” states of synthetic biobricks. Next, we demonstrated that purified serine integrases were able to assemble “Donor” and “Acceptor” plasmids *in vitro*, generating expression vectors for protein synthesis *in vivo*. More interestingly, we applied our SYMBIOSIS *in vitro* to construct various plasmids harboring different combinations of chromoprotein genes. When transformed, these genes created some unique colored *E. coli* cells that, to our knowledge, have never been reported so far.

## Results

### Establishing the SYMBIOSIS framework

The idea for designing SYMBIOSIS was inspired by the mathematical concept of “set”. The elements that constitute sets can be any object. In principle, a finite number of elements can be freely combined and organized as various different non-empty subsets. If there is one element *A*, it can only form one non-empty subset {*A*}. When another element *B* is added, two new non- empty subsets {*B*} and {*A*, *B*} can be formed. Similarly, three elements *A*, *B*, and *C* can generate seven non-empty subsets {*A*}, {*B*}, {*C*}, {*A*, *B*}, {*A*, *C*}, {*B*, *C*}, and {*A*, *B*, *C*}. By parity of reasoning, the number of *n* elements can be composed into 2^n^-1 different non-empty subsets^30^ (**Fig. 1a**).

**Fig. 1.**
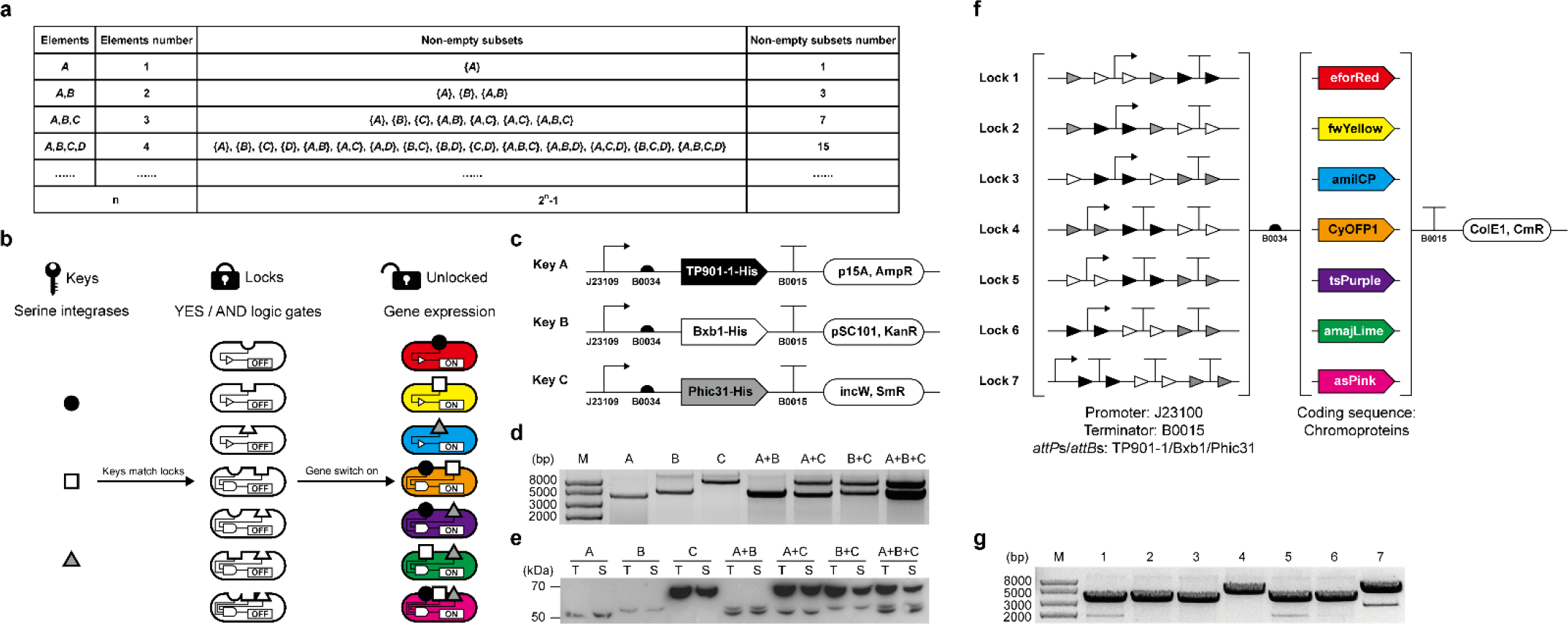
Design and validation of the SYMBIOSIS framework. (**a**) Schematic of the mathematical “set” concept. Non-empty subsets are organized with elements. The number of non-empty subsets increases exponentially with the element number (*n*) as 2^n^-1. (**b**) Design of the “Keys match Locks (KmL)” model. Three orthogonal serine integrases (TP901-1: black circle, Bxb1: white square, and Phic31: gray triangle) work as “Keys” to match *E. coli* strains (ovals with gaps) containing seven different “Locks” genetic circuits. A correct “Key-Lock” match can unlock the closed circuit and switch on chromoprotein expression. (**c**) Architectures of three “Keys” plasmids. TP901-1, Bxb1, and Phic31 are constructed in three compatible plasmids, respectively, with a weak constitutive promoter J23109 (pFB1 to pFB3, see **Supplementary Table 4**). (**d**) Compatibility of single, double or triple “Keys” plasmids in one *E. coli* strain. Extracted plasmids are digested by XbaI and then separated by agarose gel electrophoresis. (**e**) Western Blot analysis of individual expression and coexpression of TP901-1 (56.5 kDa), Bxb1 (57.2 kDa), and Phic31 (67.9 kDa) labeled with the anti-His antibody. T, total protein; S, soluble protein. (**f**) Schematic of seven “Locks” plasmids. “Lock” BioBricks (left parenthesis) are assembled with a strong promoter J23100, a strong terminator B0015, and *attP*-*attB* sites (TP901-1 *attP*/*attB*: black triangles, Bxb1 *attP*/*attB*: white triangles, and Phic31 *attP*/*attB*: gray triangles). Seven different chromoproteins (eforRed, fwYellow, amilCP, CyOFP1, tsPurple, amajLime, and asPink, right parenthesis) work as output signals. The seven “Locks” plasmids are constructed using the same vector that is compatible to three “Keys” plasmids. (**g**) Plasmid sizes of the seven “Locks” (pFB20 to pFB26, see **Supplementary Table 5**) after XbaI digestion. Expected bands of the seven digested plasmids are observed at 3371, 3401, 3356, 3521, 3506, 3509, and 3647 bp, respectively.

Based on the principle of set, we were motivated to establish the SYMBIOSIS framework by matching biological inputs (elements) with visible outputs (subsets) in living *E. coli* cells. To this end, we first chose three orthogonal serine integrases TP901-1^8, 9^, Bxb1^10^, and Phic31^11^, which were defined as input elements *A*, *B*, and *C*, respectively. Then, seven chromoproteins were used as output signals in response to three serine integrase inputs. We called this input-output relation as a “Keys match Locks (KmL)” model (**Fig. 1b**). In the KmL model, only when integrase(s) matches the correct logic gates, the “Locked” genetic circuits can be switched on, leading to the expression of corresponding chromoproteins in *E. coli*.

We began our investigation by trying two expression systems, which are L-arabinose inducible AraC-pBAD expression system^31^ and anhydrotetracycline inducible TetR expression system^32^, to express serine integrases as inputs (“Keys”) together with different “Locks” circuits.

Unfortunately, our initial experiments were not successful due to the leaky expression of integrases that caused unexpected chromoproteins expression (data not shown). We suspected that a basal leakage of integrase was enough to unlock the genetic circuit, resulting in the activation of gene expression. To tackle this problem and achieve the goal of a strict switch on/off using integrases, we decided to construct three compatible and constitutive expression plasmids for three serine integrases (**Fig. 1c**). In this context, we defined these plasmids as “Keys” once they were transformed into *E. coli* cells. We initially hypothesized that a stronger constitutively active promoter may lead to more integrase expression and higher catalytic efficiency. However, we found that the strongest promoter (J23100) resulted in an unexpected larger plasmid size (>5000 bp) than that of the correct plasmid (4007 bp) harboring the TP901-1 gene (pFB7, **Supplementary Table 5**) after approximately one day’s cultivation (**Supplementary Fig. 1**). The exact reason was unclear, but we suspected that overexpression of TP901-1 with the strongest promoter might cause cellular toxicity and therefore plasmid mutation. We then constructed a series of plasmids varying in their promoter strength to express TP901-1 integrase (see pFB7 to pFB11 in **Supplementary Table 5**). The results indicated that, like the strongest promoter (J23100), both the second (J23106) and third (J23105) strong promoters also led to an increased plasmid size on Day 4 and Day 5, respectively (**Supplementary Fig. 1**). By contrast, no plasmid mutations were observed with two low strength promoters (J23114 and J23109) even after five days of serial dilution/cultivation (**Supplementary Fig. 1**). While the above problem did not happen to Bxb1 and Phic31, we finally chose the lowest strength promoter (J23109) for all three plasmids construction to avoid potential plasmid mutations (pFB10, pFB13, and pFB15) (**Fig. 1c**). After transformation, each plasmid could stably exist in *E. coli*, and expressed integrase was active to catalyze the DNA recombination (**Fig. 1d** and **Supplementary Fig. 2**). Notably, two or three compatible plasmids were able to co-exist in *E. coli* for at least six days (**Supplementary Fig. 3**). In addition, these plasmids exhibited no obvious cell growth inhibition on *E. coli* as shown in **Supplementary Fig. 4**. All three integrases were fused with a *C*-terminal 6xHis tag and their expression was analyzed by Western-Blot. As can be seen in **Fig. 1e**, three integrases could be expressed individually or together in *E. coli*, demonstrating the compatibility of three plasmids and their ability for constitutive soluble protein expression.

With three integrase “Keys” in hand, we next constructed seven “Locks” circuits, in which two DNA parts were alterable (**Fig. 1f**). The first part consists of a promoter, terminator(s), and *attP*/*attB* sites in different orders. This part determines the matching rule between “Keys” and “Locks”. Only when proper integrases are present, the terminators between relevant *attP*/*attB* sites will be deleted, leading to the downstream gene expression (**Supplementary Fig. 5**). The second part is inserted with one of the seven protein coding sequences working as “output” signals. For easy observation and discrimination, we chose seven chromoproteins as our outputs: eforRed^33^, fwYellow^34^, amilCP^33^, CyOFP1^35^, tsPurple^34^, amajLime^36^, and asPink^34^. After plasmid construction, the correct sizes of seven “Locks” plasmids were first confirmed by a restriction digestion test (**Fig. 1g**). Since repeated genetic elements (e.g., promoter and terminator) in one plasmid might cause plasmid instability^37^, we then carried out a 6-day serial dilution/cultivation to test the stability of each “Locks” plasmid (pFB20 to pFB26). The results showed that seven “Locks” plasmids were stable with expected DNA fragment sizes across at least 6 days cultivation (**Supplementary Fig. 6**). In addition, seven chromoproteins could be expressed with distinct colors in *E. coli* using normal constitutive expression plasmids (pFB131 to pFB137, **Supplementary Fig. 7**). Therefore, they can be used as indicators for different output signals in our following studies.

### Evaluating the orthogonality of three “Keys” and seven “Locks”

Having successfully constructed the plasmids of three “Keys” and seven “Locks”, we next set out to explore whether “Keys” can manipulate “Locks” orthogonally. To this end, a 7x7 orthogonal table was designed, showing a total of forty-nine possible combinations with “Keys” and “Locks” (**Fig. 2a**). In the table, only seven proper matches along the diagonal line (from top left to bottom right, colored with light green) will result in the deletion of terminators between promoter and RBS and activate the subsequent chromoproteins expression. However, the rest of the mismatches will make incorrect organization of promoter and terminator in front of the RBS and thus lead to the failure expression of chromoproteins. To test the orthogonality, we first transformed seven different combinations of “Keys” plasmids into *E. coli* to create seven distinct *E. coli* strains. Then, seven “Locks” plasmids were individually transformed into above seven *E. coli* strains, resulting in a total of forty-nine *E. coli* strains with all possible combinations of “Keys” and “Locks” (for details, see **Fig. 2b**). For comparison, we cultivated all *E. coli* strains in liquid LB and collected cell pellets in 1.5 mL tubes. Then, the pellets were photographed under bright field and LED transilluminator. The results indicated that our KmL model worked precisely as designed and only correct KmL strains on the diagonal line were able to express chromoproteins (**Fig. 2b**, framed with light green; for the plasmid combinations of seven correct KmL strains, see **Supplementary Fig. 8**). Clearly, colored *E. coli* pellets were visible under bright field and/or LED transilluminator, whereas other mismatched strains were observed as a wild-type *E. coli* color. Thus, our results demonstrated that the KmL system is highly orthogonal to one another. In addition, we were curious to know the recombination efficiency of our KmL system. To do this, seven KmL strains were cultivated overnight on LB-agar plates and then the colored colonies were counted to calculate the recombination efficiency (**Supplementary Fig. 8**). The data suggested that the efficiency of each KmL was remarkably high, ranging from 95% to nearly 100% (**Fig. 2c**). This is in line with previous reports that orthogonal serine integrases can precisely bind their own recognition sites and efficiently catalyze the DNA recombination^7, 16^.

**Fig. 2.**
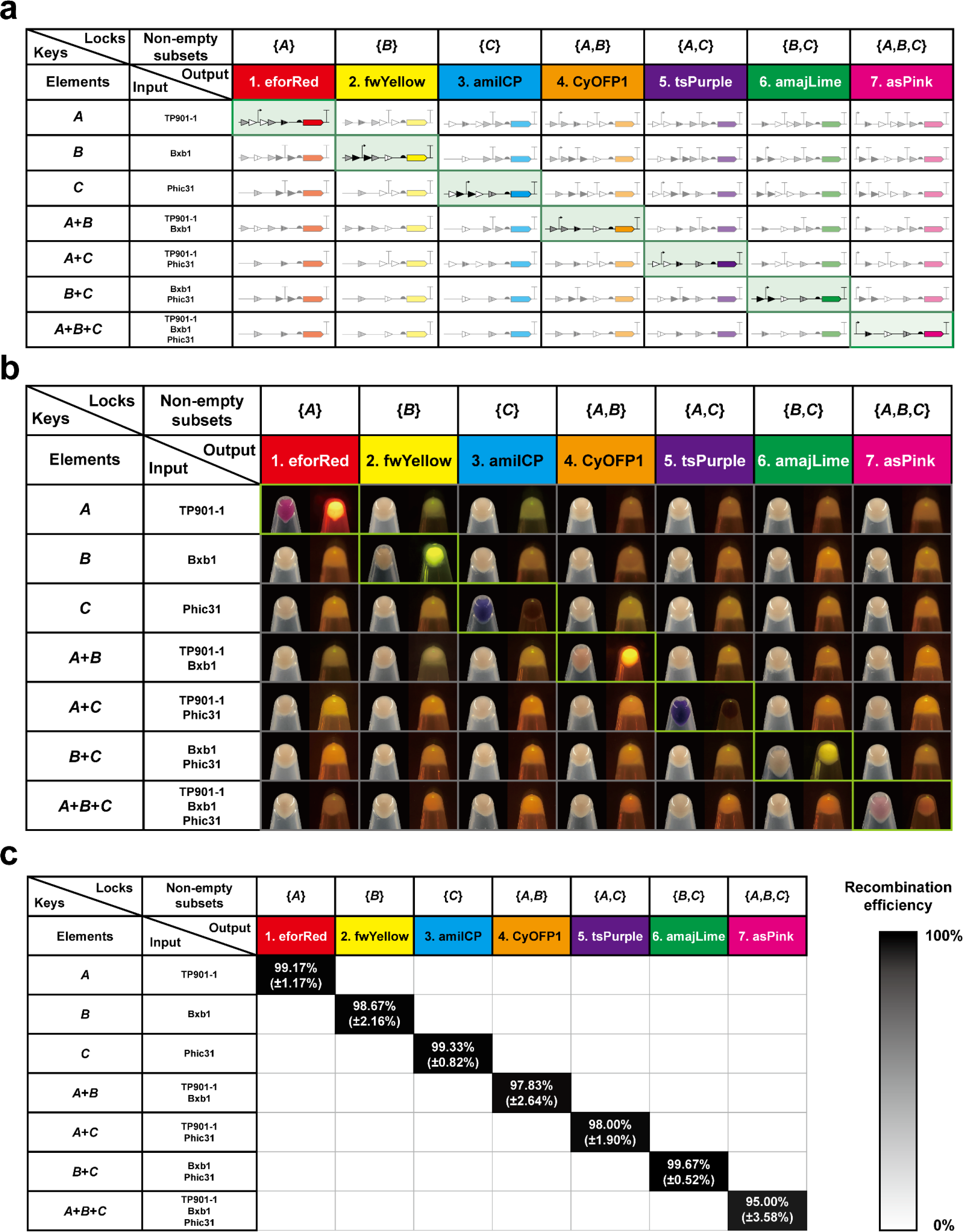
Orthogonality and performance of the “Keys match Locks” model. (**a**) Schematic diagram of the orthogonality matrix. Elements in the first column are input “Keys” (*A*: TP901-1, *B*: Bxb1, and *C*: Phic31). Non-empty subsets in the first line represent seven different “Locks” plasmids containing chromoproteins as outputs. The forty-nine organizations of DNA parts shown in the table indicate the “Keys”- switched “Locks” plasmids. Only seven proper matches on the diagonal line (colored with light green) can activate the expression of chromoproteins. The remaining mismatches make incorrect organization of DNA parts (i.e., promoter and terminator) leading to no protein expression. (**b**) *E. coli* cells are collected and photographed under bright field (left) and LED transilluminator (right). Each *E. coli* sample is collected in a 1.5 mL centrifuge tube from 3 mL overnight culture. Only cell pellets on the diagonal line (framed with light green) with proper Key-Lock matches show chromoprotein colors. All samples are repeated at least three times with similar results. (**c**) Recombination efficiency of the seven proper Key-Lock matches. Each value (mean ± s.d.) is calculated with six biological replicates by counting at least 100 colonies on LB-agar plates.

### Enhancing the SYMBIOSIS system to amplify output signals

While we have demonstrated the success of using our KmL model for SYMBIOSIS *in vivo*, we observed that the colors of the seven “Unlocked” *E. coli* pellets were lower to different levels as compared to their positive controls (pFB131 to pFB137) (**Supplementary Fig. 9**). We hypothesized that the residual *attL*/*attR* sites between the promoter and RBS in the 5’-UTR region might form mRNA secondary structures, which serve as steric blocks for ribosome binding and hinder the downstream protein translation^38, 39^. To test this hypothesis and find out the reason for low protein expression, we next sought to investigate the transcription and translation levels of designed genetic circuits. First, we replaced all chromoprotein coding sequences with the same reporter gene (i.e., superfolder green fluorescent protein, sfGFP) for easy quantification and comparison (**Fig. 3a**, **b**). After cultivation, the fluorescence of each sample was measured. We found that no sfGFP fluorescence was detected from the group of “Locked” genetic circuits (**Fig. 3c** top panel, **3d**), suggesting no sfGFP expression leakage. When each “Locked” circuit was switched on, fluorescent sfGFP was clearly expressed.

**Fig. 3.**
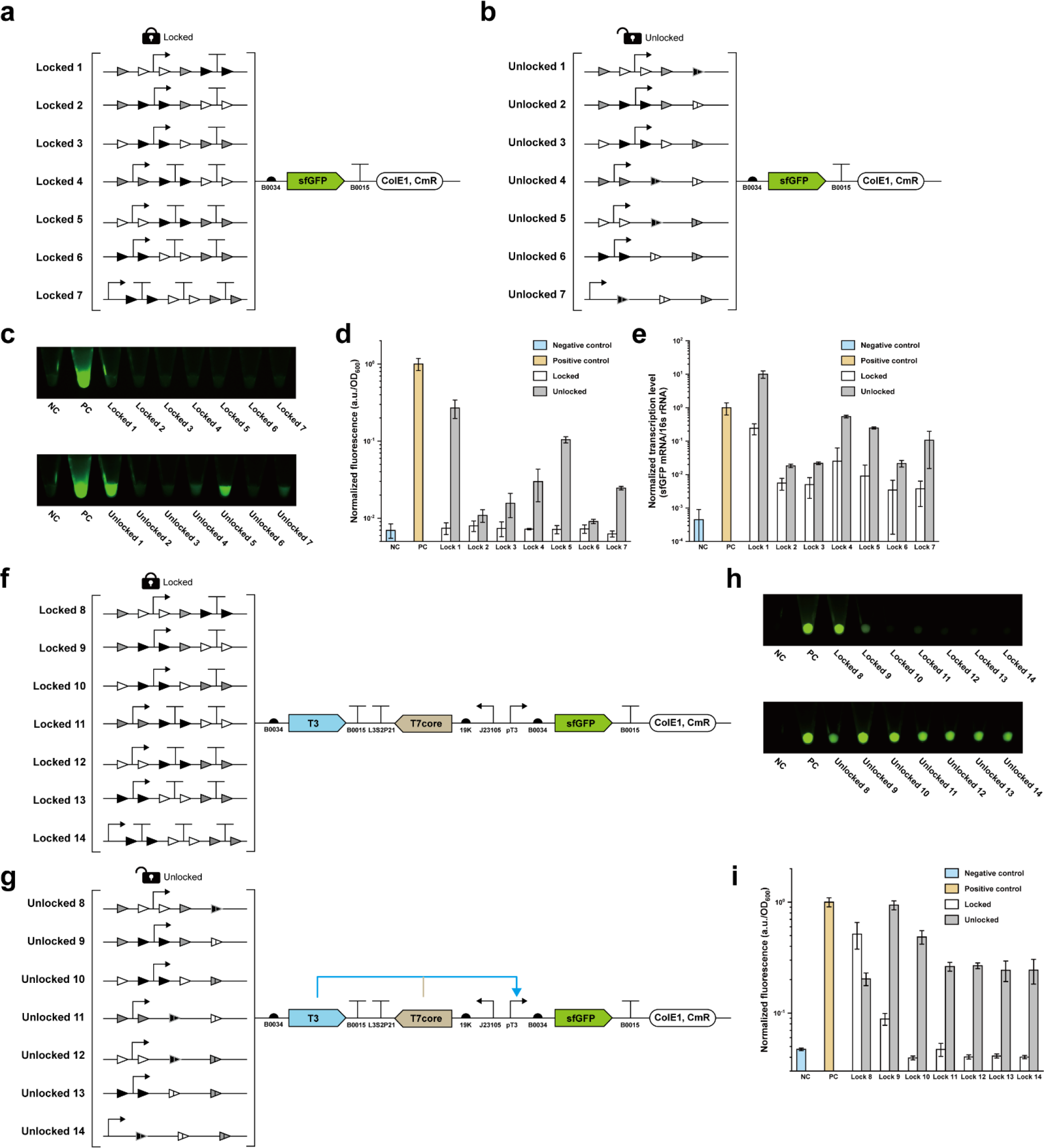
Quantitative analysis and enhancement of the SYMBIOSIS system. (**a**, **b**) Schematics of “Locked” (pFB34 to pFB40) and “Unlocked” (pFB41 to pFB47) plasmids for quantitative analysis (for details of all plasmids, see **Supplementary Table 5**). (**c**) sfGFP fluorescence of *E. coli* pellets. NC, negative control (pSB1C3); PC, positive control (pFB141). Each *E. coli* sample is collected in a 1.5 mL centrifuge tube from 3 mL overnight culture. All samples are repeated by three biological replicates with similar results. (**d**) Normalized sfGFP fluorescence values (a.u./OD600) corresponding to **c**. Fluorescence values are normalized to the one of positive control. Each value (mean ± s.d.) is calculated with three biological replicates. (**e**) Normalized transcription levels (sfGFP mRNA/16s rRNA) of the sfGFP gene. Transcription levels are normalized to the one of positive control. Each value (mean ± s.d.) is calculated with three biological replicates. (**f**, **g**) Schematics of improved “Locked” (pFB48 to pFB54) and “Unlocked” (pFB55 to pFB61) plasmids with signal amplification circuits (for details of all plasmids, see **Supplementary Table 5**). (**h**) sfGFP fluorescence of *E. coli* pellets. The experiment is performed the same as described in **c**. (**i**) Normalized sfGFP fluorescence values (a.u./OD600) corresponding to **h**. Fluorescence values are normalized to the one of positive control. Each value (mean ± s.d.) is calculated with three biological replicates.

However, their expression levels varied significantly and were lower than that of the positive control (**Fig. 3c** bottom panel, **3d**). This is in agreement with the expression results of chromoproteins (**Supplementary Fig. 9**), indicating that DNA sequence of the 5’-UTR region indeed affected the transcription and/or translation level. To quantify the transcription level, we carried out RT-qPCR experiments to calculate the ratio of sfGFP mRNA/16s rRNA in *E. coli* (**Fig. 3e**, **Supplementary Figs. 10**, **11**, **12**). Overall, the amount of sfGFP mRNA from the “Unlocked” circuits was much higher than the corresponding “Locked” ones, which turned out that the terminator(s) located between promoter and RBS significantly blocked the transcription process. Meanwhile, we also noticed that the levels of transcribed sfGFP mRNA from the “Locked” circuits were higher than that of the negative control. But the basal mRNA transcription leakage did not lead to detectable protein expression (**Fig. 3c** top panel, **3d**). This is likely because the translation process (e.g., the initiation stage) is completely blocked by the upstream 5’-UTR that contains both terminator(s) and multiple integrase recognition sites. In addition, we found that even if the transcription level of “Unlocked” circuit 1 was higher than the positive control (**Fig. 3e**), its sfGFP translation level was still lower (**Fig. 3d**). This result further suggested that residual recognition sites (*attL*/*attR*) could reduce the translation efficiency. Furthermore, we predicted the mRNA secondary structures of seven genetic circuits under “Locked” and “Unlocked” states. The models showed that stem-loop structures near the RBS sites (**Supplementary Fig. 13**, **14**) might limit ribosomes to bind RBSs^40, 41^. Taken together, our results demonstrated that diverse residual *attL*/*attR* sites (**Fig. 3b**) located in 5’-UTR played different influential roles on both transcription and translation levels (**Fig. 3d**, **e**).

In order to reduce the influence of residual *attL*/*attR* sites on the downstream protein translation, we next aimed to use “genetic amplifier” as a potential solution to enhance the output expression level^15^. Such strategy can be achieved by using a so-called transcriptional amplifier like the strong bacterial phage T7 RNA polymerase for cascaded signal amplification. However, direct overexpression of T7 RNA polymerase may lead to severe cell toxicity^42^. We, therefore, chose a reported “resource allocator” as our genetic amplifier^43–45^. The resource allocator consists of two parts: a non-active “core” fragment of T7 RNA polymerase and a T3 σ fragment that contains the DNA binding domain. Once the two fragments are expressed, they will bind together to form an active RNA polymerase to transcribe the pT3 promoter-mediated downstream gene sequence. For this, we modified our original “Lock” plasmids (**Fig. 3a**) by inserting both T3 σ and T7 core fragments between the “Locked” circuits and the sfGFP sequence (pFB48 to pFB54, **Fig. 3f**). In this case, the T7 core fragment was expressed from a constitutive promoter J23105. When the circuits were unlocked, T3 σ fragments would be expressed and bind to the T7 core fragments to transcribe the reporter sfGFP gene (**Fig. 3g**). By using this “genetic amplifier” strategy, the sfGFP expression levels were substantially improved. The “Unlocked” circuit 9 enabled the expression level as high as the positive control (**Fig. 3h** bottom panel, **3i**). However, it was unexpected that “Locked” circuit 8 also gave rise to notable sfGFP expression, which is even higher than the “Unlocked” circuit 8 (**Fig. 3h**, **i**). We ascribed this observation to two reasons. First, the leaky transcription (see the transcription level of “Lock” 1 in **Fig. 3e**) and translation of T3 σ fragments could form sufficient T3-T7 RNA polymerase to activate sfGFP synthesis. Second, overexpression of T3 σ fragments with the “Unlocked” circuit 8 might take up too much cellular resources, leading to a low sfGFP expression. Nonetheless, our results demonstrated that (i) the inhibiting effects of residual *attL*/*attR* sites if any on protein translation vary based on different DNA sequences (**Fig. 3d**); (ii) the final output signal in most cases can be amplified to a higher level by using a “genetic amplifier” (**Fig. 3i**); and (iii) our SYMBIOSIS system is highly adaptable and can be rationally modified and improved for other synthetic biology applications.

### Expanding SYMBIOSIS to assemble multiple plasmids *in vitro*

To expand the concept of SYMBIOSIS for synthetic biology applications, we next aimed to establish *in vitro* SYMBIOSIS for assembling multiple plasmids (**Fig. 4a**), which can be transformed into cells for *in vivo* protein expression. To do this, two groups of plasmids named “Acceptor” and “Donor” are initially designed. “Acceptor” plasmids have various *attB* sites. “Donor” plasmids consist of three parts: (i) the origin of replication (ori) of R6K, which prevents the plasmids from correct replication in *E. coli* strains without the gene *pir*^46^, (ii) an antibiotic resistance gene that is unique to the specific “Donor” plasmids, and (iii) an *attP* site, which can be recognized by serine integrase and mediates *attP*-*attB* site-specific recombination with an “Acceptor” plasmid. Furthermore, the “Donor” plasmids can carry target biobricks, for example, gene coding sequences. From this, the “Donor” plasmid can be integrated into the “Acceptor” backbone after the first-round recombination, forming a new combined “Acceptor-Donor” plasmid. This newly constructed plasmid can serve as an “Acceptor” in the second-round integration to accept another donor DNA sequence. After several rounds, different composite “Acceptor-Donor(s)” plasmids can be assembled for the subsequent transformation and gene expression.

**Fig. 4.**
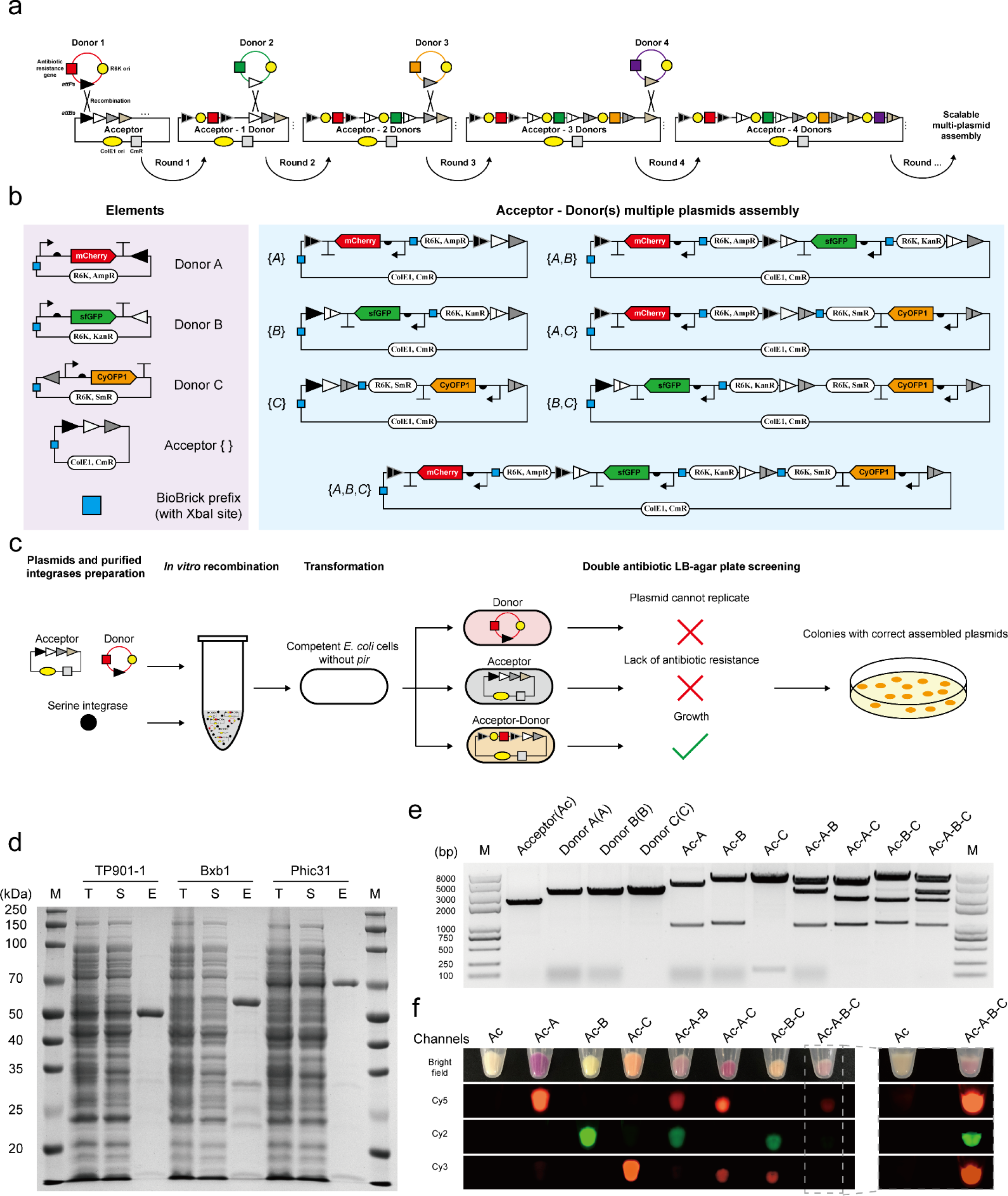
SYMBIOSIS enables *in vitro* multiple plasmids assembly. (**a**) Principle and design of scalable “Acceptor-Donors” assembly. In the first round, the “Donor 1” plasmid is integrated into the “Acceptor” backbone by one serine integrase, forming a combined “Acceptor - 1 Donor”, which can serve as a new “Acceptor” in the next round assembly. The “Acceptor” plasmid has different *attB* sites, which are paired to their cognate *attP* sites in the “Donor” plasmids. “Donor” plasmids use the same R6K ori and each one has its own unique antibiotic resistance gene, allowing for the subsequent plasmid screening. In addition, each “Donor” plasmid carries a target biobrick (e.g., gene coding sequence) for desired application. (**b**) A paradigm of *in vitro* SYMBIOSIS for multiple plasmids assembly. Left, three “Donor” plasmids (pFB63 to pFB65) harboring three fluorescent proteins, respectively, and one “Acceptor” (pFB62). Right, seven composite plasmids after assembly (pFB66 to pFB72). The details of these plasmids are shown in **Supplementary Table 5**. (**c**) Schematic workflow of *in vitro* SYMBIOSIS. (**d**) SDS-PAGE analysis of three purified serine integrases (TP901-1: 56.5 kDa, Bxb1: 57.2 kDa, and Phic31: 67.9 kDa). T, cell lysate; S, supernatant fraction; E, purified enzyme. (**e**) DNA fragment sizes of the “Acceptor”, “Donors”, and assembled plasmids after XbaI digestion. All expected bands are observed on the agarose gel (from left to right, pFB62: 2218 bp, pFB63: 3284 bp, pFB64: 3243 bp, pFB65: 3276 bp, pFB66: 4516+986 bp, pFB67: 4410+1051 bp, pFB68: 5341+153 bp, pFB69: 4410+3349+986 bp, pFB70: 5341+2451+986 bp, pFB71: 5341+2345+1051 bp, pFB72: 5341+3349+2345+986 bp). (**f**) Orthogonal fluorescence detection of *E. coli* with assembled plasmids. Four channels (bright field, Cy5, Cy2, and Cy3) are used to detect the fluorescence of all cell pellet samples. For *E. coli* containing the triple assembled plasmid (pFB72, Ac-A-B-C), the pellet sample is also independently detected with the negative control of “Acceptor” (pFB62, Ac) to show clear fluorescent signals. All samples are repeated by three biological replicates with similar results.

To demonstrate the capability of *in vitro* SYMBIOSIS, we first constructed three “Donor” plasmids with different *attP* sites and one “Acceptor” plasmid containing three sequential *attB* sites (**Fig. 4b**, left). In each plasmid, a “BioBrick prefix” part with one XbaI restriction site was inserted in an appropriate position to check the correctness of assembled plasmids. In addition, each “Donor” plasmid contained its own fluorescent protein gene, which can be integrated into the “Acceptors” to verify protein expression (**Fig. 4b**, right). As shown in the schematic workflow (**Fig. 4c**), “Donor” and “Acceptor” plasmids are mixed with purified serine integrases (TP901-1, Bxb1, and Phic31) for plasmid recombination *in vitro*. Then, the reaction mixture is transformed into *E. coli* (*pir^-^*) competent cells. By double antibiotic resistance screening, only the cells harboring the correctly assembled “Acceptor-Donor” plasmids can grow on the LB-agar plates (**Fig. 4c**).

The expression of three serine integrases was carried out in *E. coli* Rosetta-gami(DE3) pLysS. However, the initial expressed enzymes of TP901-1 and Phic31 were predominantly insoluble (**Supplementary Fig. 15**). Then, we switched to a low concentration of inducer and a low cultivation temperature (for details, see Methods). By doing this, all serine integrases were expressed in soluble forms and could be purified with high purity (**Fig. 4d**). Next, the conditions for *in vitro* plasmid recombination were investigated. The optimal reaction contained 1 μM of integrase and was performed at 30°C for 120 min (**Supplementary Fig. 16**). In total, seven assembled plasmids were generated (**Fig. 4b**, right) and their correctness was verified by XbaI digestion, each showing the correct digested DNA fragments on the gel (**Fig. 4e**). Afterwards, they were each transformed into *E. coli* cells, respectively, for protein expression. The results showed that all fluorescent protein genes on the relevant plasmids could be successfully expressed in single, in pairs, or in trios (**Fig. 4f**). This demonstrated that our SYMBIOSIS system was able to efficiently catalyze multiple plasmids assembly *in vitro*, providing a promising and reliable approach for plasmid construction.

**Fig. 5.**
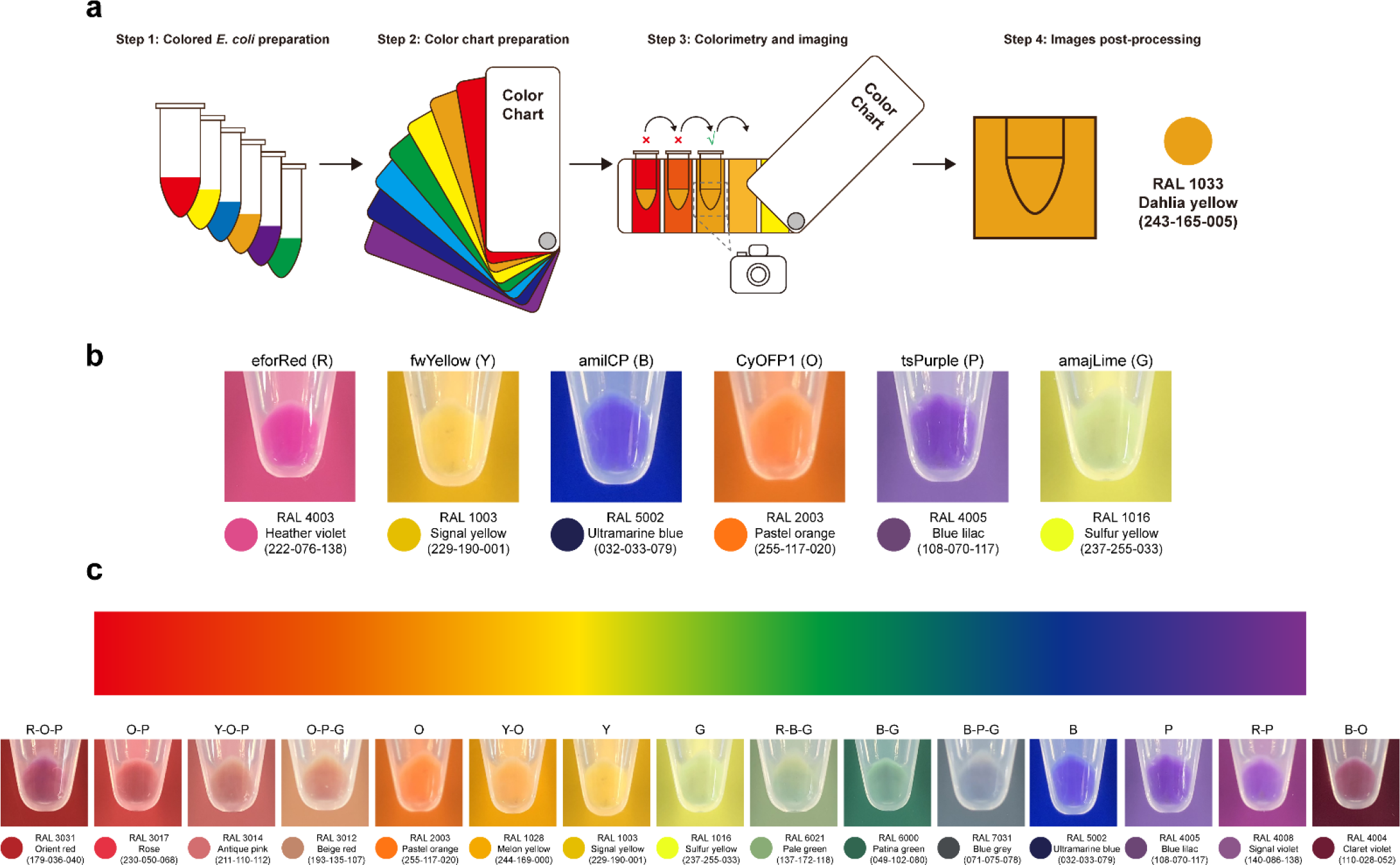
Creation of colored *E. coli* by SYMBIOSIS. (**a**) Workflow for the preparation of colored *E. coli* cells and colorimetry. Each *E. coli* sample is collected in a 1.5 mL centrifuge tube from 3 mL overnight culture and matched with the color chart. When the cell color matches the standard color, the image is taken in a mini photo studio (**Supplementary Fig. 17**) and further processed to convert the real cell color to a digital RGB color. (**b**) Six basic colors of red (R), yellow (Y), blue (B), orange (O), purple (P), and green (G) are generated with six chromoproteins (pFB90 to pFB95, **Supplementary Table 5**), respectively. They are used for double or triple combinations to create colored *E. coli* cells. (**c**) Representative colored *E. coli* cells spanning the color gradient. All samples are repeated by three biological replicates with similar results.

SYMBIOSIS creates colored *E. coli*

Following the demonstration of *in vitro* SYMBIOSIS, we next sought to further extend this proof-of-concept to create colored *E. coli* strains by recombination of different chromoprotein genes. While *E. coli* cells were able to show a featured color with a single chromoprotein (**Supplementary Fig. 7**), *E. coli* cells with composite colors have not been showed before. Therefore, we chose six different chromoproteins eforRed, fwYellow, amilCP, CyOFP1, tsPurple, and amajLime, which represent six basic colors of red, yellow, blue, orange, purple, and green, respectively. The goal was to use them to make various double or triple combinations with the hope to generate novel composite colors. To achieve this goal, the selected chromoprotein genes were individually constructed in three “Donor” vectors (pFB65, pFB73 to pFB89, **Supplementary Table 5**). Then, we ran *in vitro* SYMBIOSIS to generate 15 double and 20 triple combinations, leading to a total of 35 different expression plasmids.

To standardize the colorimetry process, we built up a step-by-step procedure for *E. coli* imaging and colorimetry (**Fig. 5a**). First, colored *E. coli* pellets were collected in transparent 1.5 mL Eppendorf tubes. Then, the color of cell pellets was compared to a color chart. When the best standard color was matched, the image was taken in a photo studio (**Supplementary Fig. 17**) and further edited by post-processing on a computer. This standardization procedure was performed for all color conversions from real cell colors to digital RGB colors. To begin, we compared the colors of *E. coli* pellets, each containing one of the six basic chromoproteins, with the color chart. We observed that each color matched well with the standard color and could be converted to a RGB color (**Fig. 5b**). Next, we used the six chromoproteins to make different combinations via *in vitro* SYMBIOSIS, yielding 35 assembled plasmids. Their correctness was verified by XbaI digestion and the sizes of digested DNA fragments were confirmed on the gel (**Supplementary Figs. 18**, **19**). All plasmids were transformed into *E. coli* for chromoprotein coexpression, generating a variety of colored cells (**Supplementary Figs. 20**, **21**). Most interestingly, some colors have never been observed before in *E. coli*, for example, orient red (pFB118), patina green (pFB107), blue grey (pFB129), and claret violet (pFB105). In particular, we were able to choose representative colors from the 6 basic colors and our 35 composite colors to span the color gradient (**Fig. 5c**). Taken together, these results demonstrated the power of using *in vitro* SYMBIOSIS to construct large, complex plasmids for defined applications such as the creation of novel colored *E. coli* cells in this work.

## Discussion

We demonstrate a serine integrase-based tool platform, called SYMBIOSIS, which can be used to directly and precisely manipulate DNA parts *in vivo* and *in vitro*. In one scenario, we propose a KmL model framework, in which three “Keys” (i.e., three orthogonal serine integrases TP901- 1, Bxb1, and Phic31) and their combinations can successfully unlock seven genetic circuits with high efficiency in *E. coli* cells. If the output signal (the protein expression level) is low due to the inhibitory effect of residual *attL*/*attR* sites, it can be amplified and enhanced by inserting a “genetic amplifier” to the “Lock” plasmids. In another scenario, we apply SYMBIOSIS *in vitro* to assemble multiple plasmids, which gives rise to a novel method that catalyze the assembly of circular plasmids rather than linear DNA fragments. Using this strategy, we construct 35 plasmids that carry different combinations of six chromoprotein genes. These plasmids are able to mediate *in vivo* protein coexpression, creating various colored *E. coli* cells with some novel, and never reported colors such as patina green, blue grey, and claret violet.

Overall, our work highlights the unique applications of serine integrases, which will be a promising addition to the current genetic toolkit of synthetic biology. Although we only use three serine integrases to verify the SYMBIOSIS system in this study, such a key concept may be applied to many other orthogonal phage integrases that have been increasingly discovered^30^. This will allow researchers to design and manipulate more complex genetic circuits and study their performance *in vivo*. Using serine integrases, we also establish a new strategy for *in vitro* multigene assembly based on “Donor” and “Acceptor” plasmids as direct inputs, but not linear PCR fragments. As compared to a previous “ACEMBL” method^28, 29^, which enables plasmid fusion via Cre-LoxP recombination but unavoidably generates plasmid variants due to only one kind of LoxP sites available, our SYMBIOSIS technology utilizes orthogonal serine integrases and each of them has a specific recognition site, making the plasmid architecture and the position of inserted genes more predictable. Moreover, we showcase the success of using *in vitro* SYMBIOSIS to assemble two or three chromoprotein genes on one plasmid for *in vivo* heterologous coexpression. This endows *E. coli* cells with featured new colors. As such, other new functionalities might be generated for *E. coli* to perform desirable applications, for example, construction of complex metabolic pathways when coexpression of all enzymes using multiple plasmids in one cell remains difficult or not possible.

Looking forward, we anticipate that SYMBIOSIS together with currently versatile genetic tools will further facilitate DNA manipulation for synthetic biology applications. Especially, our SYMBIOSIS, driven by serine integrases, can address two requirements. First, fewer integrases are able to mediate more genetic circuits *in vivo* by designing a logic relationship (e.g., the KmL model). Second, multicomponent DNA assembly *in vitro* enables tunable construction of one plasmid carrying multiple genes for executing *in vivo* biological functions. Building on these achievements, future developments of SYMBIOSIS could seek to discover and study more serine integrases, which will expand the repertoire of serine integrase-based tools and thus help advance the efforts to design, build, and test for fundamental and applied research in synthetic biology.

## Methods

### Strains, plasmids, primers, and reagents

The details of *E. coli* strains and plasmids used in this study are listed in **Supplementary Table 1** and **Table 5**, respectively. All general plasmids were checked correctly with Sanger Sequencing (GENEWIZ) unless otherwise noted. All assembled plasmids (pFB66 to pFB72, pFB96 to pFB130) were checked correctly by agarose gel electrophoresis after FastDigest XbaI (Thermo Scientific) digestion. Oligonucleotides were synthesized by GENEWIZ and listed in **Supplementary Table 6**. Q5 High-Fidelity DNA Polymerase (New England Biolabs), FastPure Gel DNA Extraction Mini Kit (Vazyme), and NovoRec plus One step PCR Cloning Kit (Novoprotein) were used for molecular cloning. DreamTaq Green PCR Master Mix (Thermo Scientific) was used for colony PCR. Lysogeny Broth (LB) liquid medium containes10 g tryptone (Oxoid), 5 g yeast extract (Oxoid), and 10 g sodium chloride (Titan/Greagent) in 1 L ddH2O. LB agar plate was prepared by adding 15 g agar (Sangon Biotech) per liter LB liquid medium. Antibiotic stocks (1000x) are 100 mg/mL ampicillin, 50 mg/mL kanamycin, 50 mg/mL streptomycin, and 34 mg/mL chloramphenicol.

### Genetic parts

All genetic parts (promoter, RBS, terminator, integrase recognition site, CDS, and 16s rRNA fragment) used in this study are listed in **Supplementary Table 2**. Short genetic parts (promoter, RBS, terminator, and integrase recognition site) were synthesized within the oligonucleotides (GENEWIZ) and inserted into PCR fragments during molecular cloning. 16s rRNA fragments were amplified from *E. coli* Mach1-T1 genome (GenBank: CP002185.1). The CDS of TP901- 1, Phic31, mKate2, CyOFP1, and sfGFP were synthesized by GENEWIZ. The CDS of mCherry, core-sz17, and sz18-σ-T3 were gifts from Dr. Chao Zhong Lab, ShanghaiTech University. The other CDS were collected from iGEM Registry Distribution kit (Spring 2016 and Spring 2017 distribution).

### Construction of general plasmids

All general plasmids (except assembled plasmids) were constructed by Gibson Assembly. In brief, all linear polymerase chain reaction (PCR) products (Q5 High-Fidelity DNA Polymerase, New England Biolabs) were extracted by gel extraction and then ligated using NovoRec plus One step PCR Cloning Kit (Novoprotein). After ligation, the reaction mixture was added to 50 μL of competent *E. coli* Mach1-T1 cells, which were incubated overnight on LB-agar plate. Then, DreamTaq Green PCR Master Mix (Thermo Scientific) was used for colony PCR. The possible PCR products were sent to GENEWEZ for Sanger Sequencing. Especially, the plasmids with repeated sequences were constructed with three to four fragments in one Gibson Assembly reaction. For example, pFB26 was constructed with four PCR fragments: fragment one (*attB*(Phic31) homolog-B0034-asPink-B0015-pSB1C3-J23100-*attP*(TP901-1)homolog) derived from pFB137, fragment two (J23100 homolog-*attP*(TP901-1)-B0015-*attB*(TP901-1)- *attP*(Bxb1) homolog), fragment three (*attB*(TP901-1) homolog-*attP*(Bxb1)-B0015- *attB*(Bxb1)-*attP*(Phic31) homolog) and fragment four (*attB*(Bxb1) homolog-*attP*(Phic31)- B0015-*attB*(Phic31)-B0034 homolog).

### Western-Blot assay

The *E. coli* Mach1-T1 strains containing compatible single, double or triple plasmids (pFB10, pFB13, pFB15) were cultivated in 5 mL LB liquid medium at 30°C and 250 rpm for 16 h. Cell pellets were collected in 1.5 mL centrifuge tubes at 5000 g and 4°C for 10 min. The supernatant was discarded and the pellets were resuspended with 1 mL 1x Phosphate-buffered saline (pH 7.4) and lysed by sonication (Q125 sonicator, Qsonica, 10 s on/off, 50% of amplitude, input energy ∼600 Joules). The lysate was then centrifuged at 12000 g and 4°C for 10 min. The total (cell lysate) and soluble (supernatant) fractions were separated by SDS-PAGE (Omni-Easy One-Step PAGE Gel Fast Preparation Kit, EpiZyme), followed by wet transferring to PVDF membrane (Bio-Rad) with 1x transfer buffer (25 mM Tris-HCl, 192 mM glycine and 20% v/v methanol in 1 L ddH2O, pH 8.3). Then, the PVDF membrane was blocked (Protein Free Rapid Blocking Buffer, EpiZyme) for 1 h at room temperature. After washing thrice with TBST for each 5 min, 1:10000 (TBST buffer based) diluted His-Tag Mouse Monoclonal Antibody (Proteintech) solution was added to the membrane and incubated for 1 h at room temperature. After washing thrice with TBST for each 5 min, 1:10000 (TBST buffer based) diluted HRP-Goat Anti-Mouse IgG (H+L) Antibody (Proteintech) solution was added to the membrane and incubated for another 1 h at room temperature. After the last washing with TBST thrice for each 5 min, the membrane was visualized using Omni ECL reagent (EpiZyme) under UVP ChemStudio (analytikjena).

### Assay of *in vivo* SYMBIOSIS for “Keys match Locks” manipulation

*E. coli* Mach1-T1 strains containing seven different “Keys” plasmid combinations (pFB10, pFB13, pFB15, pFB10+pFB13, pFB10+pFB15, pFB13+pFB15, and pFB10+pFB13+pFB15) worked as “Keys”, respectively. Each of the strains was transformed with the “Locks” plasmids pFB20-pFB26 for evaluating the “Keys match Locks” model. After “Locks” transformation, all forty-nine “unlocking” state strains were incubated overnight on LB-agar plates with appropriate antibiotics at 30°C for at least 24 h growth and *in vivo* recombination. The plates were photographed for recombination efficiency characterization and the matured colonies were then picked up for overnight incubation (each colony in 5 mL LB at 30°C and 250 rpm for 18 h). On the second day, *E. coli* Mach1-T1 pellets were collected from 3 mL culture in 1.5 mL centrifuge tubes and then photographed. The other 2 mL culture was used for plasmid extraction. After the “Keys match Locks” process, seven “Locked” strains would be chromoprotein-visualized, the other forty-two strains would kept locked and colorless. For electro-transformation, Gene Pulser Xcell system (Bio-Rad) and Gene Pulser/Micropulser Electroporation Cuvettes, 0.1 cm gap (Bio-Rad) were used in this study.

### Visualization and imaging of *E. coli*containing chromoproteins

*E. coli* Mach1-T1 strains containing chromoproteins unexpressed/expressed plasmids (pFB20 to pFB26, pFB27 to pFB33) were visualized and imaged under both bright field and LED Transilluminator (SLB-01W UltraSlim, MaestroGen) (see **Supplementary Figures 7**, **8** and **9**). All photos used in this study were taken by a mobile phone without post-processing.

### Recombination efficiency characterization of the “Keys match Locks” process

After the “Keys match Locks” process, matured colonies on the overnight LB agar plates were visualized and characterized with or without appropriate colors. When the “Unlocked” state (**Supplementary Figure 8**) occurred, colored colonies were counted as positive and few colorless colonies were counted as negative. The recombination efficiency was calculated as the positive percentage of total colonies. Each value was calculated with six biological replicates by counting at least 100 colonies on LB-agar plates.

### *E. coli* total RNA extraction

3 mL of each overnight cultured (30°C, 250 rpm, 18 h) *E. coli* Mach1-T1 pellet was collected in 1.5 mL centrifuge tube and then added 1 mL Trizol (Sangon Biotech) at room temperature for 10 min. 0.4 mL of phenol-chloroform-isoamyl alcohol (25:24:1, Aladdin) was then added and shaken vigorously for 15 s. Afterwards, the samples were centrifuged at 4°C and 12000 g for 10 min. The resulting supernatant (approximately 600 μL) was transferred to a new tube, mixed with equal volume of isopropanol, and stood at room temperature for 20 min. After centrifugation at 4°C and 12000 g for 10 min, the supernatant was discarded and the pellet was washed with 1 mL ethanol. Then, the sample was centrifuged again for 3 min at 4°C and 12000 The supernatant ethanol was discarded and total RNA (white flocculent precipitate) was collected at the bottom of the tube. After drying for 10 min at room temperature, the precipitate was resuspended in 100 μL nuclease-free water (Invitrogen). The concentration of total RNA was measured by the NanoDrop 2000/2000c Spectrophotometer (Thermo Scientific).

### RT-qPCR

1 μg of *E. coli* total RNA was used for reverse transcription to cDNA (Hifair III 1^st^ Strand cDNA Synthesis Kit (gDNA digester plus), Yeasen). 1 μL of cDNA product was used for qPCR (ChamQ SYBR Color qPCR Master Mix, Vazyme) quantification. Bio-Rad CFX96 Touch Real-Time PCR Detection System was used for the experiments and data analysis. For standard curve calculation, pFB141/pFB145 were used as sfGFP/16s rRNA quantification templates. Two plasmids were diluted in the range of 10^9^ to 10^4^ copies for calculation.

### Expression and purification of serine integrases

All three CDS of serine integrases TP901-1/Bxb1/Phic31 were constructed into pET22b vector with a *C*-terminal 6xHis tag and named as pFB142/pFB143/pFB144. Three serine integrases used in this study were expressed and purified by the following same protocol. The *E. coli* strain Rosetta-gami(DE3) pLysS was transformed with the relevant overexpression plasmid. Starter cultures (LB liquid medium containing ampicillin (100 μg/mL) and chloramphenicol (34 μg/mL) were inoculated from single transformant colonies and grown overnight at 37°C for 16 5 mL aliquot of the starter culture was used to inoculate 1 L LB medium containing ampicillin (100 μg/mL) and chloramphenicol (34 μg/mL) in a 2 L conical flask. The cultures were incubated at 37°C with shaking (250 rpm) until OD600 reached 0.6-0.8. The cultures were then cooled down on ice rapidly to 20°C, and isopropyl-β-D-thiogalactopyranoside (IPTG) was added for induction at a final concentration of 0.5 mM. The cultures were then grown for 16 h at 20°C and 220 rpm. After overnight induction, cells were harvested by centrifugation (Avanti JXN-26 High-Speed Centrifuge, Beckman Coulter) at 5000 g and 4°C for 10 min. The collected cell pellet (approximately 4 g) was resuspended by 25 mL 1x PBS and transferred into 50 mL centrifuge tubes. After a second centrifugation at 5000 g and 4°C for 10 min, the pellet was resuspended in 45 mL Buffer A, containing 20 mM sodium phosphate (pH 7.4), 1 M sodium chloride, 1 mM dithiothreitol (DTT), and 50 mM imidazole. The suspension was cooled on ice and then lysed thrice at 1500 bar by ultra-high pressure homogenization (JNBIO). The lysate was then centrifuged at 4°C and 20000 g for 30 min. The supernatant was collected and passed through a 0.22 μm filter. The filtered solution was then purified by Ni^2+^ affinity chromatography using a 1 mL HisTrap FF column (GE Healthcare). The column was equilibrated with 25 mL Buffer A (20 mM sodium phosphate (pH 7.4), 1 M sodium chloride, 1 mM dithiothreitol (DTT), and 50 mM imidazole), at a constant flow rate of 1 mL/min. After column equilibration, the filtered protein solution was loaded, followed by washing with Buffer A (approximately 25 mL). Then, bounded proteins were eluted with Buffer B (Buffer A, but with 500 mM imidazole) and collected into 1.5 mL centrifugal tubes with 0.5 mL elution. Each 0.5 mL elution was analyzed by SDS-PAGE. Eluted protein samples were combined in one tube and desalinated using an ultrafiltration tube (Amicon Ultra 3 kDa molecular weight cut-off, Merck/Millipore) with Buffer C (50 mM Tris-HCl (pH 7.5), 2 mM DTT, and 2 M sodium chloride). The desalinated and concentrated protein solution was then mixed with 40% glycerol (v/v in water,) with a volume ratio of 1:1. The concentration of the final protein solution was measured at 280 nm by the NanoDrop 2000/2000c Spectrophotometer (Thermo Scientific). Purified proteins were flash frozen by liquid nitrogen and stored at -80°C until further use.

### Optimization of *in vitro* recombination reactions and product analysis

Each of the three purified serine integrases was diluted in Buffer D (25 mM Tris-HCl (pH 7.5), 1 mM DTT, 1 M sodium chloride, and 20% v/v glycerol) to the final concentrations of 10 μM, 1 μM, and 0.1 μM. In a typical reaction, 3 μL of diluted serine integrase was added to a 27 μL reaction containing 400 ng of substrate plasmid (pFB16), 50 mM Tris-HCl (pH 7.5), 0.1 mM EDTA, and 100 μg/mL bovine serum albumin. Then, the recombination reaction was incubated at various temperatures (16°C/30°C/37°C) and time intervals (10 min/30 min/60 min/90 min/120 min/150 min) for optimization. The reactions were terminated by heating at 80°C for 20 min, then the samples were separated by agarose gel electrophoresis and visualized under UVP ChemStudio (analytikjena). Darker bands (except the plasmid bands) on agarose gel indicated a higher recombination efficiency.

### *In vitro* SYMBIOSIS for multi-plasmid assembly

For each *in vitro*SYMBIOSIS reaction, only one of the purified serine integrases was used for assembly between one “Donor” plasmid and one “Acceptor series” plasmid. The “Acceptor series” plasmids refer to three groups: “empty” Acceptor (pFB62), “one Donor assembled” Acceptors (pFB66 to pFB68, pFB90 to pFB95), and “double Donors assembled” Acceptors (pFB69 to pFB71, pFB96 to pFB110). 3 μL of diluted serine integrase (10 μM) was added to a 27 μL solution containing “Donor” plasmid and “Acceptor series” plasmid (each plasmid was added as “base pair/10” ng, for example, 221.8 ng of the plasmid pFB62 (2218 bp) was added per reaction), 50 mM Tris-HCl (pH 7.5), 0.1 mM EDTA, and 100 μg/mL bovine serum albumin. Then, the recombination reaction was incubated at 30°C for 120 min, followed by termination with heating at 80°C for 20 min. Afterwards, 10 μL of the reaction mixture was added to 100 μL *E. coli* Mach1-T1 competent cells for chemical transformation. To screen the correctly assembled plasmids, LB agar plates were supplied with multiple antibiotics matched to the multiple antibiotic resistant genes on the correctly assembled plasmids. Only the colonies harboring assembled plasmids could grow on the plates, while the colonies containing unreacted “Donor” or “Acceptor series” plasmids did not.

### Imaging of colored *E. coli* pellets

3 mL of each overnight cultivated (37°C, 250 rpm, 18 h) *E. coli* Mach1-T1 cells was collected in 1.5 mL centrifuge tubes and then visualized under bright field and imager (ChemiDoc MP Imaging System, Bio-Rad). For imaging the fluorescence of mCherry, sfGFP, and CyOFP1, three different programs were used, respectively, which are Cy5 (light: Red Epi illumination, filter: 695/55 Filter), Cy2 (light: Blue Epi illumination, filter: 530/28 Filter), and Cy3 (light: Green Epi illumination, filter: 605/50 Filter). All the samples were photographed in one photo without post-splicing.

### Colorimetry and imaging of composite colored *E. coli*

A RAL K7 classic color chart was used for colorimetry with composite colored *E. coli* Mach1- T1 pellets. 3 mL of each overnight cultivated (37°C, 250 rpm, 18 h) composite colored *E. coli* Mach1-T1 cells was collected in 1.5 mL centrifuge tubes and then colorimetry and imaging under bright field. The tubes were continuously comparing to different color charts unless the most matched situation occurred. Then, the photos (the tube was placed in the middle of photo and the matched color chart acted as background) were taken in a mini photo studio (**Supplementary Figure 17**) with a mobile phone without post-processing.

## Acknowledgements

This work was supported by grants from the National Natural Science Foundation of China (Nos. 31971348, 31800720, and 32171427) and the Natural Science Foundation of Shanghai (No. 19ZR1477200). J.L. also acknowledges the starting grant of ShanghaiTech University.

## Author contributions

J. L. and F. B. designed the experiments. F. B. performed all experiments. X. T. performed protein analysis experiments. F. B. analyzed the data and drafted the manuscript. J. L., Y. L., and W. Q. L. revised and edited the manuscript. J. L. conceived and supervised the study. All authors read and approved the final manuscript.

## Supplementary Information

### I. Supplementary Tables

**Supplementary Table 1.**
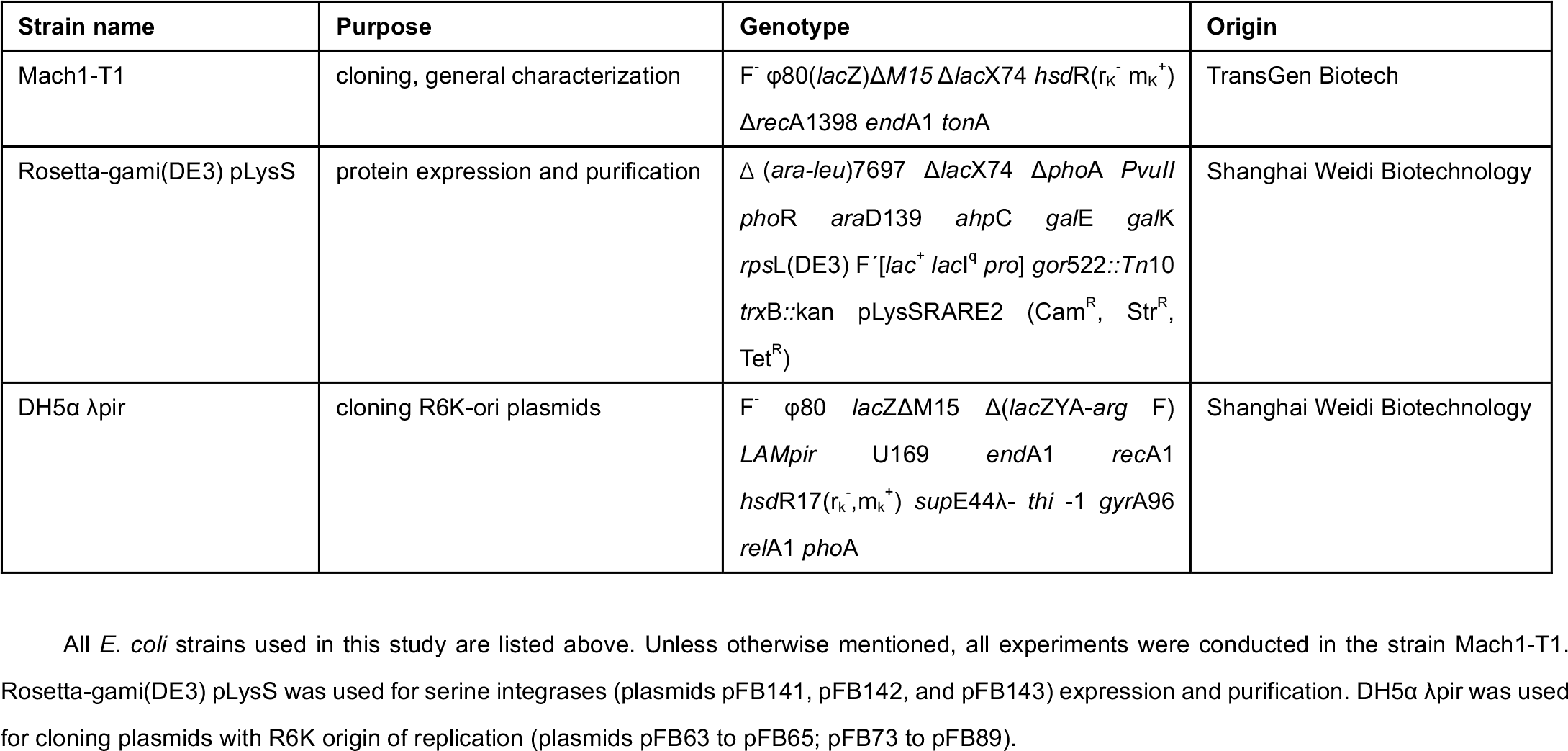
*E. coli* strains.

**Supplementary Table 2.**
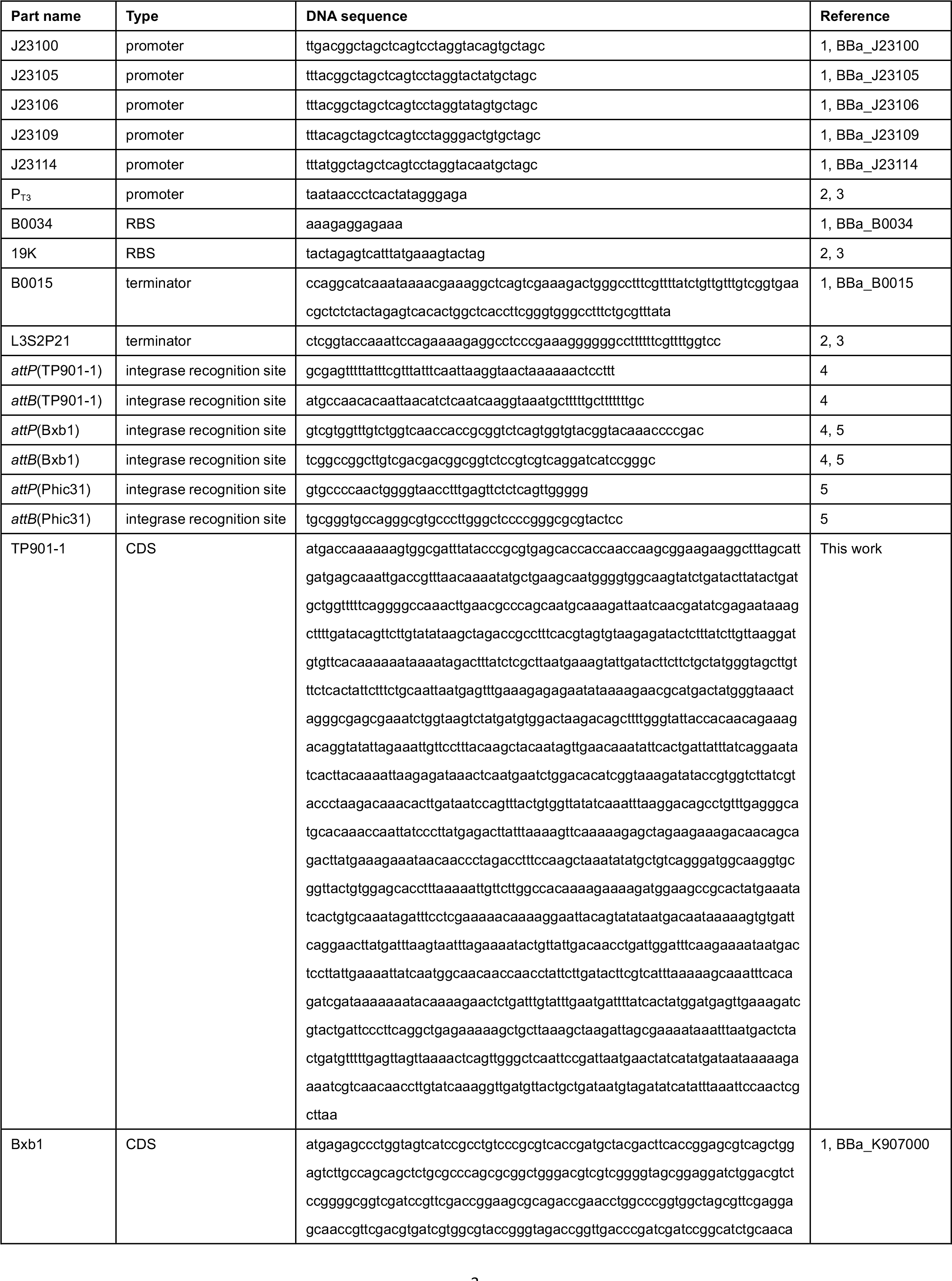
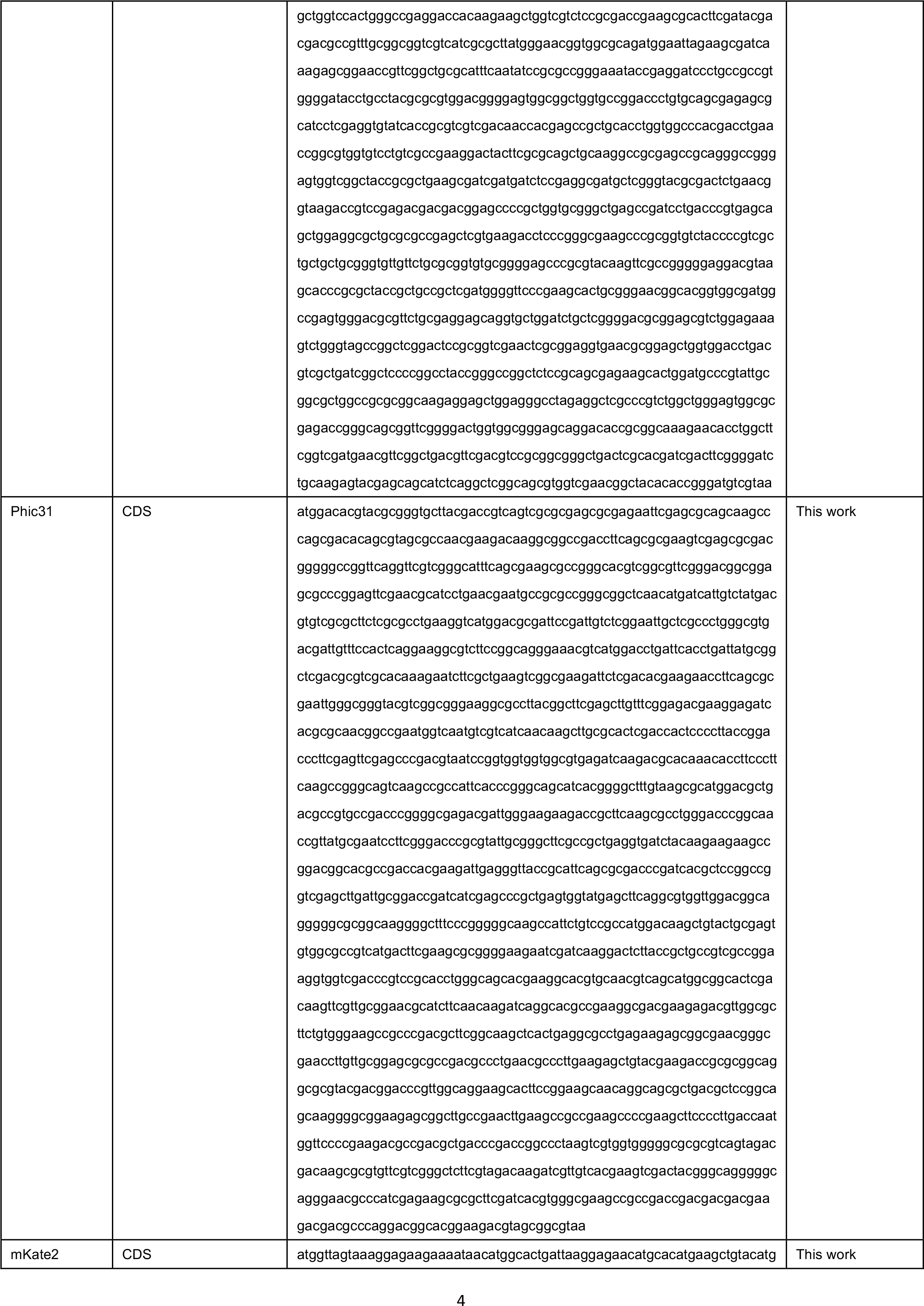
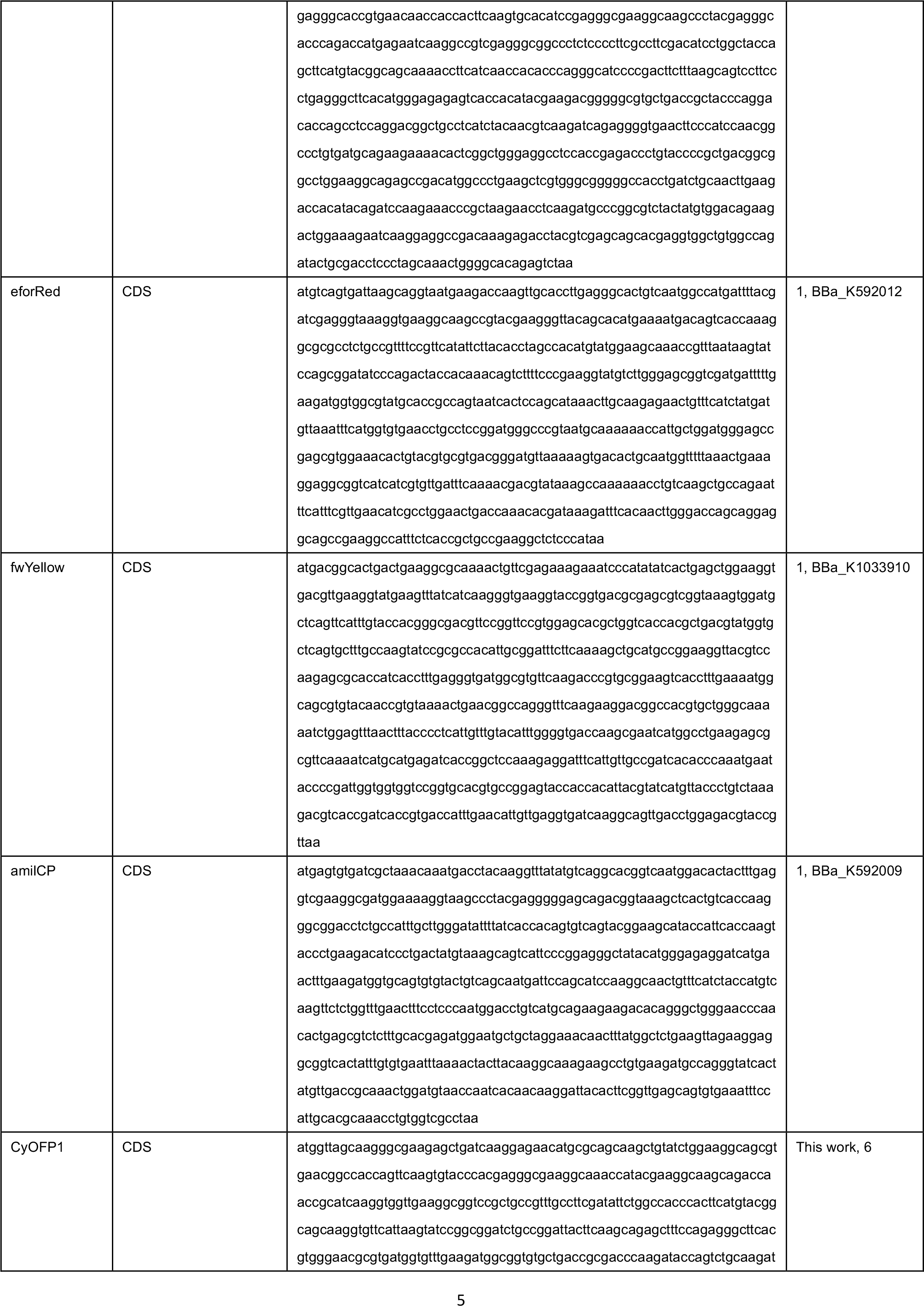
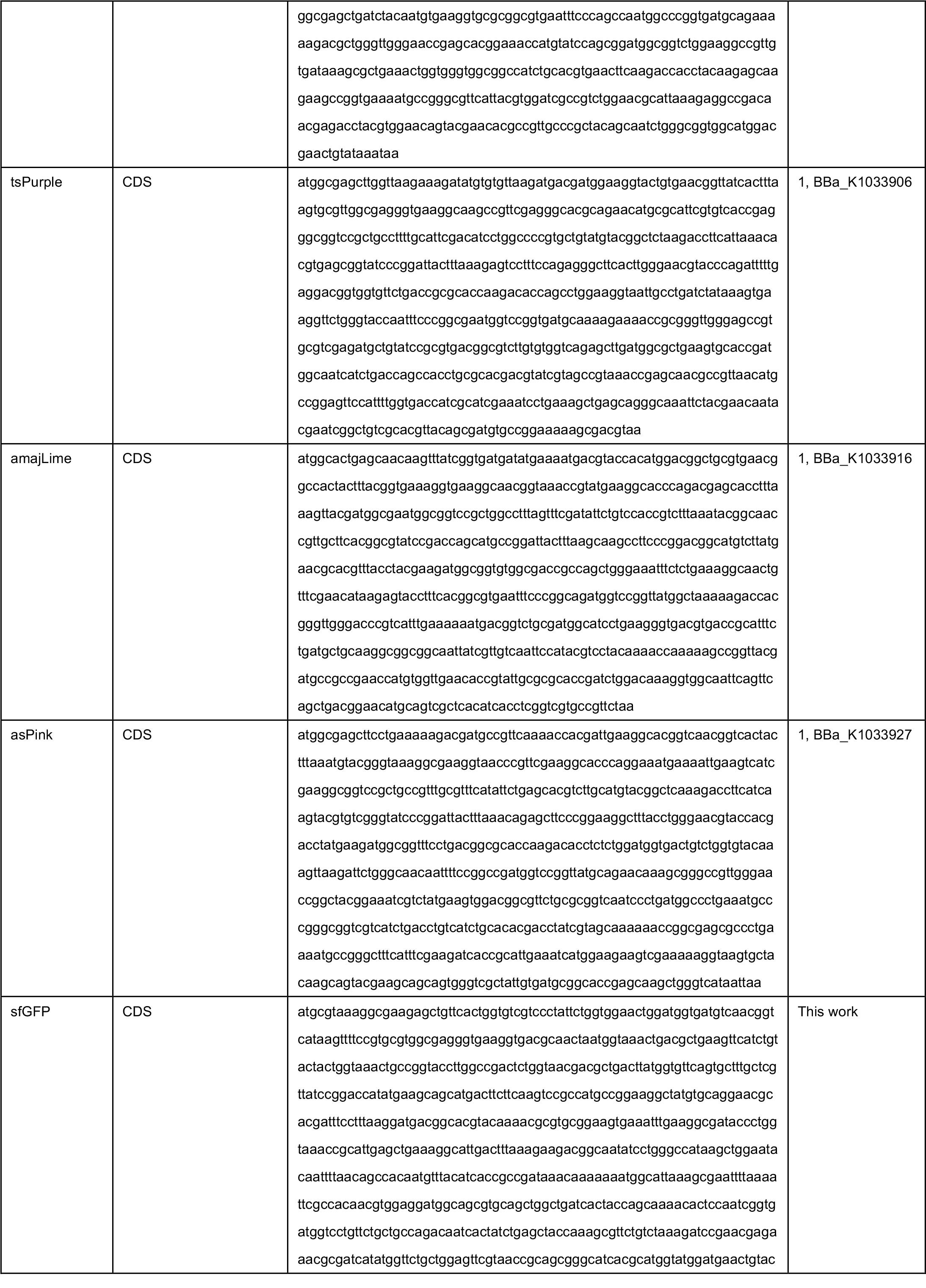
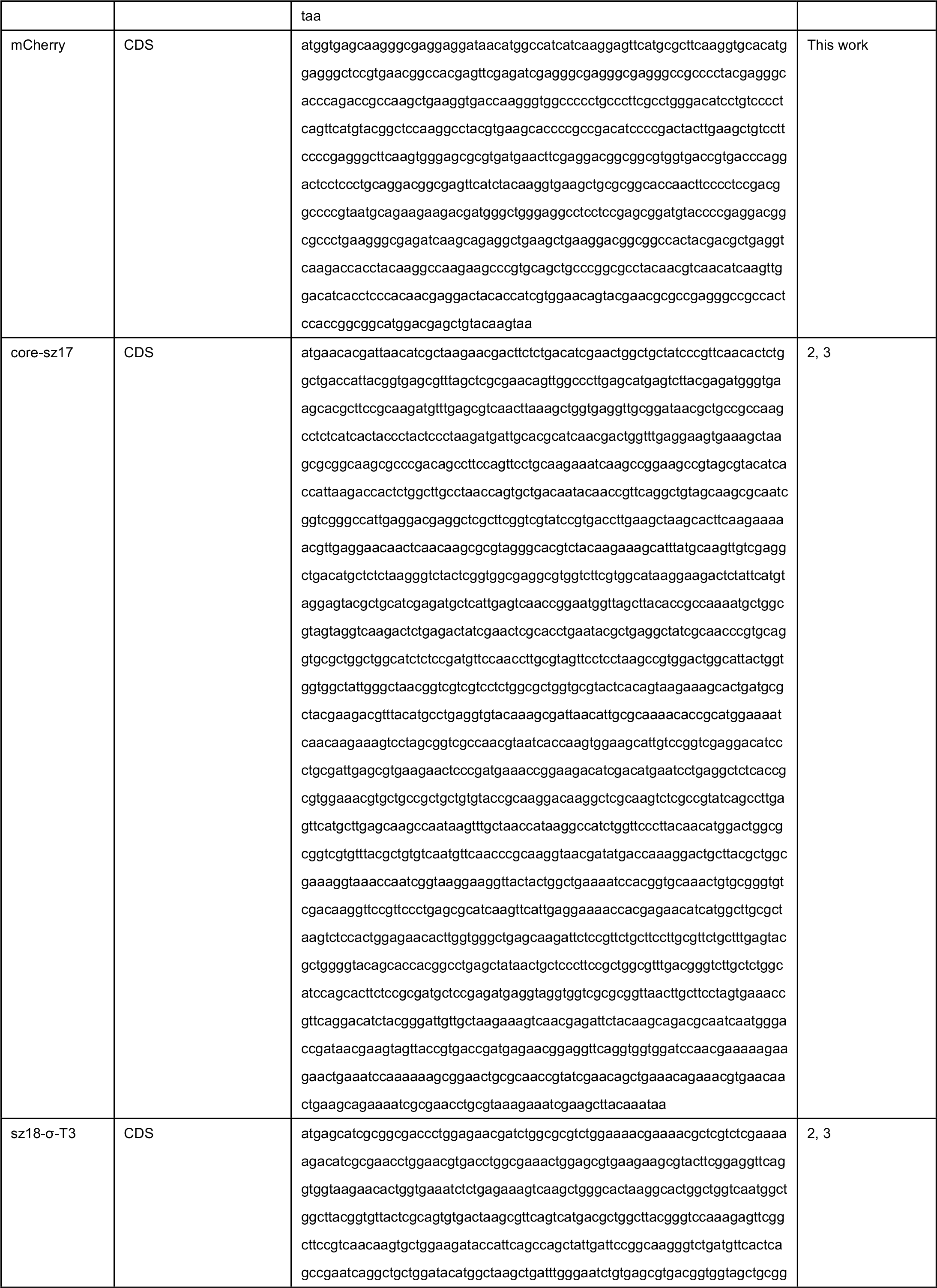
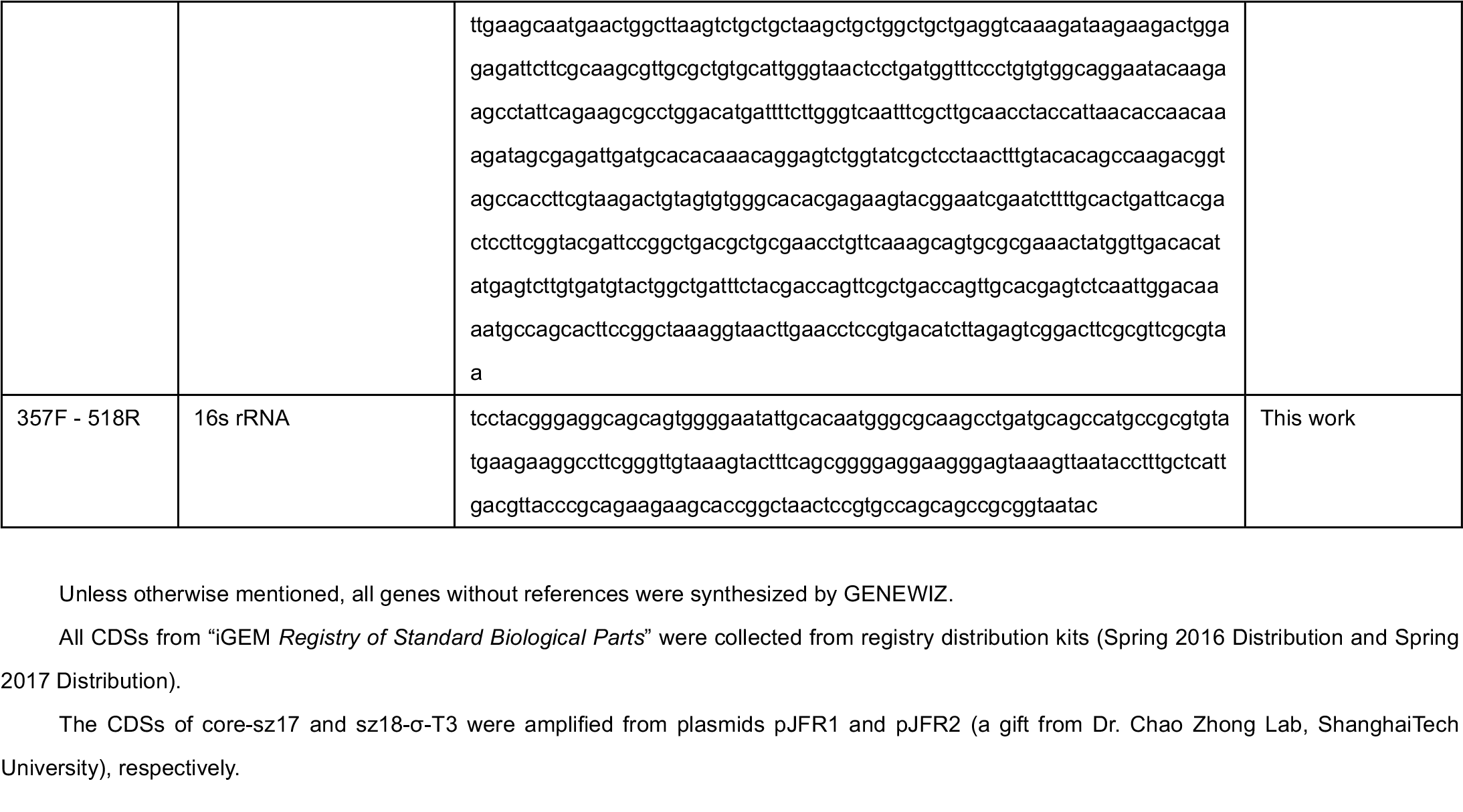
Genetic parts.

**Supplementary Table 3.**
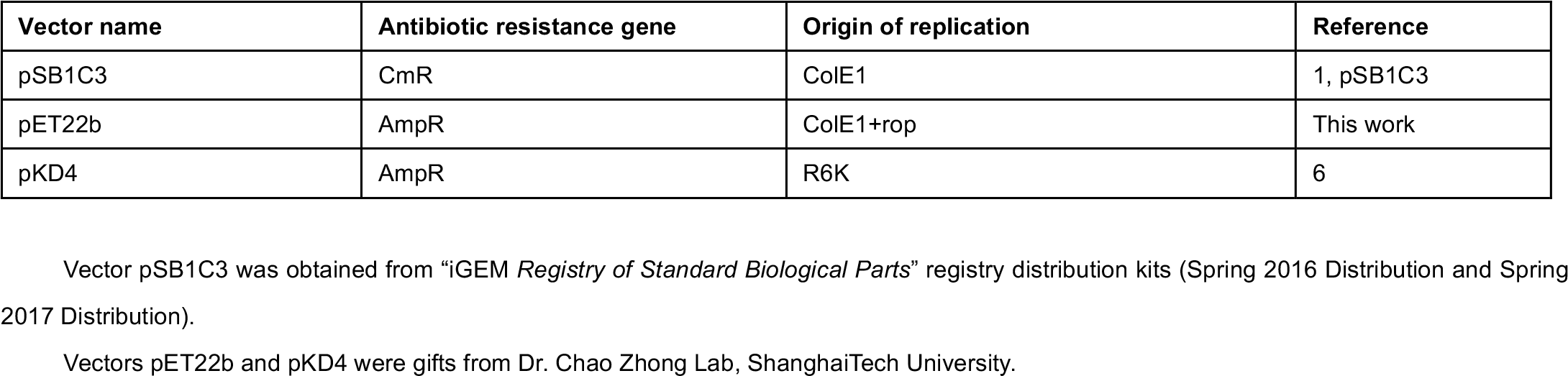
Vectors.

**Supplementary Table 4.**
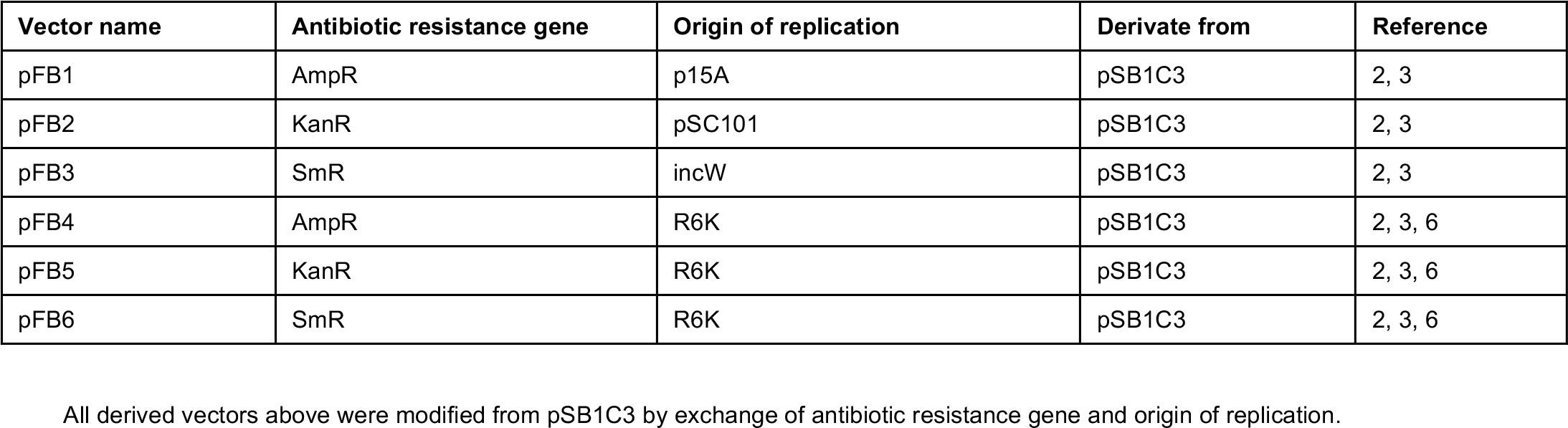
Derived vectors.

**Supplementary Table 5.**
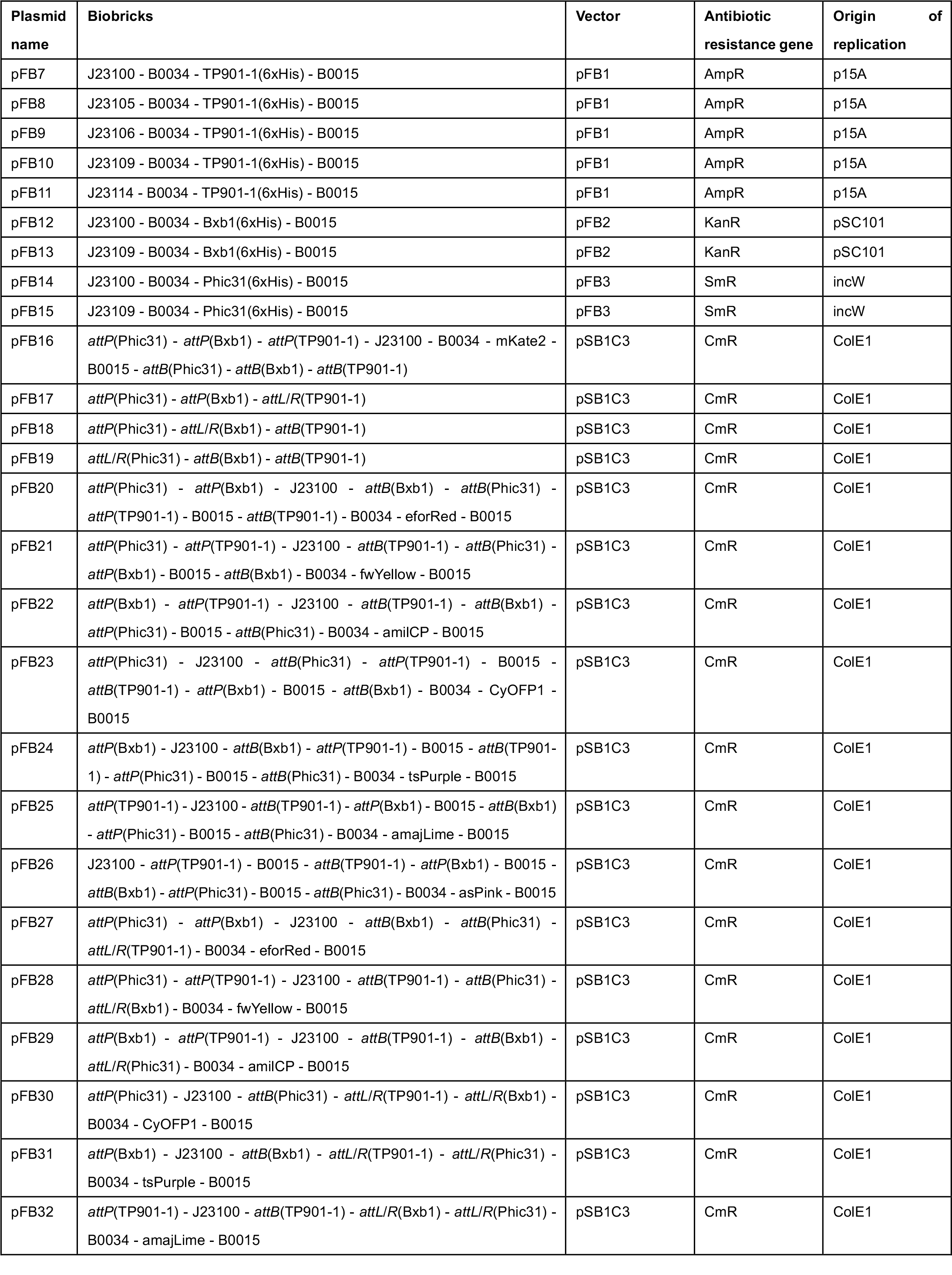
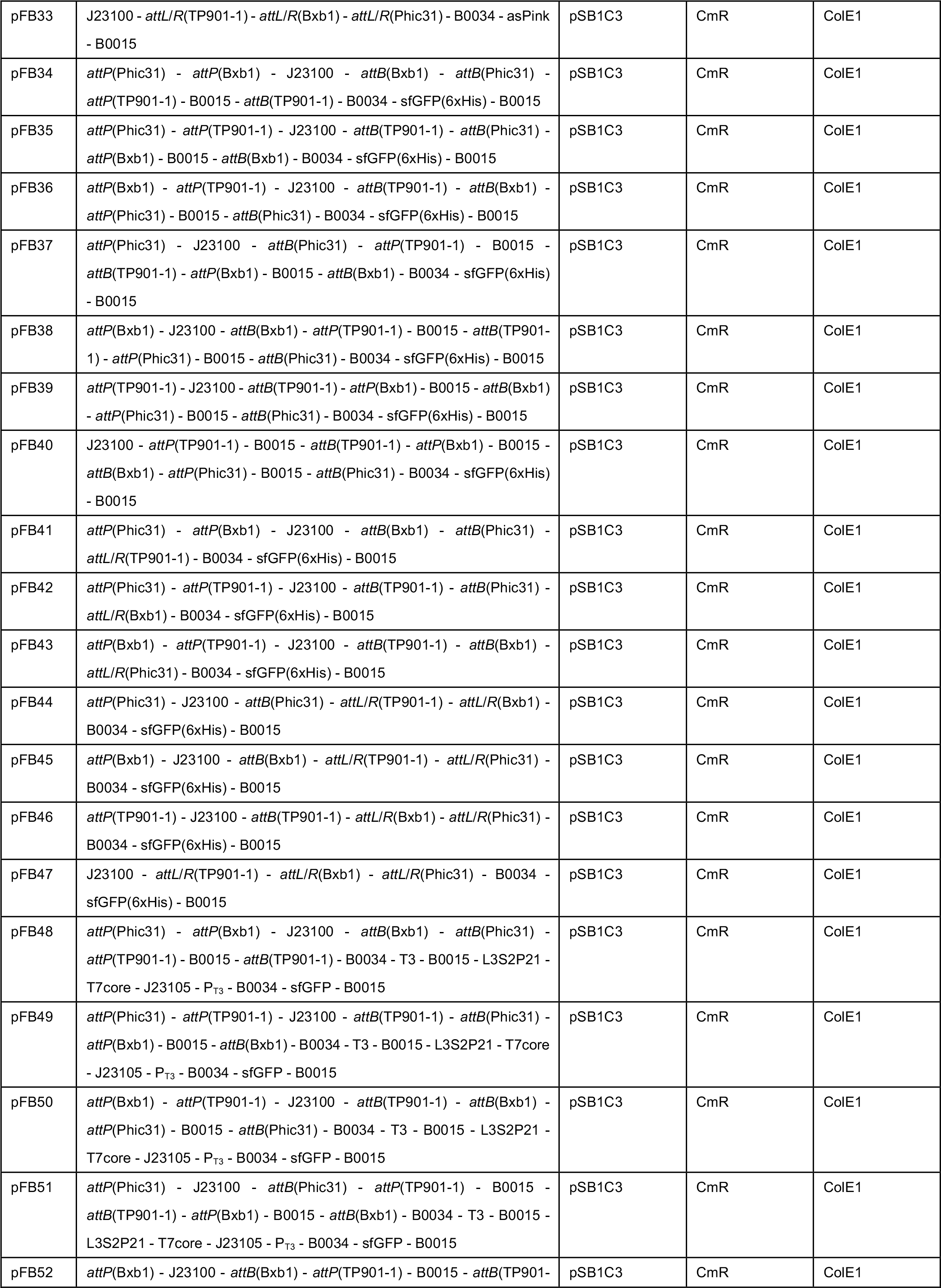
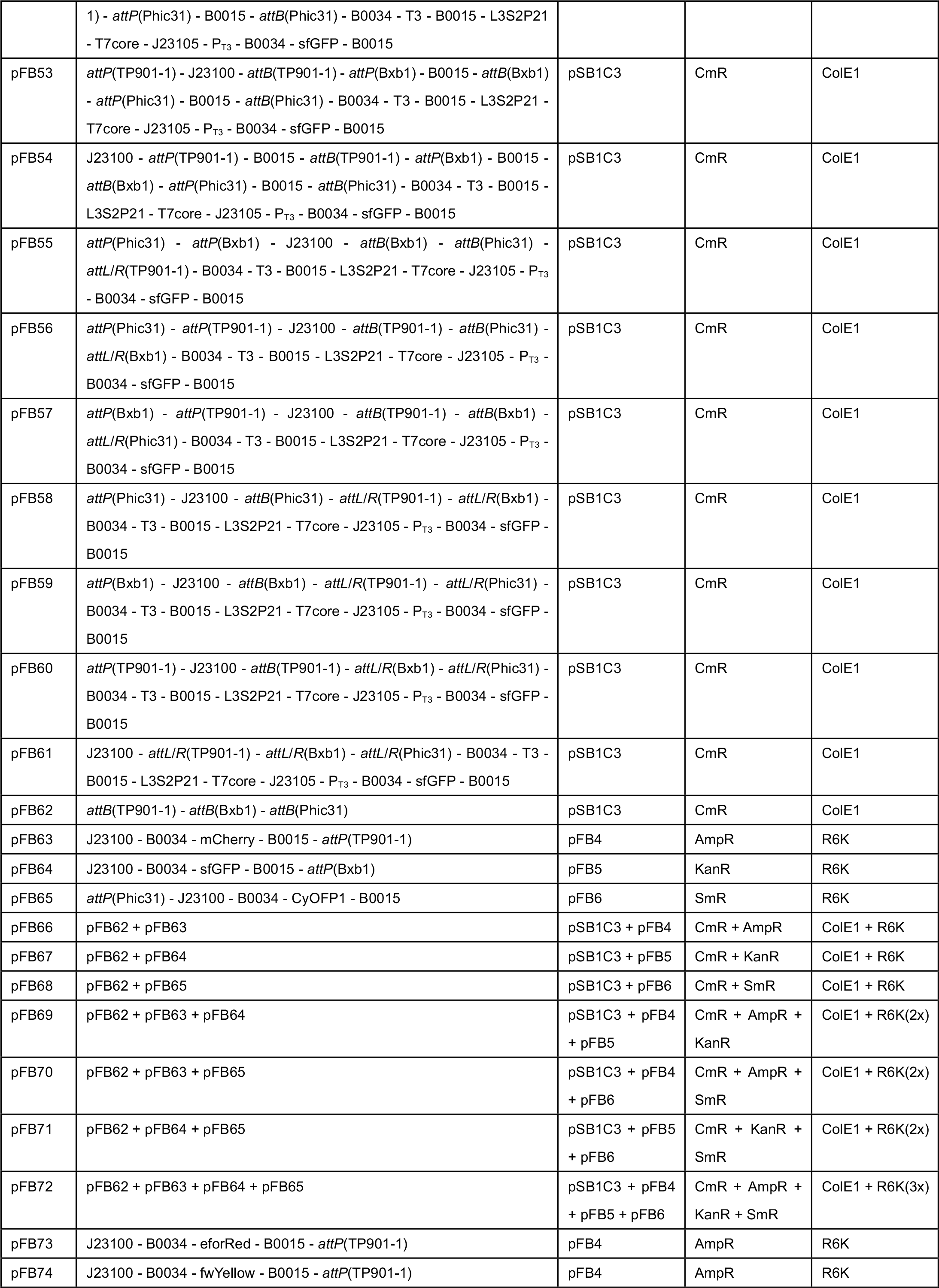
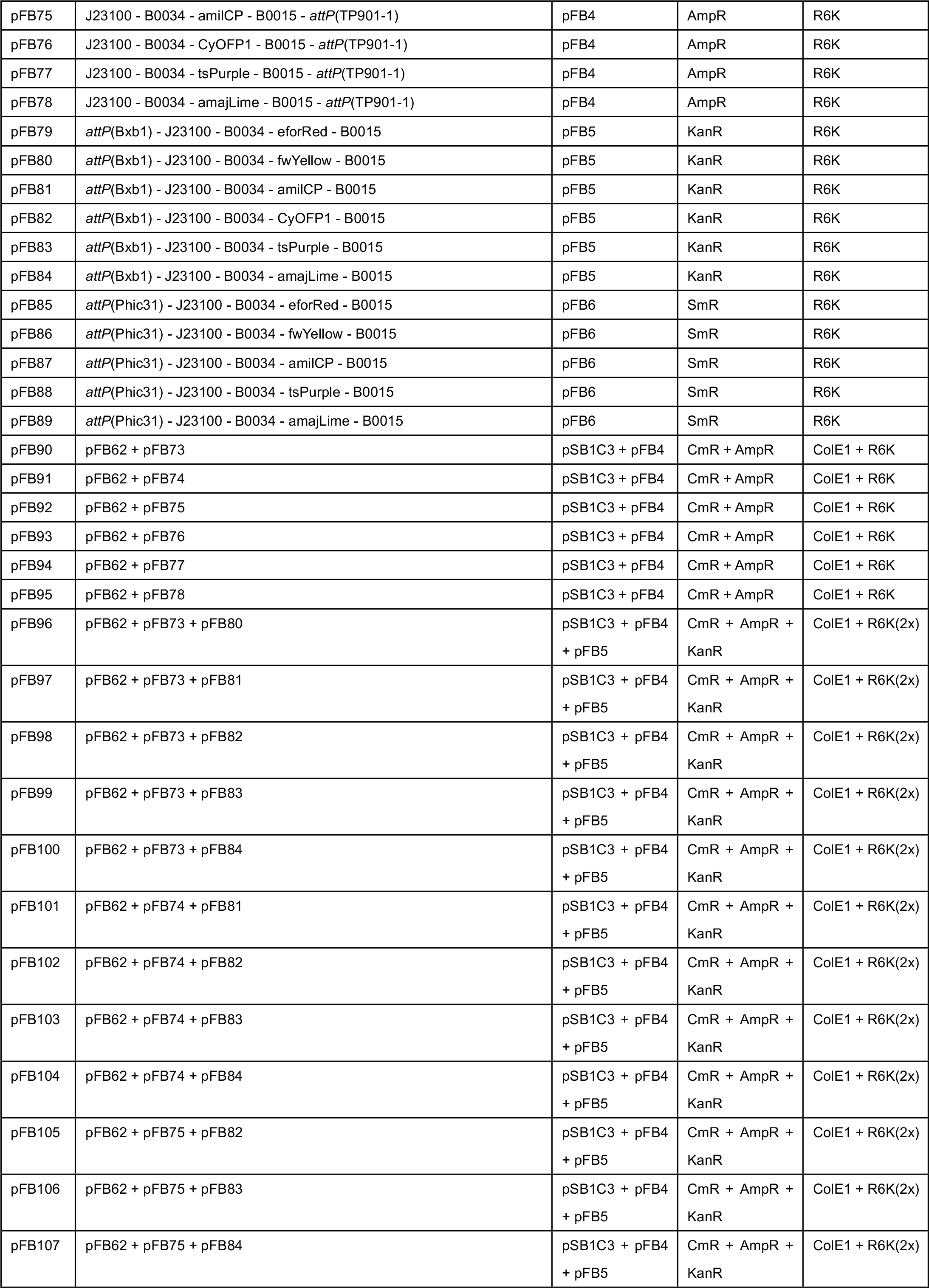
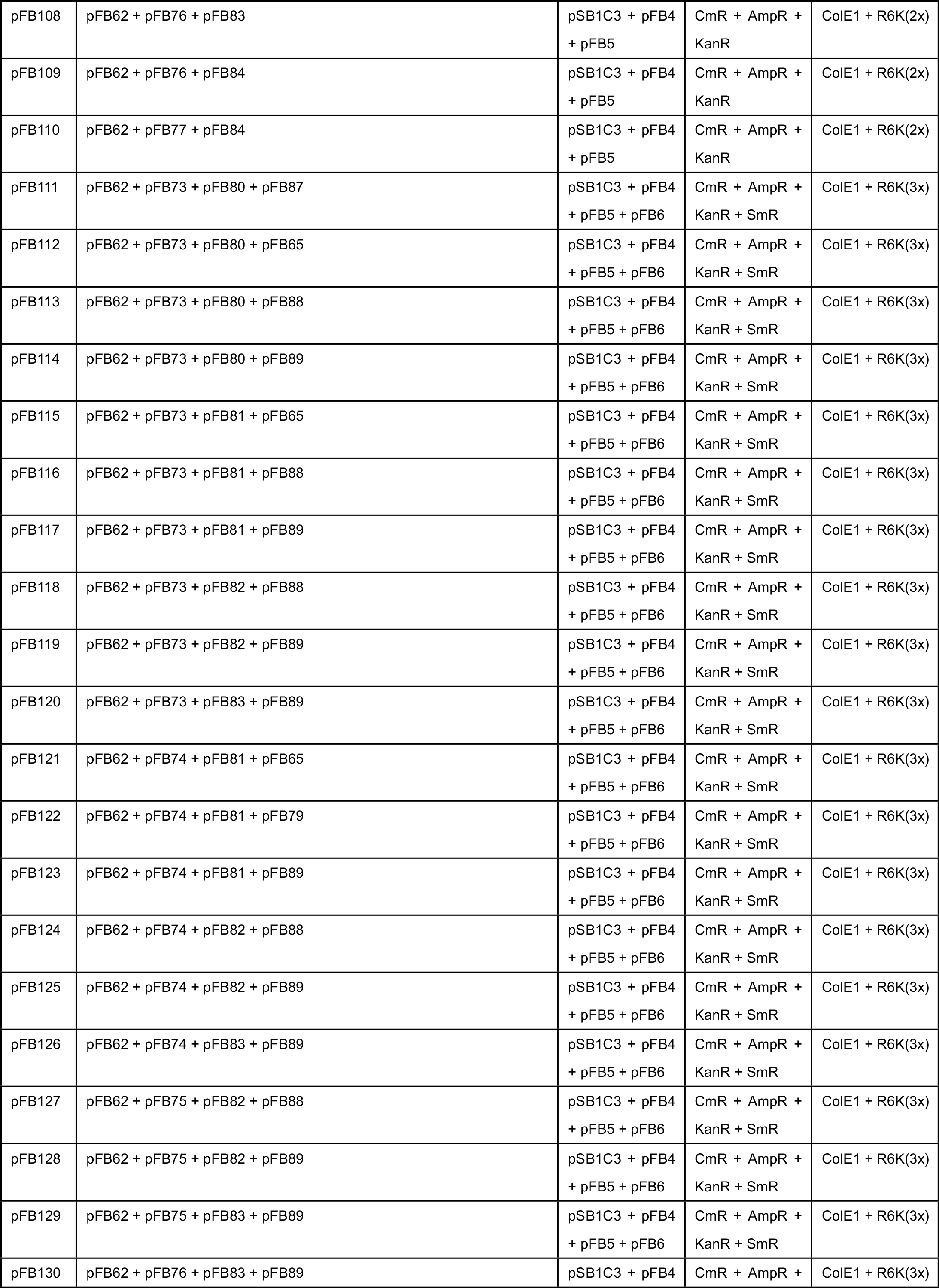
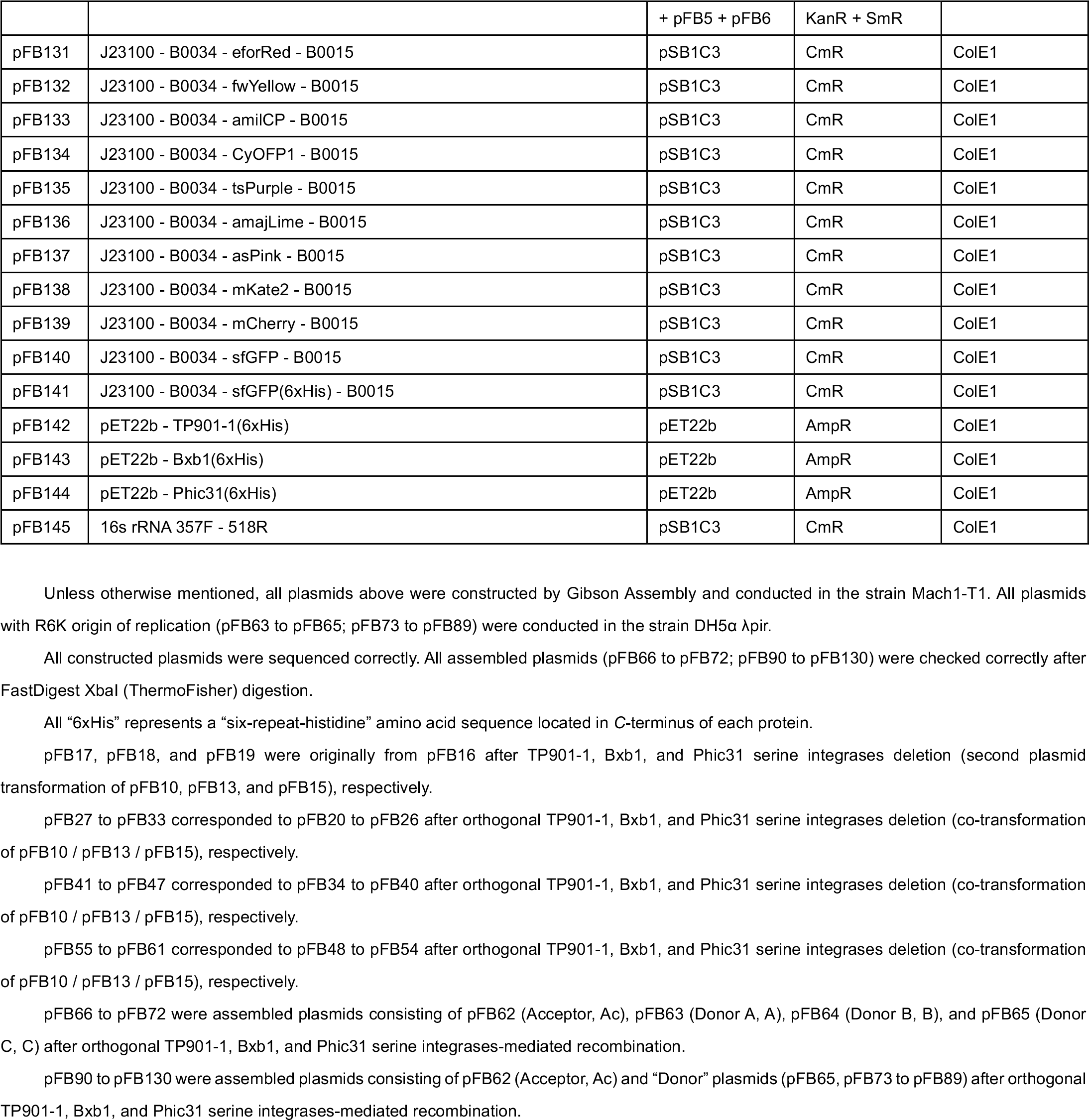
Plasmids.

**Supplementary Table 6.**
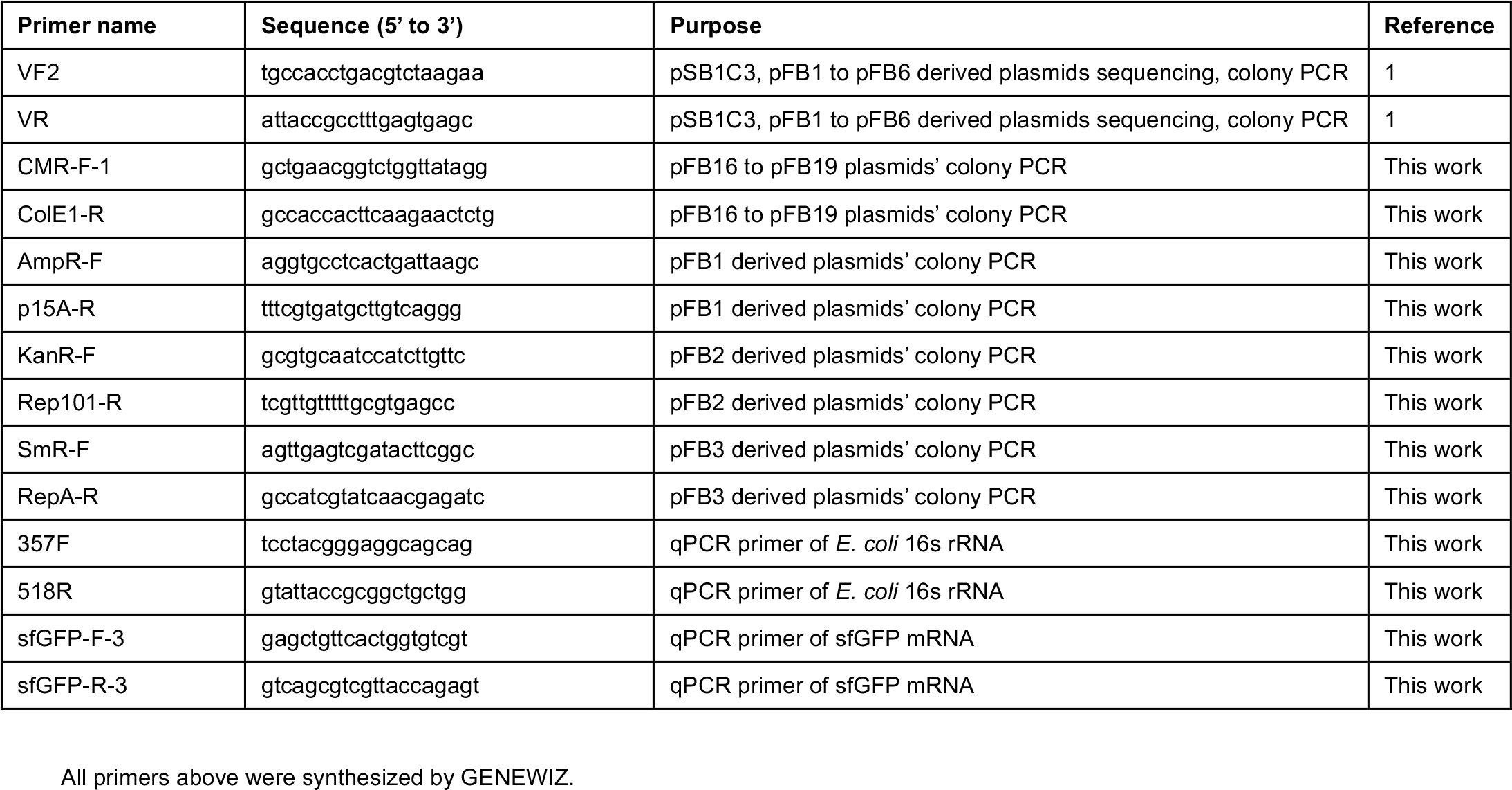
Primers.

### II. Supplementary Results

**Supplementary Figure 1.**
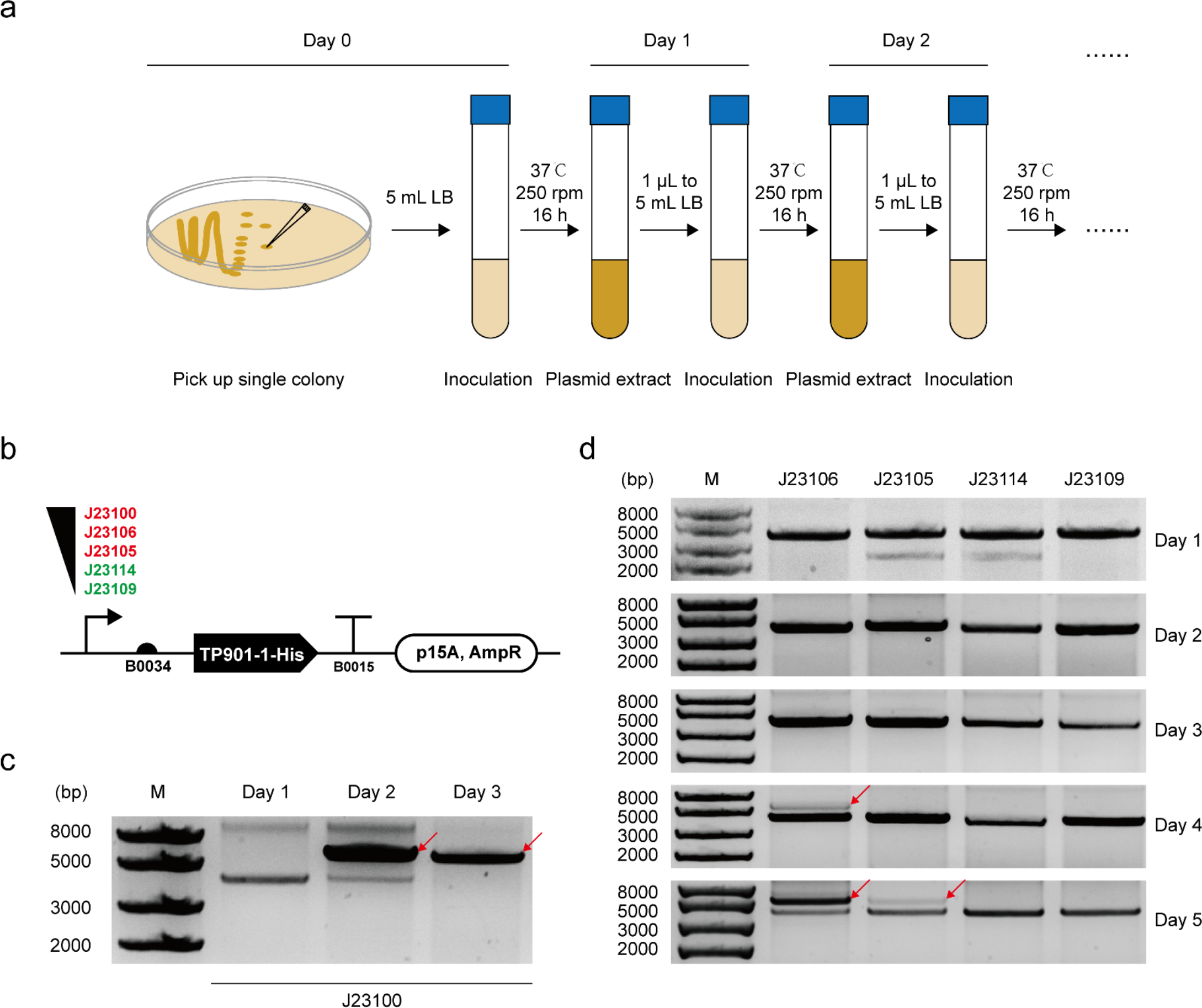
*In vivo* TP901-­-1 overexpression causes plasmid mutation. (**a**) Plasmids were extracted each day during serial dilution/cultivation. Extracted plasmids were separated by agarose gel electrophoresis after FastDigest XbaI (Thermo Scientific) digestion. (**b**) Schematic of TP901-­-1 constitutive expression plasmids. A series of constitutive promoters (J23100, J23106, J23105, J23114, and J23109 with transcription strength from strong to weak) were selected and assembled with common residual parts (B0034-­-TP901-­-1(6xHis)-­- B0015 on the derived vector pFB1). (**c**) The strongest TP901-­-1 expression plasmid pFB7 was mutated on Day 2. An unexpected incorrect band (greater than 5000 bp, red arrow) appeared except for the correct band (4007 bp). On Day 3, plasmids were completely mutated showing only one incorrect band (red arrow). (**d**) Other plasmids (pFB8 to pFB11) were checked similarly. The second and the third strongest expression plasmids (pFB9/J23106 and pFB8/J23105) were mutated as well on Day 4 and Day 5 (red arrows), respectively. Mutations of the other two plasmids pFB11 (J23114) and pFB10 (J23109) were not detected from Day 1 to Day 5.

**Supplementary Figure 2.**
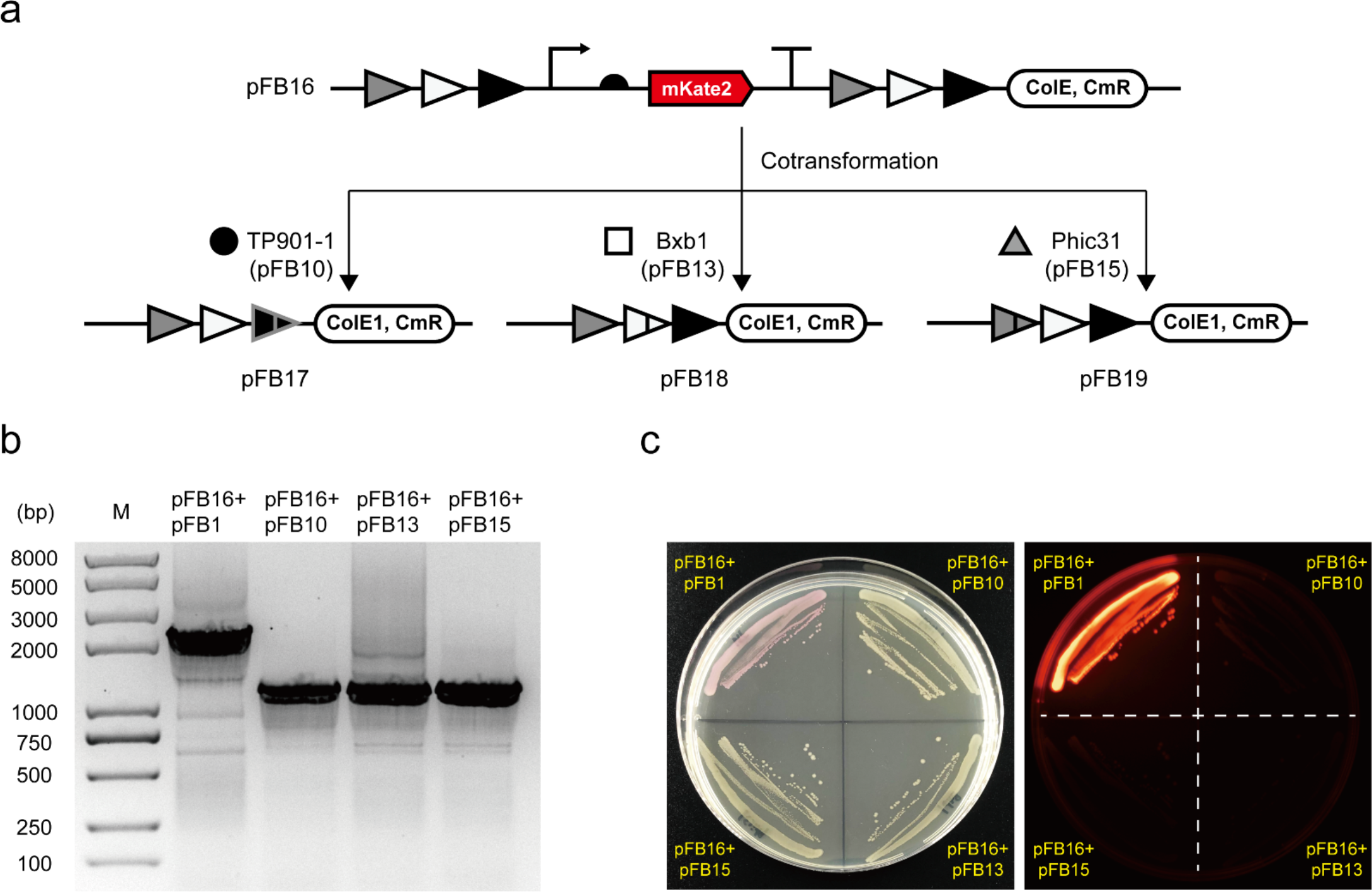
Characterization of constitutive serine integrases expression plasmids. (**a**) Schematic of characterization process. pFB16 was derived from pFB138 (J23100-­-B0034-­-mKate2-­-B0015) with assembly of three orthogonal *attP*/*attB* site pairs. Plasmid co-­-transformation of pFB16 with pFB10/pFB13/pFB15 mediated *attP*-­-*attB* deletion and then converted to pFB17/pFB18/pFB19, respectively. (**b**) Image of a gel post electrophoresis showing orthogonal serine integrases mediated *attP*-­-*attB* deletion. After co-­-transformation (pFB16+pFB10, pFB16+pFB13, pFB16+pFB15), colony PCR (primer pair: CMR-­-F-­-1/ColE1-­-R) showed a high efficiency of three orthogonal serine integrases. Expected PCR products after TP901-­-1/Bxb1/Phic31 deletion were observed (1213 bp of pFB17, 1207 bp of pFB18, and 1206 bp of pFB19), respectively, rather than the negative control (2277 bp of pFB16). (**c**) Agar plate image of *E. coli* growth with co-­-transformed plasmids. *E. coli* cells harboring pFB16 were transformed with pFB1 (negative control, empty vector), pFB10 (low level constitutive TP901-­-1 expression), pFB13 (low level constitutive Bxb1 expression), and pFB15 (low level constitutive Phic31 expression), respectively. The resulting *E. coli* cells were inoculated on each quarter area of the same LB-­-agar plate (100 μg/mL ampicillin and 34 μg/mL chloramphenicol). After *attP*-­-*attB* deletion in the cells containing pFB16+pFB10, pFB16+pFB13 or pFB16+pFB15, the mKate2 gene was deleted and no red color cells were detected under bright field (left) and imager (right).

**Supplementary Figure 3.**
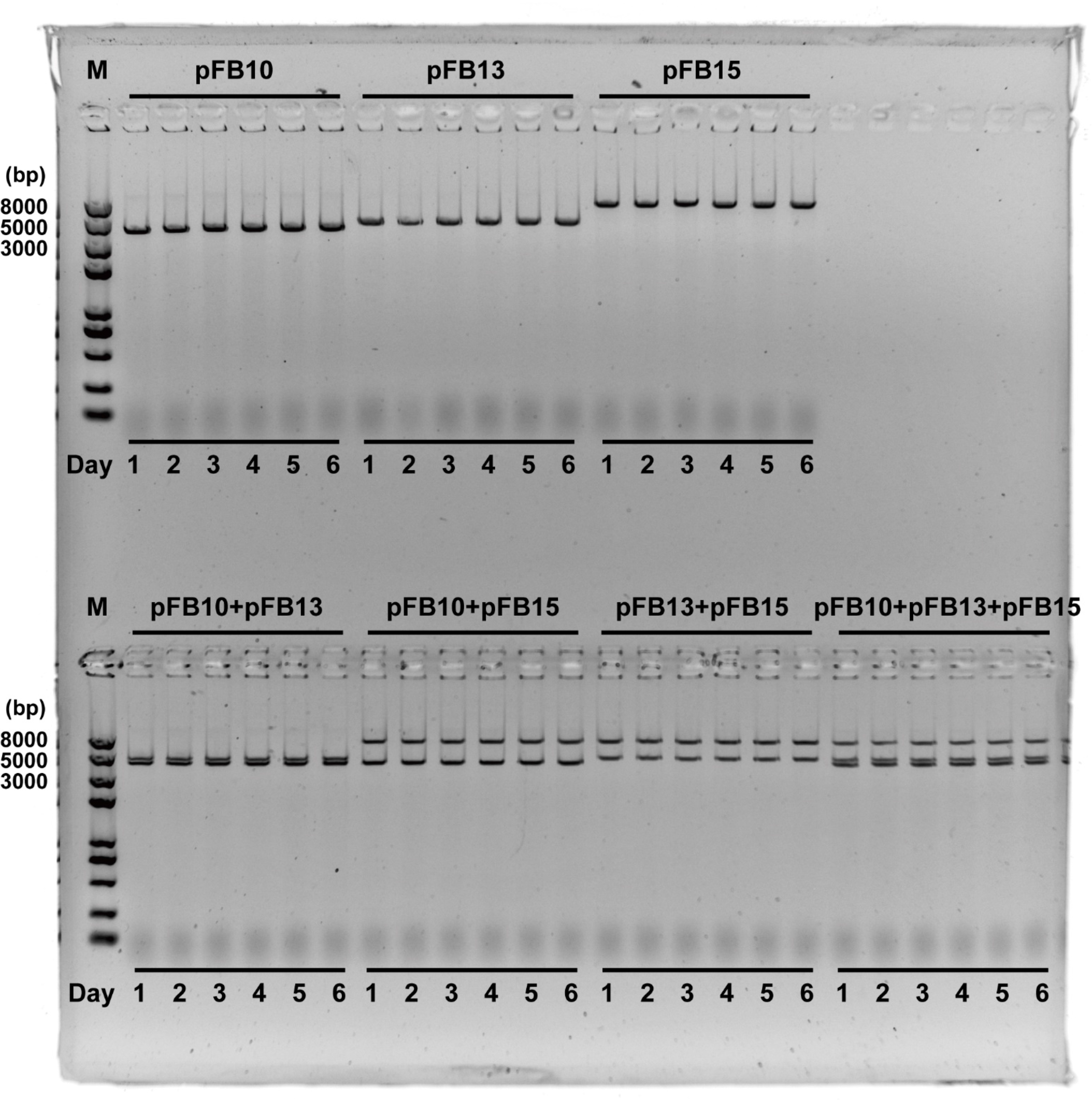
Stability of constitutive serine integrases expression plasmids. Image of gels post electrophoresis showing the genetic stability of low level constitutive serine integrases expression plasmids under single, double, and triple co-­-existing conditions. pFB10 (J23109-­-B0034-­-TP901-­-1(6xHis)-­-B0015, p15A/AmpR), pFB13 (J23109-­-B0034-­-Bxb1(6xHis)-­-B0015, pSC101, KanR), and pFB15 (J23109-­-B0034-­-Phic31(6xHis)-­-B0015, incW, SmR) belong to different compatibility groups and are able to co-­-exist in the same *E. coli* strain. Each *E. coli* strain characterization process was the same as described in **Supplementary Figure 1** and lasted for 6 days. All expected bands were detected (4007 bp of pFB10, 4490 bp of pFB13, and 7244 bp of pFB15) without mutation.

**Supplementary Figure 4.**
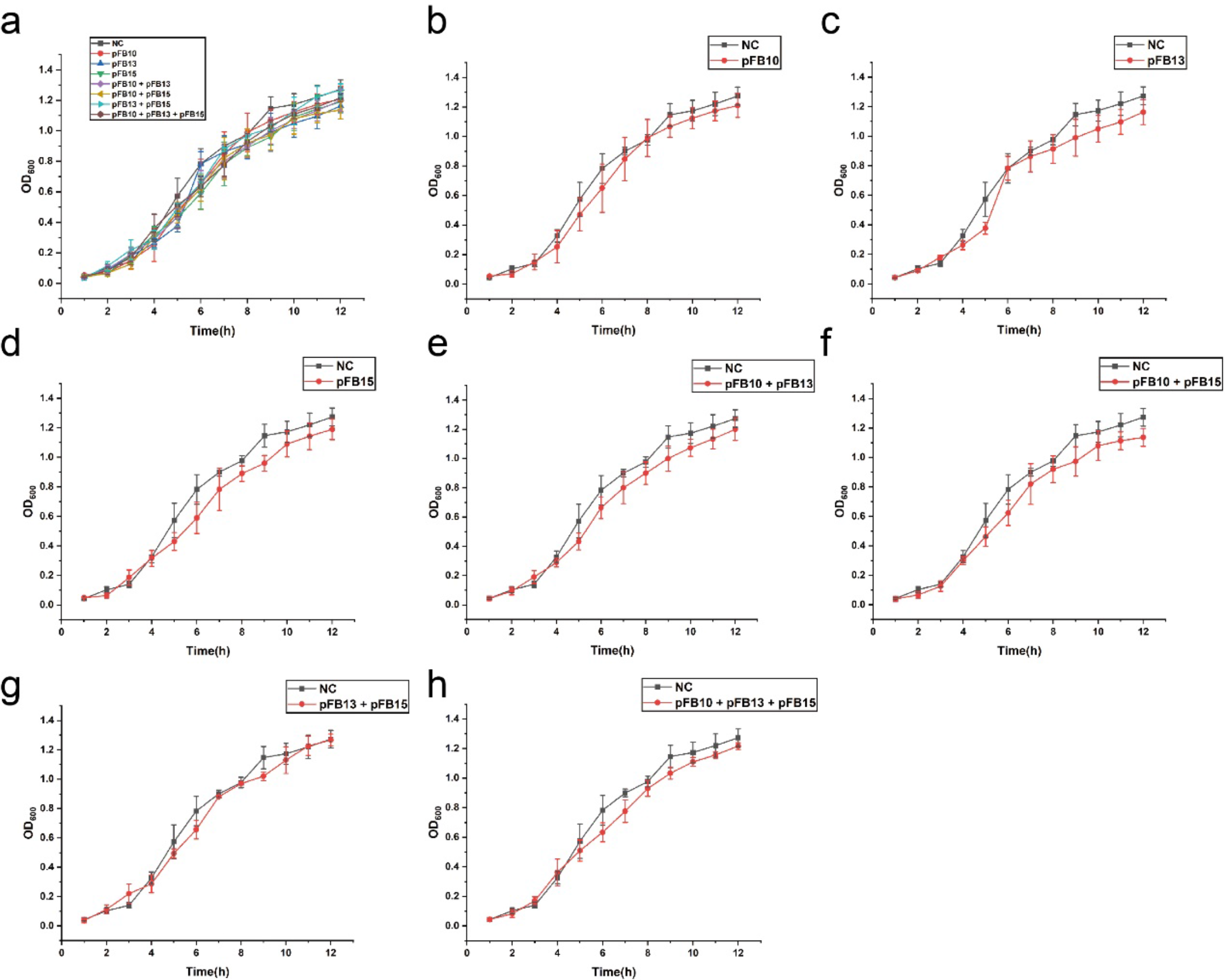
Constitutive low expression of serine integrases has no toxicity on *E. coli* growth. (**a**) Merged growth curves of *E. coli* strains containing different plasmids showing almost no toxicity on cell growth. All independent curves are shown from (**b**) to (**h**). Each “OD_600_ -­- Time” point was collected as three biological replicates.

**Supplementary Figure 5.**
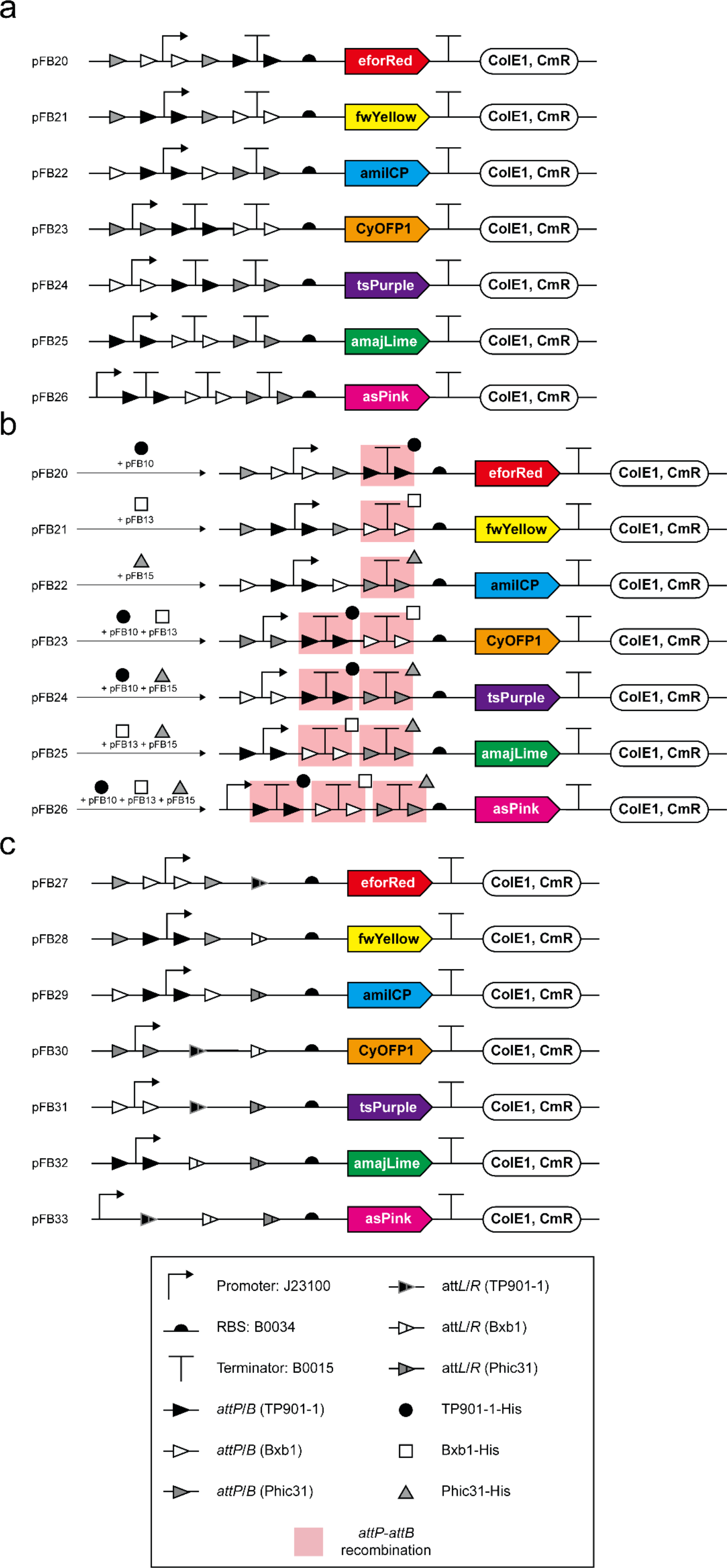
Seven “Locks” manipulated by three orthogonal “Keys”. (**a**) Detailed genetic circuit design of seven “Locks” circuits (pFB20 to pFB26). Terminator(s) between promoter and RBS blocked transcription (“Locked” state) and chromoproteins could not be expressed. (**b**) After plasmids co-­- transformation, the terminator(s) between promoter and RBS would be deleted by three orthogonal “Keys” (TP901-­-1, Bxb1, and Phic31), which represents a “Keys match Locks” process. (**c**) After deletion, there is no terminator(s) between promoter and RBS (pFB27 to pFB33), which leads to chromoproteins expression (“Unlocked” state).

**Supplementary Figure 6.**
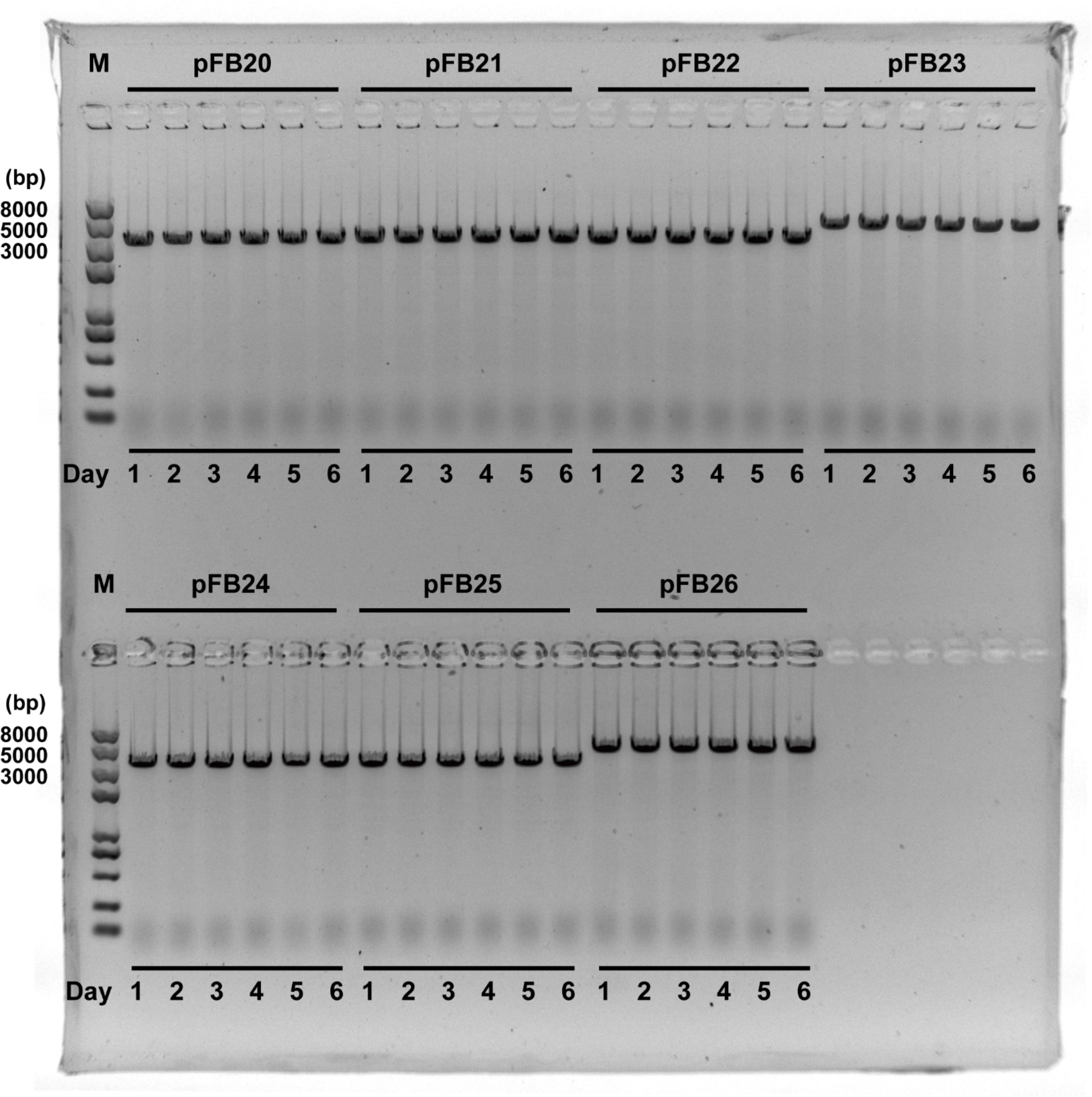
Stability of seven “Locks” plasmids. Image of gels post electrophoresis showing the genetic stability of seven “Locks” plasmids (pFB20 to pFB26). Each *E. coli* strain characterization process was the same as described in **Supplementary Figure 1** and lasted for 6 days. All expected bands were detected (3371 bp of pFB20, 3401 bp of pFB21, 3356 bp of pFB22, 3521 bp of pFB23, 3506 bp of pFB24, 3509 bp of pFB25, and 3647 bp of pFB26) without mutation.

**Supplementary Figure 7.**
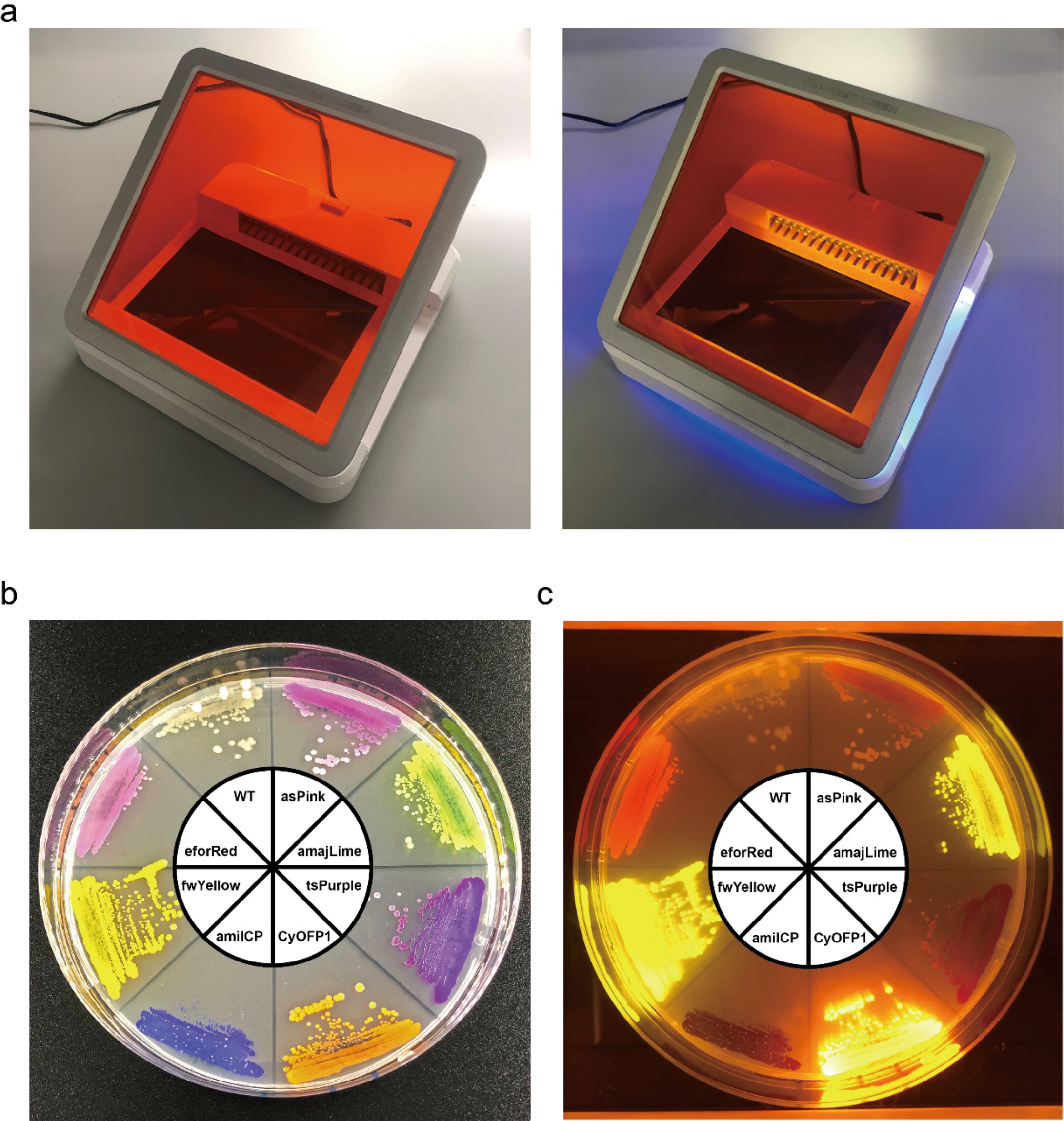
LED Transilluminator and chromoproteins used in this work. (**a**) LED Transilluminator (SLB-­-01W UltraSlim, MaestroGen) used in this work. Switch-­-off (left) and switch-­-on (right) are shown, respectively. (**b**) Chromoproteins used in this work. Bright field picture of LB-­-agar plate (34 μg/mL chloramphenicol) was incubated at 37°C for 24 h. Anticlockwise starting from 12 o’clock: wild type (WT, empty pSB1C3), eforRed (pFB131), fwYellow (pFB132), amilCP (pFB133), CyOFP1 (pFB134), tsPurple (pFB135), amajLime (pFB136), and asPink (pFB137). (**c**) Picture of the same LB-­-agar plate under LED Transilluminator. Fluorescence of eforRed, fwYellow, CyOFP1, and amajLime can be clearly observed.

**Supplementary Figure 8.**
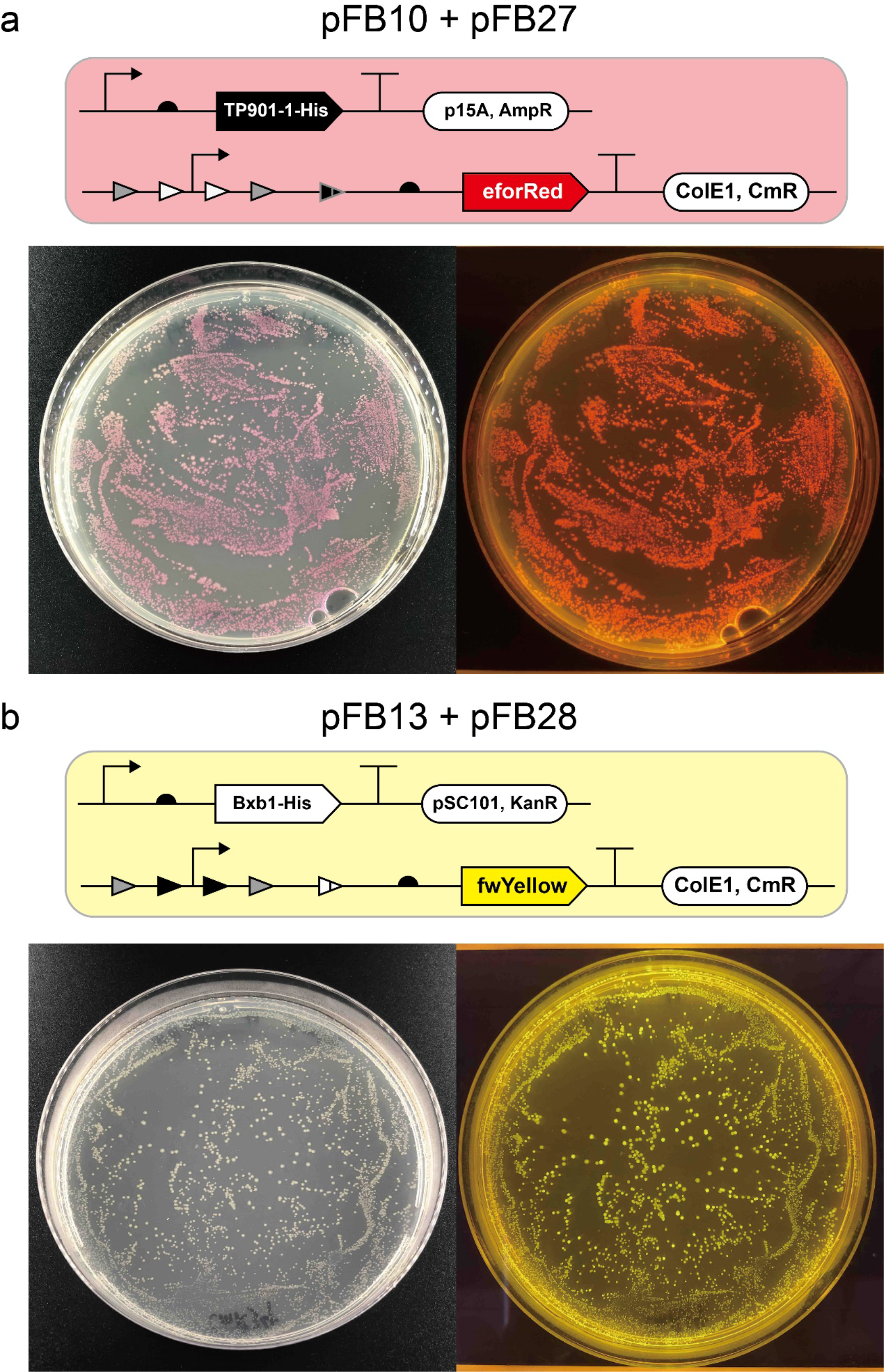
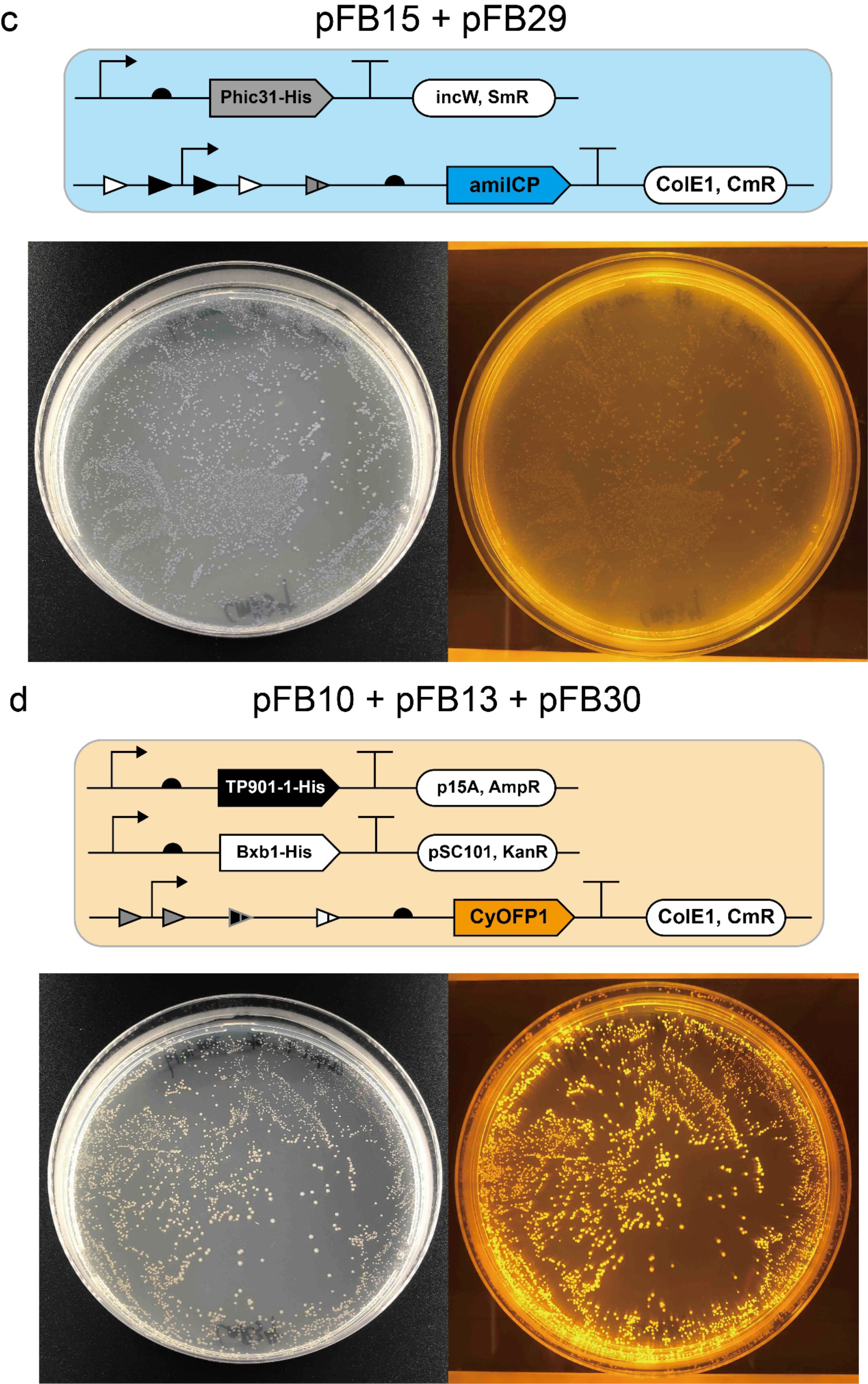
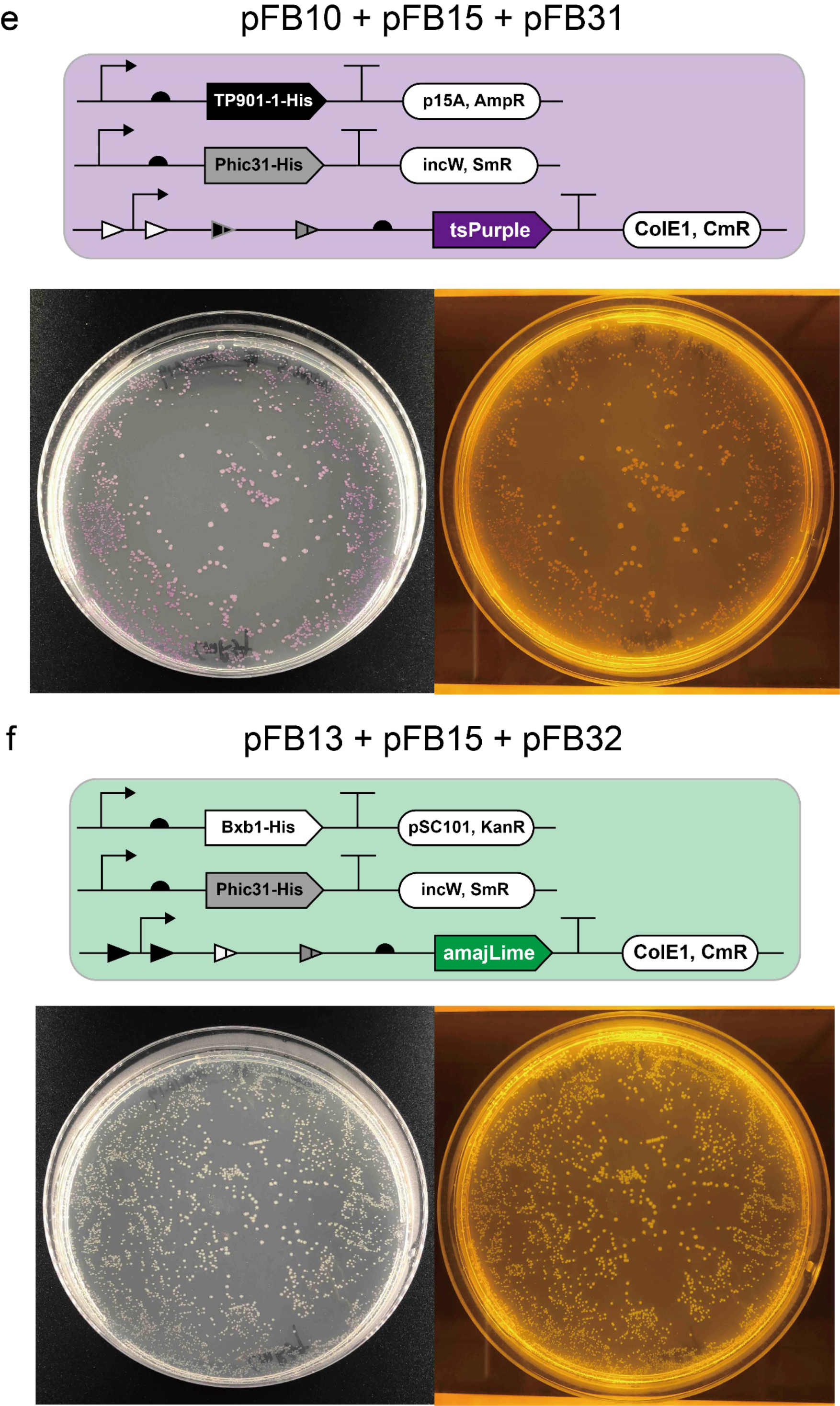
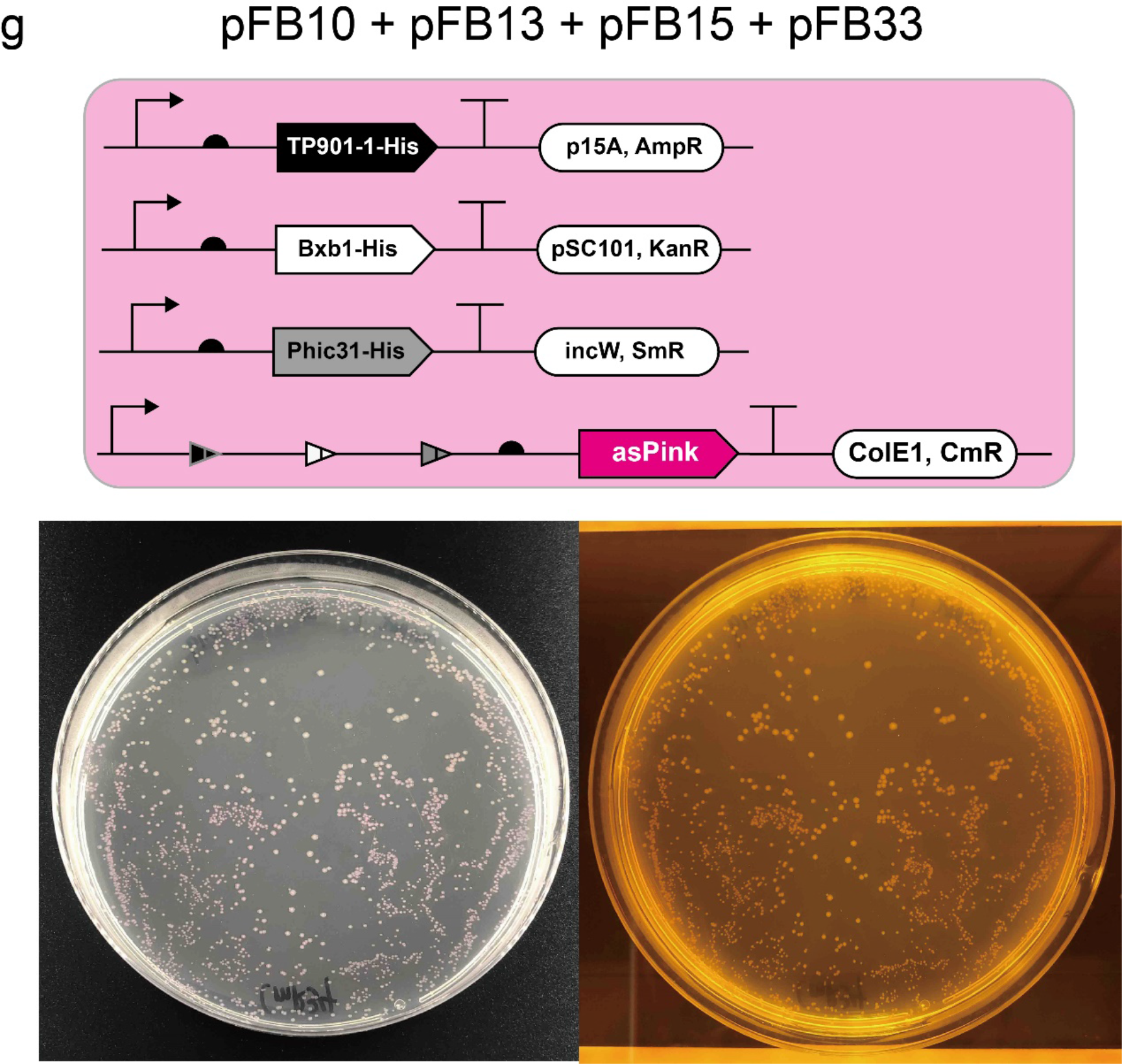
Recombination efficiency characterization. (**a**)-­-(**g**) Schematic of “Keys” and “Locks” plasmid combinations in *E. coli* (top panel) and the same LB-­-agar plate (bottom panel) under bright field (left) and LED Transilluminator (right). After co-­-transformation, each *E. coli* strain displayed corresponding colors under bright field or LED Transilluminator. Otherwise, negative *E. coli* colonies on the plate should be wild type without specific colors. The various *E. coli* color features are shown in **Supplementary Figure 7**.

**Supplementary Figure 9.**
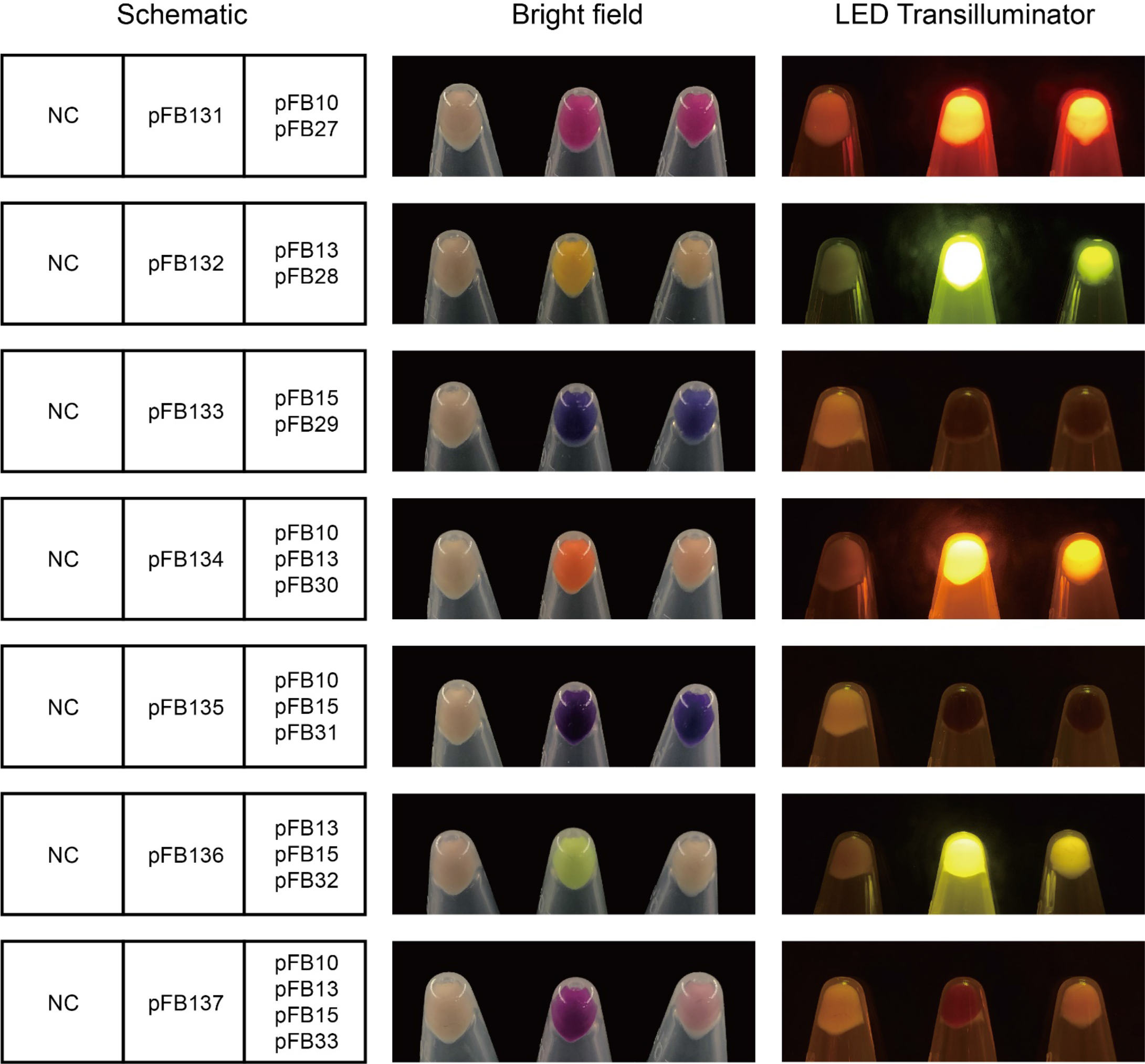
Characterization of seven “Unlocked” plasmids. Each line represents one “Unlocked” plasmid characterization. Each box (left) consists of three different *E. coli* strains: one negative control (NC, vector pSB1C3), one positive control (pFB131 to pFB137), and one experimental group. The chromoprotein’s expression level of all seven experimental groups were lower as compared to their respective positive control. However, the experimental groups’ colors could always be observed. Each 1.5 mL centrifuge tube contains *E. coli* pellets collected from 3 mL overnight culture (30°C, 250 rpm, 18 h).

**Supplementary Figure 10.**
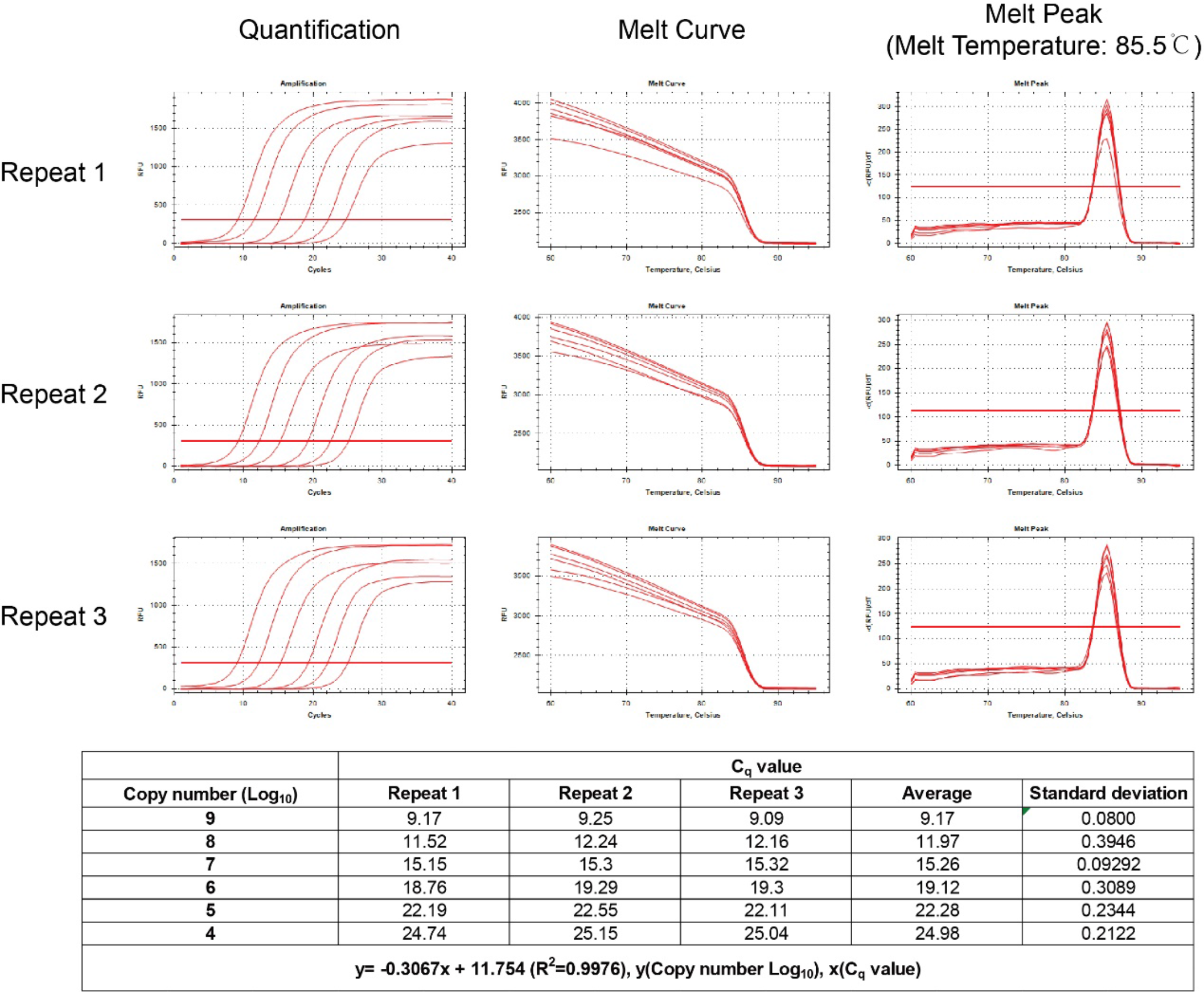
qPCR standard curve calculation of *E. coli* 16s rRNA. pFB145 acts as a template for *E. coli* 16s rRNA qPCR standard curve calculation (see **Supplementary Table 6** for the primer pair 357F/518R). This experiment was repeated with three biological replicates.

**Supplementary Figure 11.**
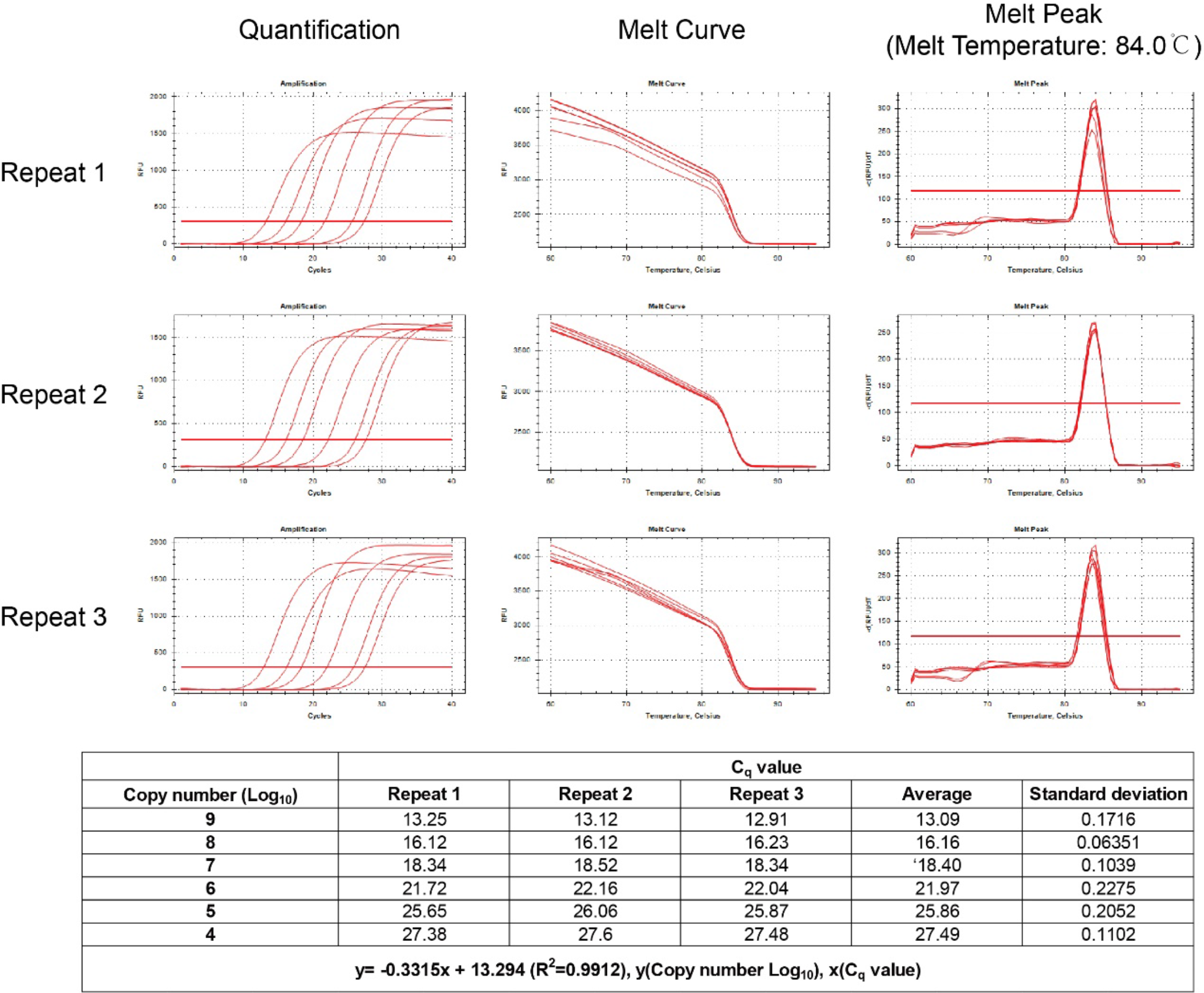
qPCR standard curve calculation of sfGFP mRNA. pFB141 acts as a template for sfGFP mRNA qPCR standard curve calculation (see **Supplementary Table 6** for the primer pair sfGFP-­-F-­-3/sfGFP-­-R-­-3). This experiment was repeated with three biological replicates.

**Supplementary Figure 12.**
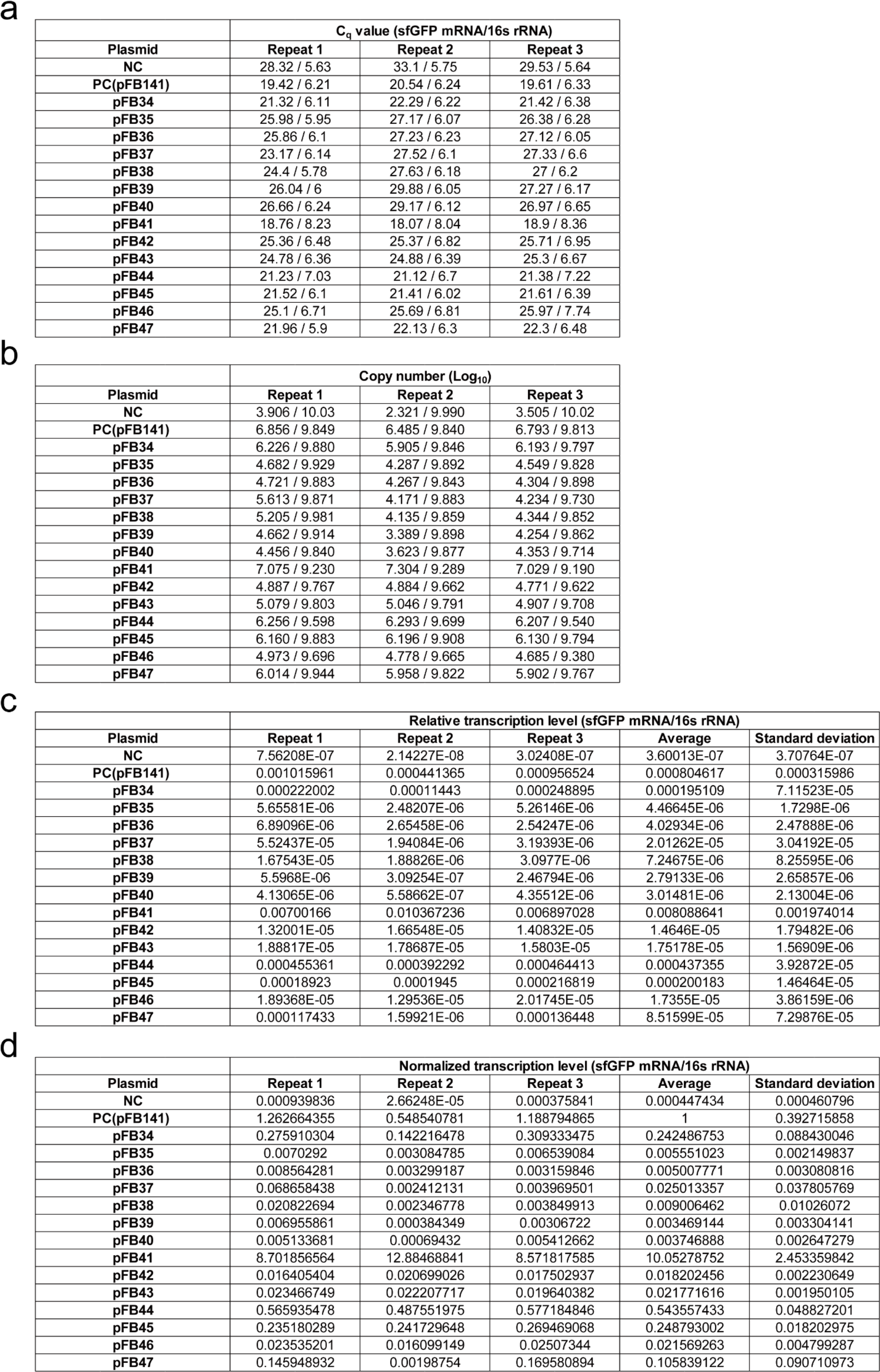
qPCR result of seven “Locks” plasmids quantification. (**a**) qPCR result of seven “Locks” plasmids before (pFB34 to pFB40) and after (pFB41 to pFB47) TP901-­-1/Bxb1/Phic31 mediated *attP*-­-*attB* deletion. Negative control (pSB1C3) and positive control (pFB141) were also tested. All cDNA samples were prepared after *E. coli* total RNA extraction and mRNA to cDNA reverse transcription. (**b**) cDNA copy number was calculated based on standard curves (**Supplementary Figures 12 and 13**). (**c**) and (**d**) were relative and normalized transcriptional level of each sample’s sfGFP mRNA/16s rRNA ratio. The value of positive control was considered as “1” for normalization.

**Supplementary Figure 13.**
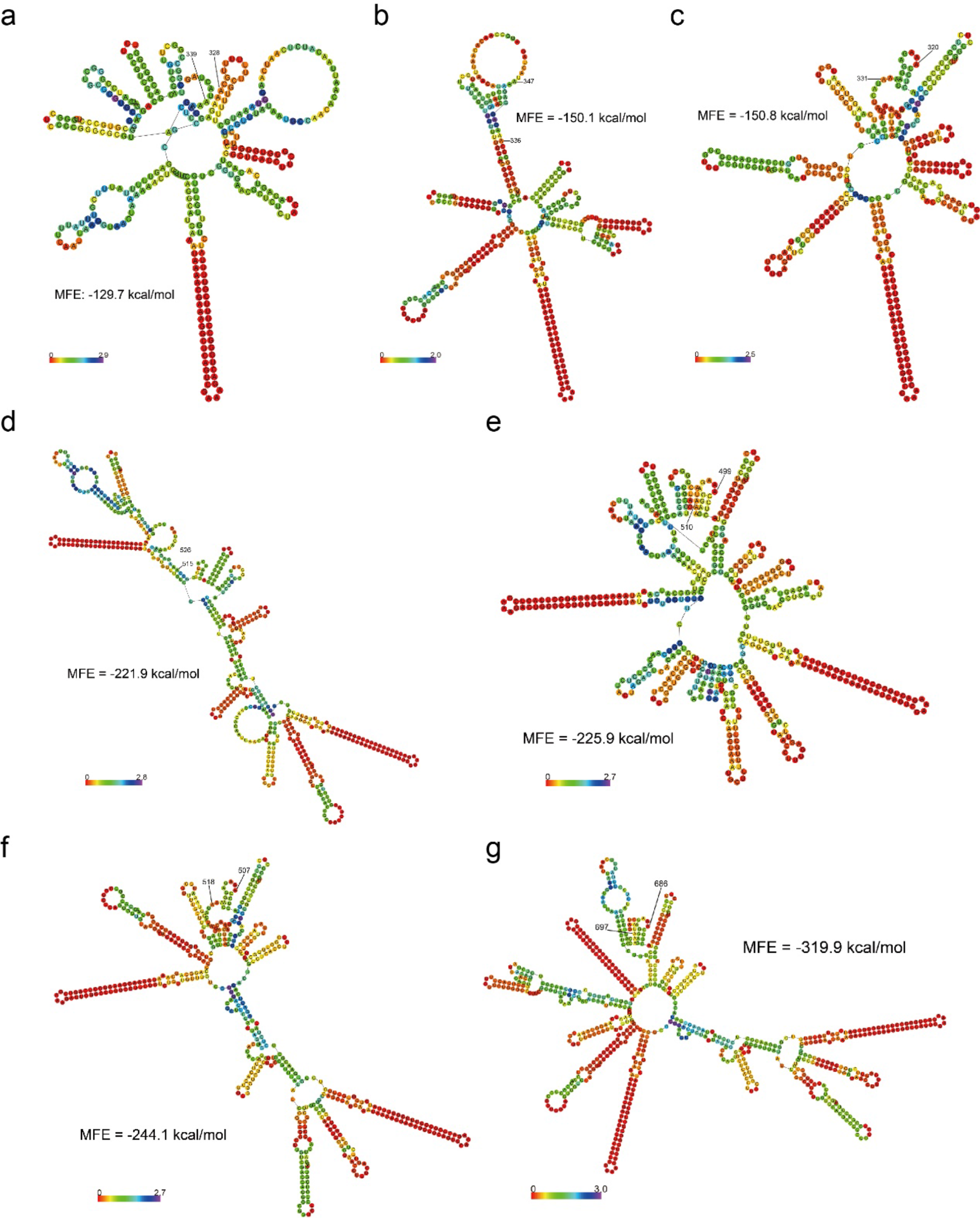
Predicted Minimum Free Energy (MFE) secondary structure of seven “Locked” 5’-­-UTRs before *attP*-­-*attB* deletion. The seven mRNA sequences are from the +1 site to the last base before start codon (ATG). All mRNA structures are annotated with predicted Minimum Free Energy (MFE) and colored positional entropy. The position of RBS BBa_B0034 (aaagaggagaaa) is labelled with numbers. (**a**)-­-(**g**) correspond to pFB34-­-pFB40. All structures are predicted using the “***RNAfold*** WebServer”.

**Supplementary Figure 14.**
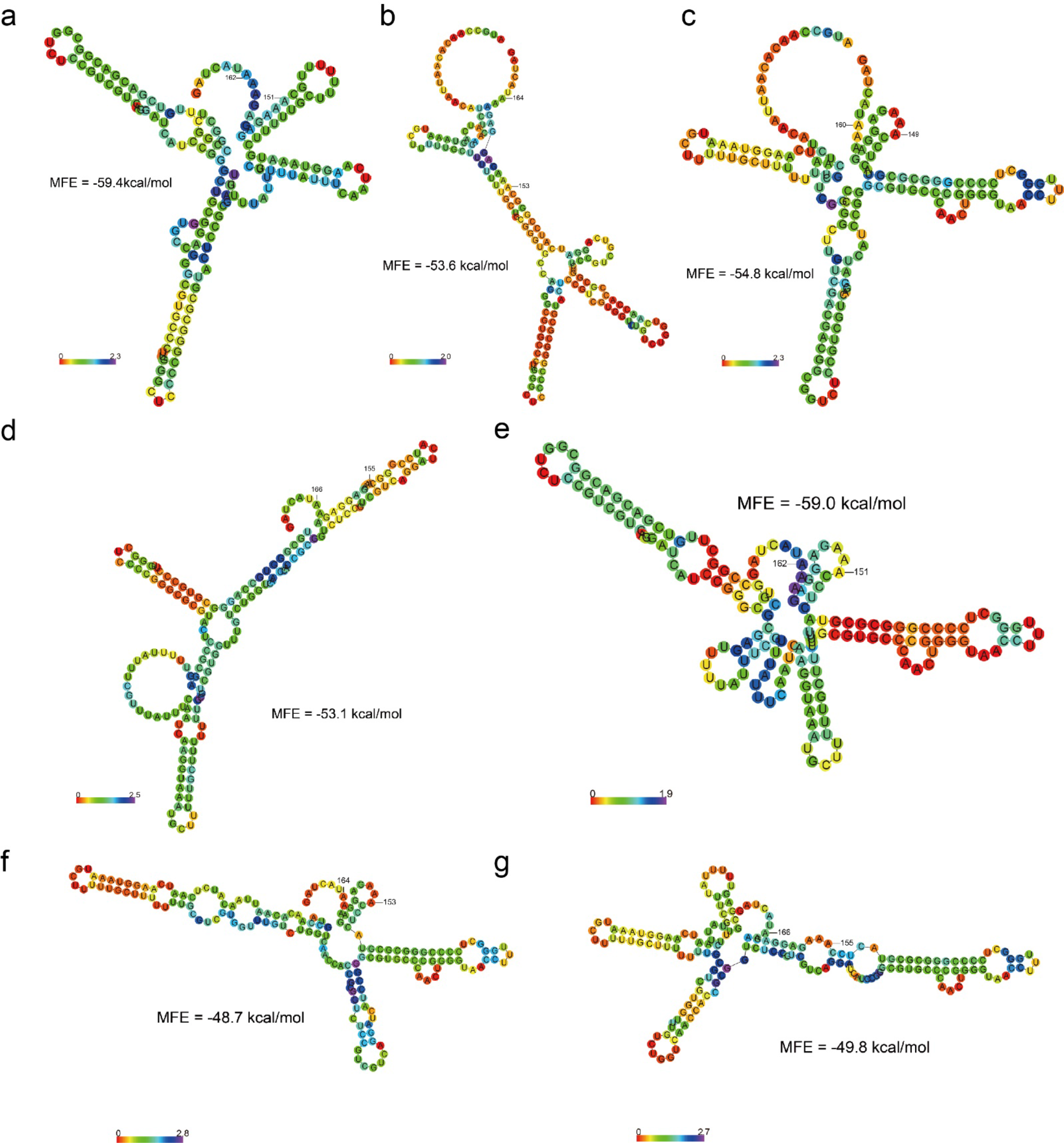
Predicted Minimum Free Energy (MFE) secondary structure of seven “Unlocked” 5’-­- UTRs after *attP*-­-*attB* deletion. The seven mRNA sequences are from the +1 site to the last base before start codon (ATG). All mRNA structures are annotated with predicted Minimum Free Energy (MFE) and colored positional entropy. The position of RBS BBa_B0034 (aaagaggagaaa) is labelled with numbers. (**a**)-­-(**g**) correspond to pFB41-­-pFB47. All structures are predicted using the “***RNAfold*** WebServer”.

**Supplementary Figure 15.**
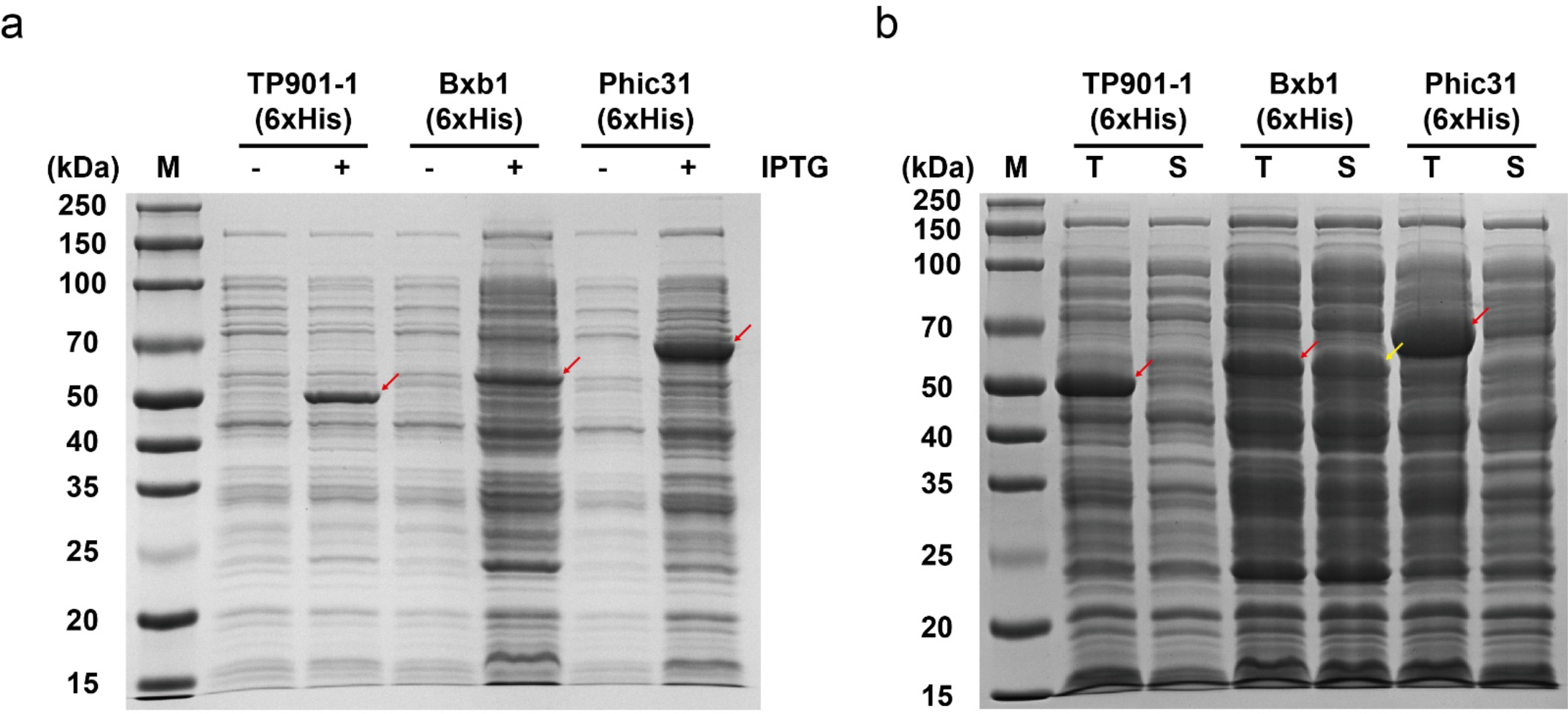
SDS-­-PAGE analysis of TP901-­-1(6xHis)/Bxb1(6xHis)/Phic31(6xHis) purification. When pFB142/pFB143/pFB144 contained *E. coli* Rosetta-­-gami(DE3) pLysS strains (inoculated in LB medium, 1 L conical flask) reached OD_600_ = 0.6-­-0.8, 1 mL 1 M IPTG were added and then cultivation for another 6 h at 30°C and 250 rpm. (a) SDS-­-PAGE of cell lysates showing a strong total expression of three serine integrases after induction (red arrows). (b) The total crude extract after sonication (T) and supernatant after centrifugation (S) analysis showing that only Bxb1(6xHis) was soluble (yellow arrow; TP901-­-1(6xHis) and Phic31(6xHis) were almost insoluble in this situation.

**Supplementary Figure 16.**
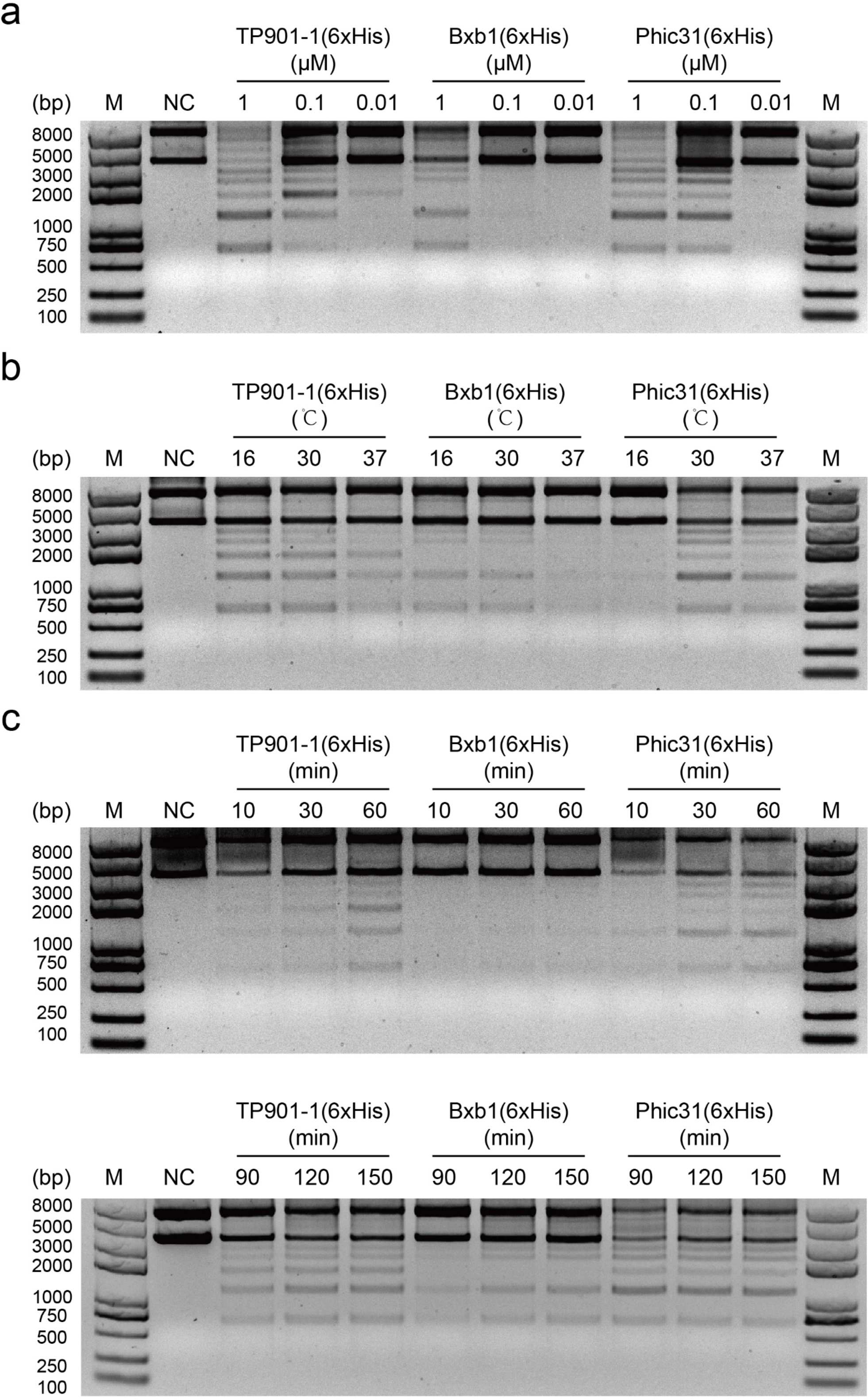
Optimization of *in vitro* SYMBIOSIS conditions. All tests were based on pFB16 *attP*-­-*attB* deletion and nonspecific fragments integration. The fragments were composed of three types of linear DNA: After deletion, pFB16 can be digested into two kinds of strands: the “*attL*-­- (*attP*s)-­-J23100-­-B0034-­-mKate2-­-B0015-­-(*attB*s)-­-*attR*” fragments and the “*attL*-­-pSB1C3-­-*attR*” vectors. Besides “fragments” and “vectors”, the third type of linear DNA was a composite of heterologous pFB16 polymers ((pFB16)_n_, a heterologous *attP*-­-*attB* recombination among different pFB16 ones). (**a**) Optimization of serine integrase concentration. All samples were incubated at 30°C for 12 h. 1 μM of all three enzymes showed the best catalytic efficiency. (**b**) Optimization of reaction temperature. All samples were incubated for 1.5 h with 1 μM of enzymes. The common preferred temperature for TP901-­-1(6xHis)/Bxb1(6xHis)/Phic31(6xHis) was 30°C. (**c**) Optimization of reaction time. All samples were incubated at 30°C with 1 μM of enzymes. The optimal reaction time was 120 min for all three enzymes.

**Supplementary Figure 17.**
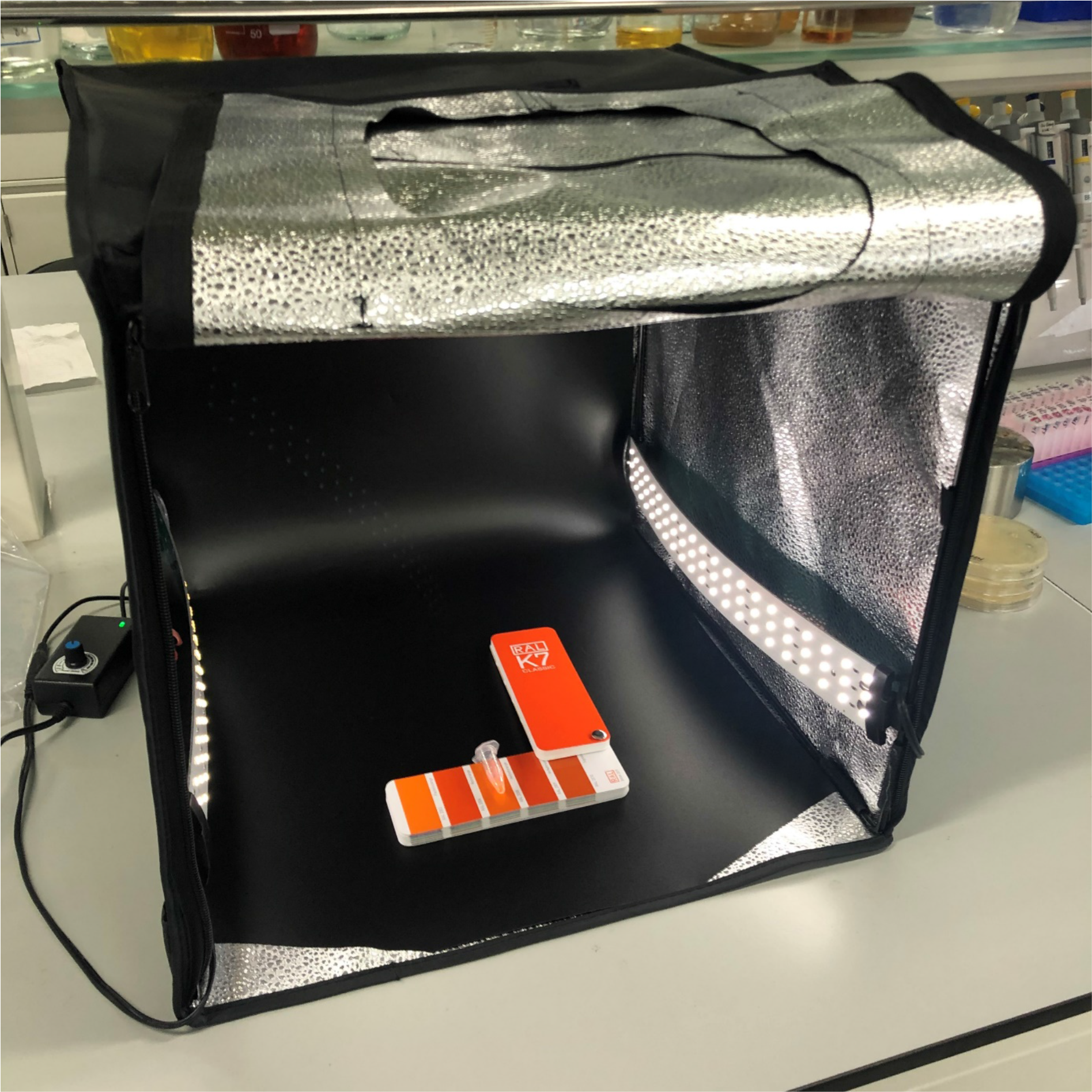
The mini photo studio for colorimetry and taking bright field images. Image showing the mini photo studio, RAL color chart, and colorimetry. All bright field images used in this study were taken here by a mobile phone without post-­-processing.

**Supplementary Figure 18.**
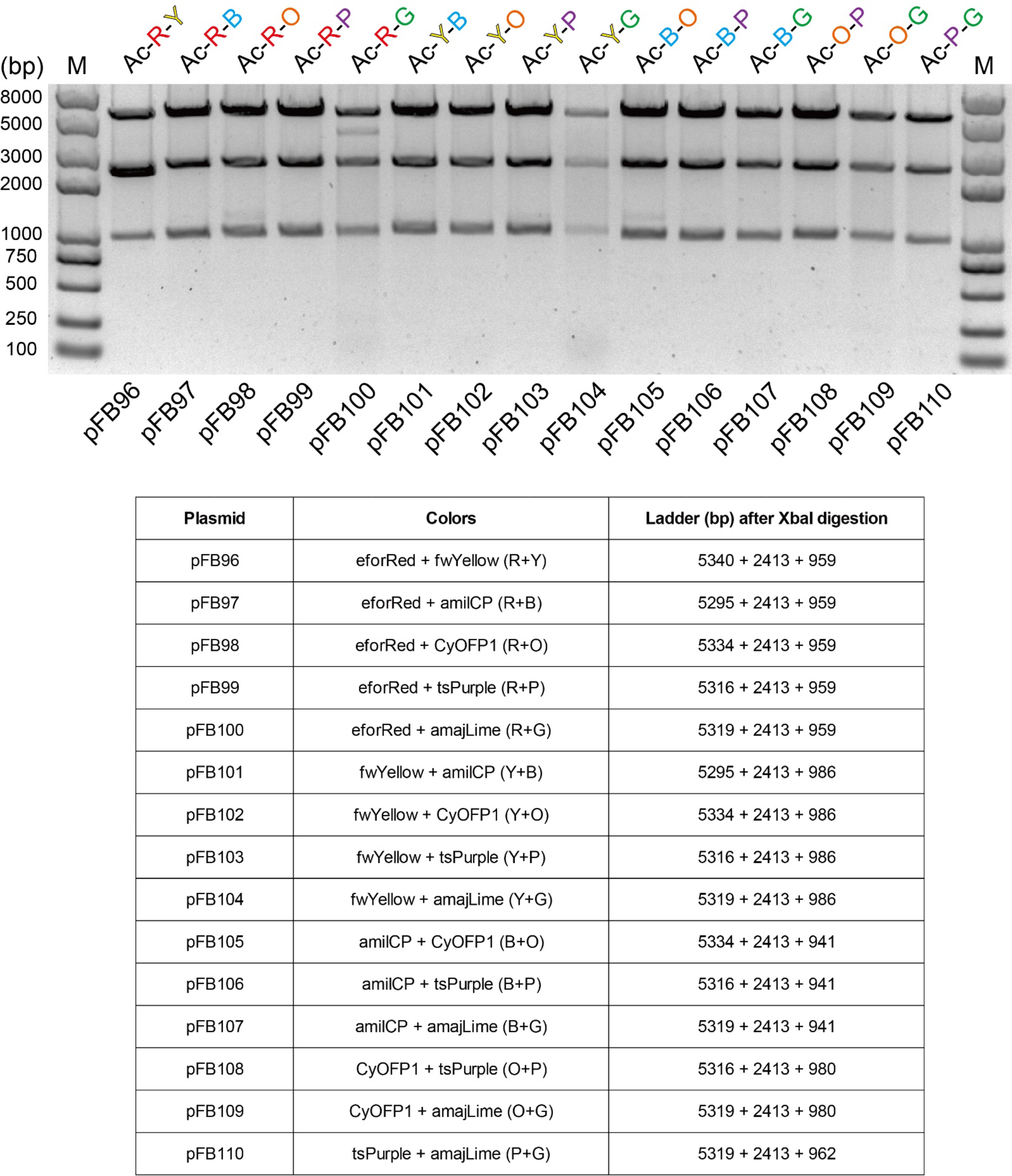
XbaI digestion of “Acceptor-­-double Donors” assembled plasmids. All 15 “Acceptor-­-double Donors” assembled plasmids were separated by agarose gel electrophoresis after FastDigest XbaI (Thermo Scientific) digestion.

**Supplementary Figure 19.**
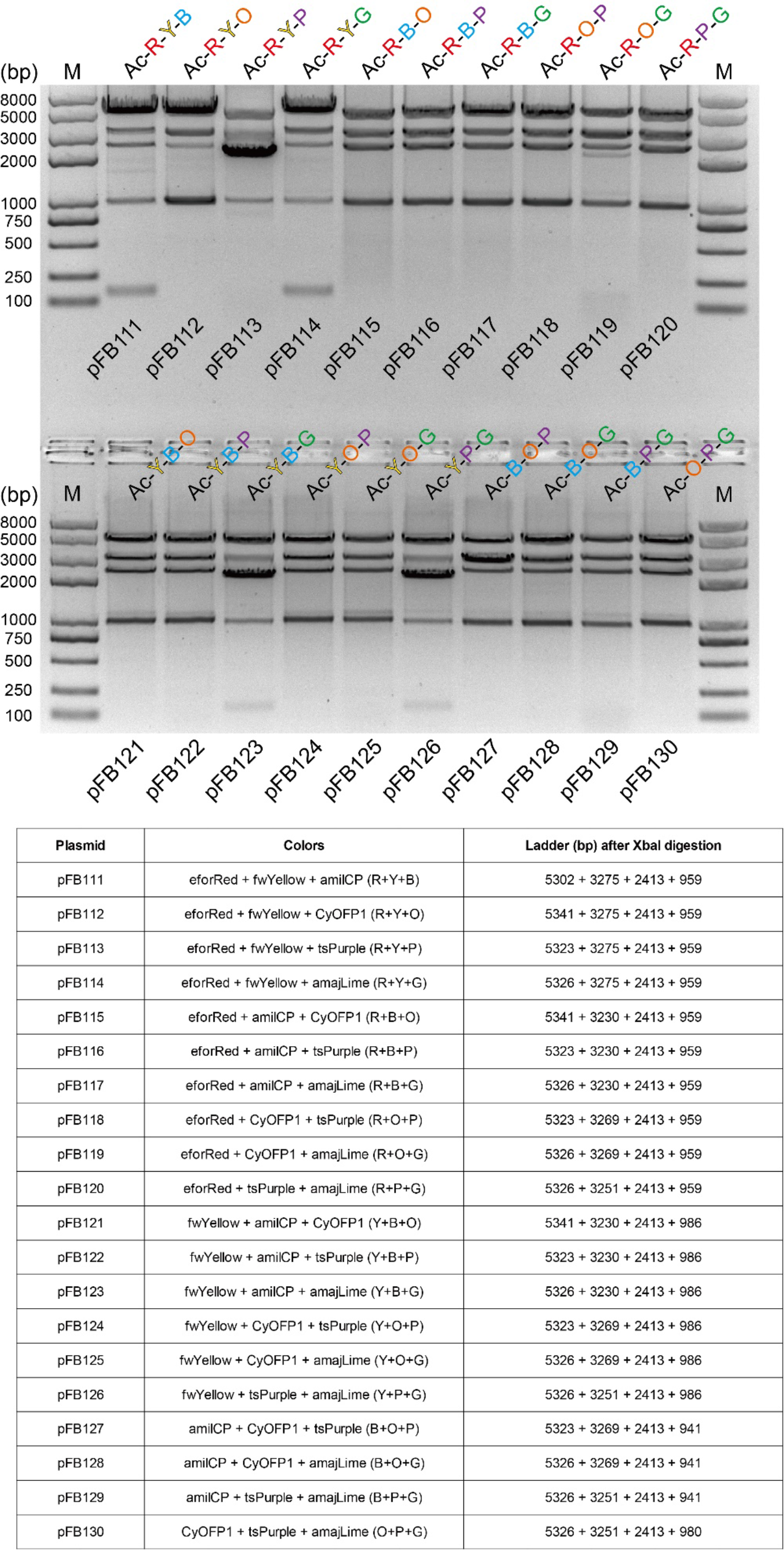
XbaI digestion of “Acceptor-­-triple Donors” assembled plasmids. All 20 “Acceptor-­-triple Donors” assembled plasmids were separated by agarose gel electrophoresis after FastDigest XbaI (Thermo Scientific) digestion.

**Supplementary Figure 20.**
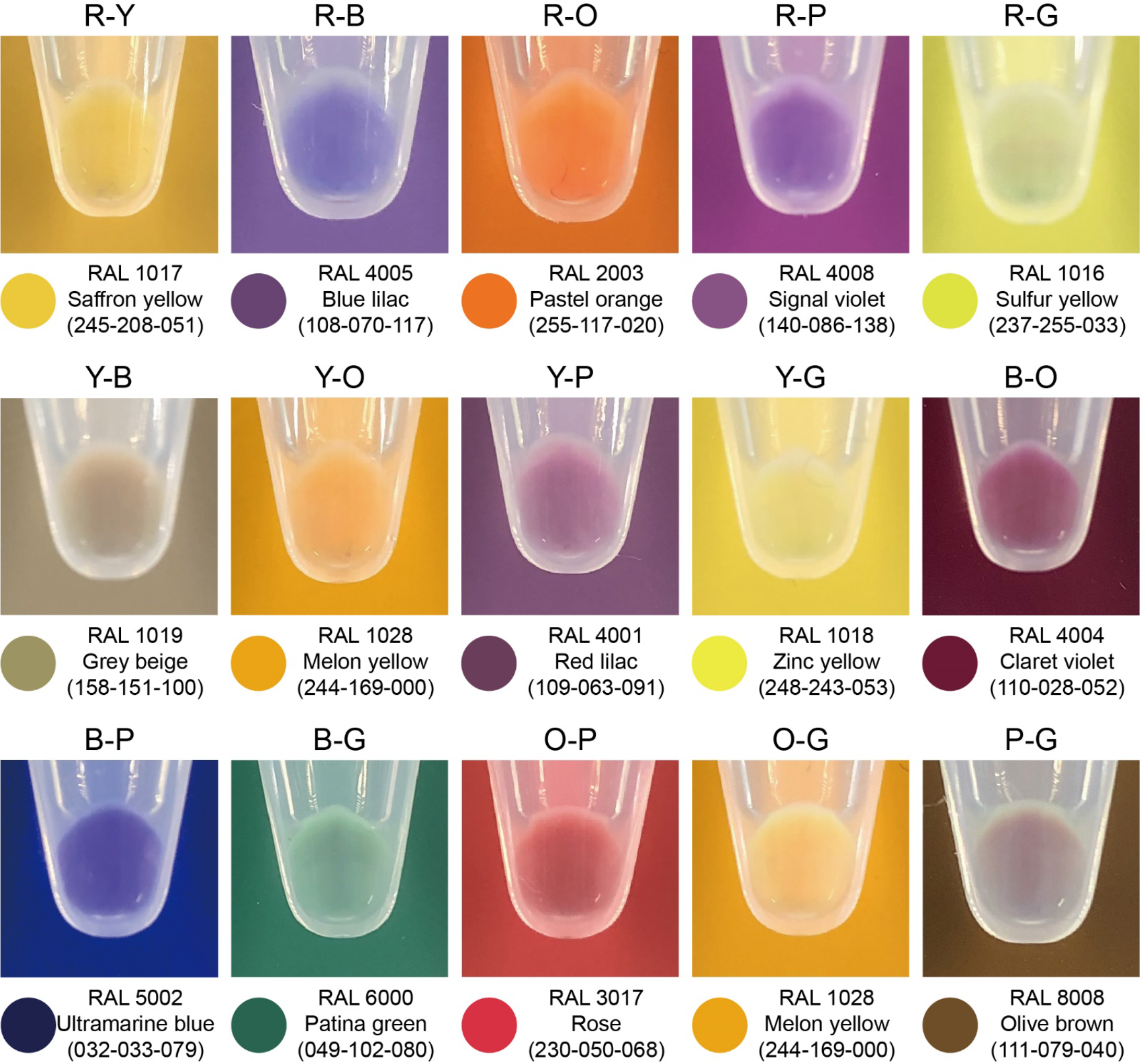
Composite colors created by “Acceptor-­-double Donors” assembled plasmids. All 15 composite colors were created by different chromoproteins and observed within *E. coli* Mach1-­-T1 pellets. The 15 assembled plasmids were pFB96 to pFB110, respectively.

**Supplementary Figure 21.**
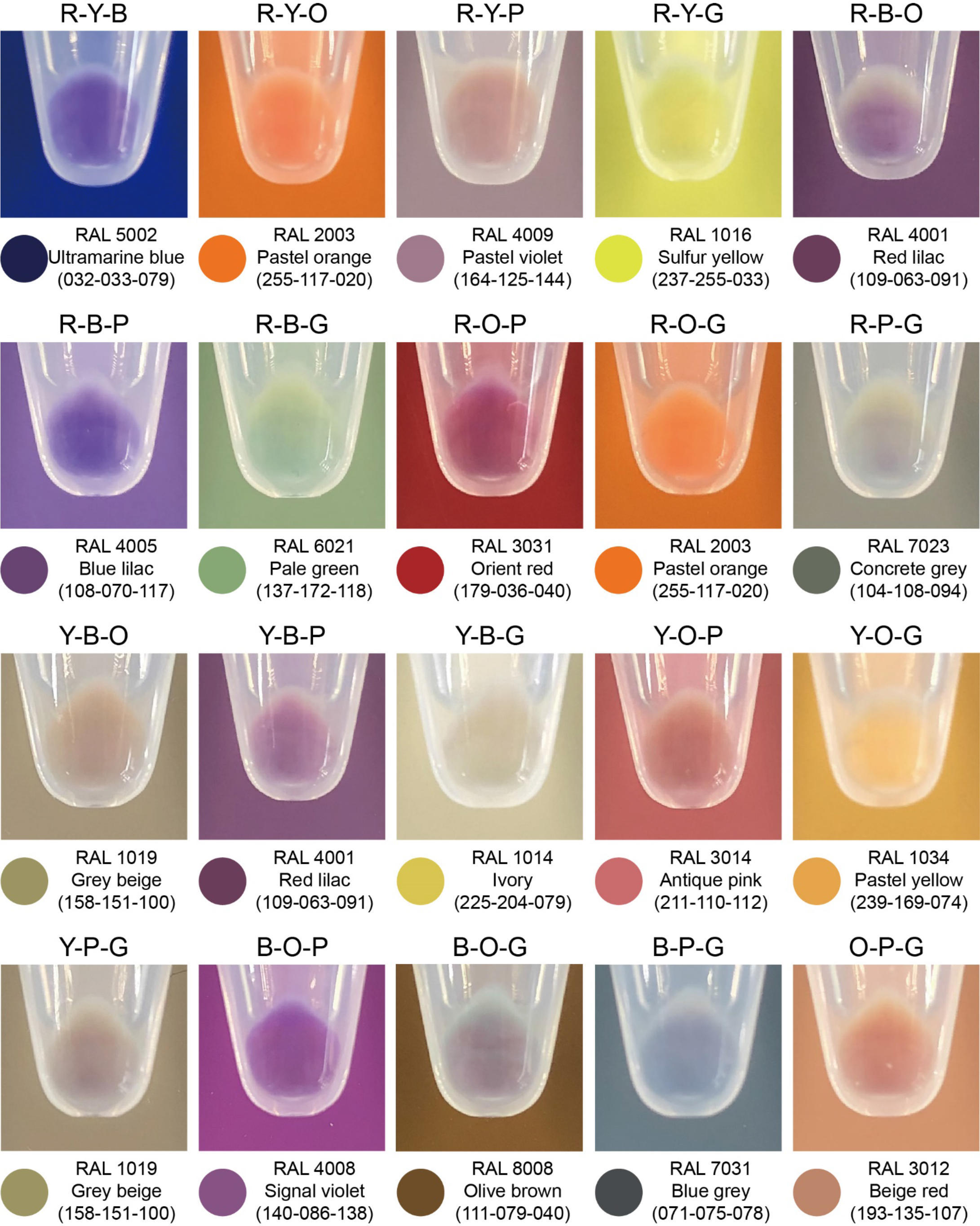
Composite colors created by “Acceptor-­-triple Donors” assembled plasmids. All 20 composite colors were created by different chromoproteins and observed within *E. coli* Mach1-­-T1 pellets. The 20 assembled plasmids were pFB111 to pFB130, respectively.

